# Measurement reliability bounds functional benchmarks and relocates where variant effect prediction fails

**DOI:** 10.64898/2026.09.14.751496

**Authors:** Ningyi Zhang

## Abstract

**Background:** Variant effect predictors are increasingly benchmarked against multiplexed assays of variant effect (MAVEs) rather than clinical labels, which removes label circularity but introduces a new problem: a correlation against a measurement cannot exceed the measurement’s own reproducibility, and precision varies sharply across the territories compared.

**Results:** We scored nineteen predictors across sixteen strata of a frozen atlas of 64,178 saturation genome editing variants in seven cancer-susceptibility genes. From published replicate scores and standard errors we estimated each territory’s reliability ceiling, and showed by simulation that the correction reduces error above a ceiling of about 0.45 and amplifies it below. Ceilings vary more across territory than predictors do, and correcting for them redraws the map at the splice extremes. The collapse at canonical splice sites is largely a property of the assay: the median shortfall relative to coding narrows from 1.7-to 1.4-fold; this convergence survives dropping *BARD1* or *PALB2* but inverts when *BRCA1* is dropped, so we report all three leave-one-gene-out folds rather than claim gene independence, and the frontier parity rests on one deposit. Genuine failure lies 11–50 bp into the intron, which the uncorrected map presents as modest. Across MaveDB, 2,452 of 2,803 score sets carry, at the upper bound, what a reliability estimate needs, though a conventional column-name search finds only a tenth; among 674 human deposits with a computable ceiling, 29.9–51.8% fall below 0.90. Scored as classification against the assays’ own functional calls in three genes, the same predictors separate damaging from tolerated better than their correlations suggest, though none reaches the strongest evidence band at the 95%-specificity operating point.

**Conclusions:** Territory-resolved benchmarks should report a per-stratum reliability estimate, or state that the assay permits none. It asks nothing of depositors and applies today, at the upper bound, to most (87%) of MaveDB.

## Background

Interpreting human genetic variation remains a rate-limiting step of genomic medicine, and computational variant effect predictors fill the gap for the variants of uncertain significance that clinical observation cannot resolve. Successive generations of predictor — integrative supervised scores [1], conservation metrics [2, 3], dedicated splice predictors [4, 5], protein-centric deep models [6], the meta-predictors that dominate clinical practice [7, 8, 9, 10, 11], and genomic foundation models trained by self-supervision on sequence alone [12, 13, 14, 15] — are now routinely compared. How that comparison should be conducted is itself an open problem.

Correlation against ClinVar labels [16] suffers from circularity, because many predictors were trained on or thresholded against those labels; from ascertainment bias, because recorded variants are a clinically selected subset; and from territory blindness, because pooled evaluation mixes coding, splice-proximal and deep-intronic variants whose mechanisms and predictability differ. Multiplexed assays of variant effect (MAVEs), and saturation genome editing (SGE) in particular [17, 18, 19], offer an alternative ground truth: dense, experimentally measured functional scores with no dependence on clinical annotation, now collected at scale in MaveDB [20, 21]. Benchmarks built on MAVEs have reshaped predictor evaluation for protein-coding variants [22, 23, 24] and, more recently, for splicing [25]; what a predictor release should publish is now codified as well [26].

Substituting an experimental measurement for a clinical label introduces a difficulty that label-based benchmarks never had to face. A rank correlation against a noisy measurement is bounded above by the square root of that measurement’s reliability [27, 28], and prior work has recognised that assay reproducibility limits what a benchmark can observe. Livesey and Marsh [22, 23] assessed the reproducibility of deep mutational scanning datasets directly, by correlating independent experiments on the same protein, as a check on their suitability as a benchmark. What has not been done is to estimate that bound per functional territory, correct the reported performance for it, and establish the range over which such a correction is trustworthy.

This matters because an assay’s precision is not constant across the variants it measures: read depth, the dynamic range of the selection, and how close a variant class sits to the assay’s saturation point all vary by territory. A territory-resolved benchmark that reports only the achieved correlation therefore cannot separate a territory where predictors are weak from one where the assay is imprecise, and will attribute the latter to the former.

A related question is how sensitive a benchmark is to the scoring definition it pins, since predictors expose several scores, windows and aggregations; we treat that as a robustness check on the main result rather than as a separate problem.

Here we address that gap on a frozen functional standard of 64,178 saturation genome editing variants across seven cancer susceptibility genes. The claim we make is a general one about benchmarking against noisy measurements: the achievable correlation is bounded by assay reliability, that bound is estimable per territory from error models the assay authors already publish, the correction has a definable range of validity, and applying it relocates where predictors genuinely fail. The splice-site result is the demonstration rather than the thesis. Four further analyses support it: paired dependent-correlation tests in place of interval overlap, the same grid scored as classification and as American College of Medical Genetics and Genomics (ACMG)-style evidence strength [29, 30, 31, 32], a comparison against a second readout of the same assays, and a systematic sweep of one predictor’s definition space as a robustness check.

The atlas assembled here exists because territory-resolved benchmarking needs a functional standard that is public, uniformly processed, and auditable variant by variant. Saturation genome editing deposits satisfy the first condition but not the other two. Each is released against its own transcript, in its own orientation, under its own scoring convention, and none carries the variant-level region annotation a stratified comparison requires. Assembling one is therefore not a matter of concatenation, and the intermediate product — a frozen matrix in which every variant has a genomic coordinate, a region class, an oriented functional score and nineteen predictor scores — is reusable independently of the analyses reported here.

That matrix supports any benchmark against these seven assays without re-scoring, and the nineteen-predictor by sixteen-stratum grid it yields is released as a table in its own right. Because reliability estimates and attenuation ceilings are computed per stratum and published alongside, a reader can recompute any comparison reported here under a different correction, a different stratification, or none.

## Results

### A frozen functional-standard atlas of 64,178 SGE variants

We assembled seven saturation genome editing assays from MaveDB [20, 21] covering *BAP1*, *BARD1*, *BRCA1*, *BRCA2*, *PALB2*, *RAD51C* and *VHL* (Table 1), spanning 64,178 variants — 46,392 single-nucleotide variants (SNVs) and 17,786 indels — mapped to GRCh38 on MANE-Select transcripts via Mutalyzer 3 [33, 34] (version note in Methods), with per-assay score orientation harmonised so that larger values indicate more damaging function (Fig. 1A,B; Methods). Variants fall into five mutually exclusive regions: coding/UTR (n = 52,523), splice ±1–2 (n = 1,056), splice 3–10 bp (n = 3,955), splice 11–50 bp (n = 5,961) and deep intronic >50 bp (n = 683). All assay rows mapped successfully. Protein consequence was re-derived locally from the same transcript models, giving 26,022 missense, 7,886 synonymous and 1,656 nonsense SNVs; as an external check the derived missense set is identical, variant for variant, to the set AlphaMissense scores.

**Figure 1.**
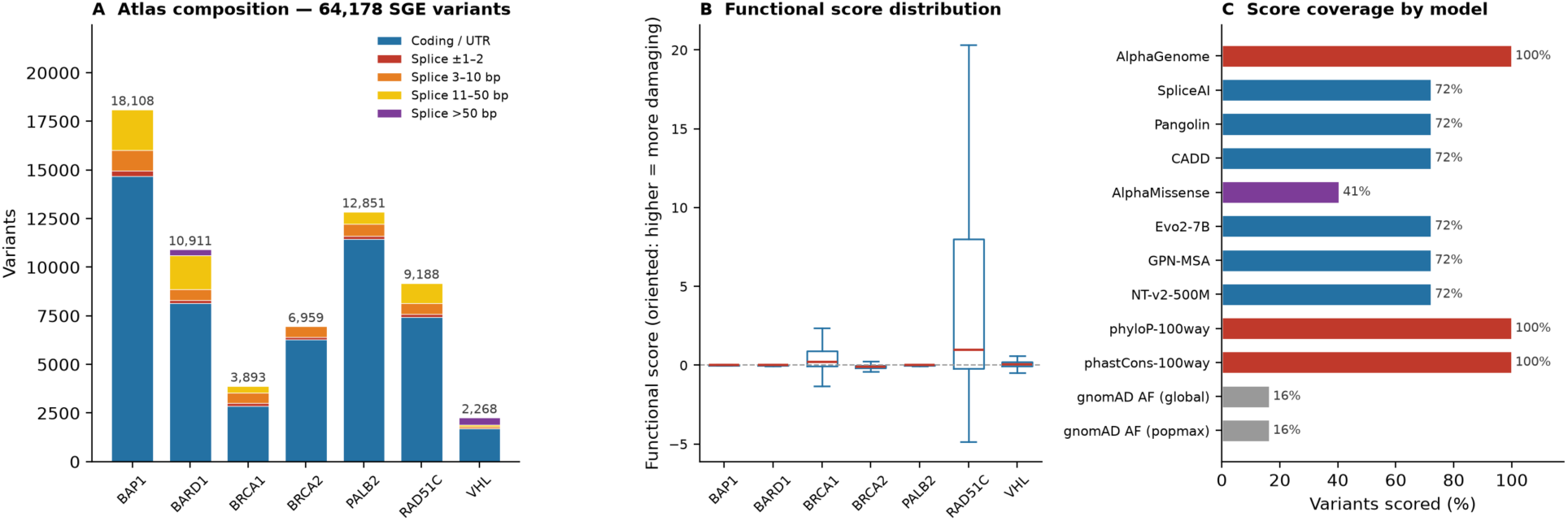
The functional-standard atlas: 64,178 variants across seven genes, nineteen scored predictors. (*A*) Variants per gene, stacked by region class. (*B*) Distribution of oriented functional scores per gene (box: median and interquartile range; whiskers 1.5 × IQR; outliers omitted; higher = more damaging). Raw scales differ between assays — per-gene standard deviations span 0.062 to 6.84 — which is why all analyses correlate within gene. (*C*) Score coverage per predictor as a percentage of atlas variants.

**Table 1.** Composition of the frozen functional-standard atlas. Per-assay gene, MaveDB accession, MANE-Select transcript and variant counts by region class. Indels (17,786) are the difference between the Variants and SNVs columns. Primary assay publications: *BRCA1* [19], *BAP1* [35], *RAD51C* [36], *VHL* [37], *BRCA2* [38], *PALB2* [39], *BARD1* ([40], preprint); each deposit is additionally cited as a data reference [41–47]. Nine *BRCA1* variants in an intron within the 5’UTR are counted as splice variants (six at offsets −1 and −2 fall in the ±1–2 stratum; the three at −3 fall in 3–10 bp); an earlier form of our region classifier filed them as coding (Methods). *BRCA2* targets exons 15–26 with only limited flanking intron coverage, which is why it contributes no variants to the 11–50 bp and >50 bp strata.

| Gene | MaveDB URN | MANE-Select<br>transcript | Variants | SNVs | Coding/<br>UTR | Splice<br>$\pm 1-2$ | Splice<br>3-10<br>bp | Splice<br>11-50<br>bp | Splice<br>>50 bp |
| --- | --- | --- | --- | --- | --- | --- | --- | --- | --- |
| *BAP1* | urn:mavedb:00000662-0-1 | NM_004656.4 | 18,108 | 9,441 | 14,685 | 277 | 1,053 | 2,093 | 0 |
| *BARD1* | urn:mavedb:00001250-a-2 | NM_000465.4 | 10,911 | 8,818 | 8,142 | 147 | 577 | 1,749 | 296 |
| *BRCA1* | urn:mavedb:00000097-0-2 | NM_007294.4 | 3,893 | 3,893 | 2,859 | 143 | 530 | 361 | 0 |
| *BRCA2* | urn:mavedb:00001225-a-1 | NM_000059.4 | 6,959 | 6,959 | 6,270 | 138 | 551 | 0 | 0 |
| *PALB2* | urn:mavedb:00001259-a-2 | NM_024675.4 | 12,851 | 10,375 | 11,428 | 184 | 614 | 625 | 0 |
| *RAD51C* | urn:mavedb:00000673-0-1 | NM_058216.3 | 9,188 | 4,638 | 7,445 | 143 | 547 | 1,053 | 0 |
| *VHL* | urn:mavedb:00000675-a-1 | NM_000551.4 | 2,268 | 2,268 | 1,694 | 24 | 83 | 80 | 387 |
| **Total** |  |  | **64,178** | **46,392** | **52,523** | **1,056** | **3,955** | **5,961** | **683** |

Nineteen predictors were scored under pinned pipelines, with per-predictor coverage of the atlas shown in Fig. 1C (Additional file 1: Table S9). Two of them, SpliceAI and Pangolin, were re-scored at full floating-point precision for this work: both command-line tools compute in float and round only when formatting output, which placed 47% and 54% of coding variants on a single value and bounded Spearman ρ below 1 for arithmetic reasons alone. The re-scores reproduce the released columns exactly when re-rounded (Additional file 1: Table S5b), and raise the number of distinct values from 101 to 46,130 and from 92 to 46,237 respectively (Methods). Tie structure and the tie-imposed bound it places on Spearman ρ, before and after re-scoring, are tabulated for every model × stratum cell in Additional file 1: Table S5.

### The territory map, and what the assays can measure

Figure 2A gives the conventional result: pooled Spearman ρ between each predictor and the functional score, by stratum. Rankings invert across territories. Of the twelve primary predictors (the nineteen minus the seven predominantly missense dbNSFP meta-predictors; Methods), AlphaMissense leads the All stratum (0.438, 95% confidence interval (CI) 0.325–0.538) but scores only missense SNVs. Among broad-scope predictors CADD is strongest (All 0.391, 0.322–0.457; coding/UTR 0.427), followed by GPN-MSA (0.363) and Evo2-7B (0.358). Conservation occupies the middle tier (phyloP 0.292; phastCons 0.254), the splice-specialised models are weak overall (AlphaGenome 0.146; SpliceAI 0.177; Pangolin 0.190) because their signal is concentrated in one territory, and NT-v2-500M is uniformly weak (0.098). Heterogeneity between genes is substantial (I² up to 99%); per-gene estimates and forest plots for every model × stratum combination are given in Additional file 2: Figure S1–S12 and Additional file 1: Table S1.

**Figure 2.**
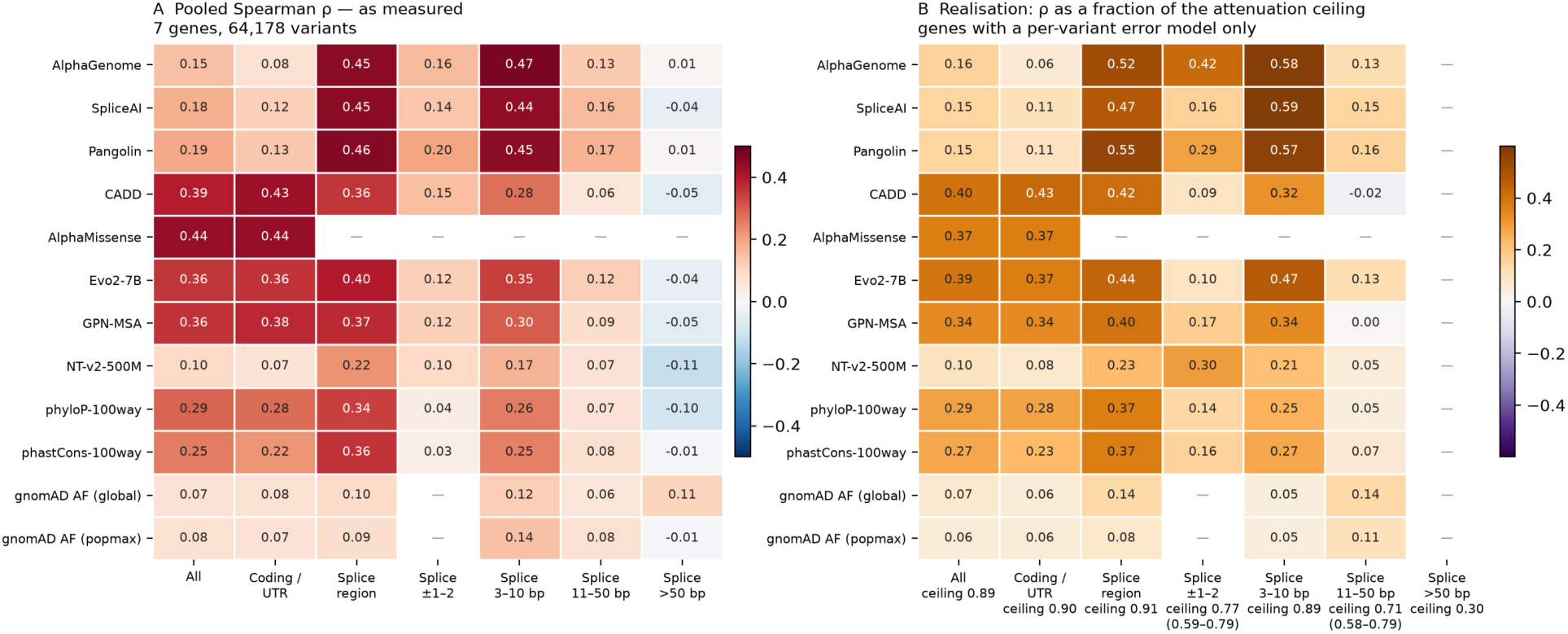
The territory map as measured, and as a fraction of what the assays can measure. (*A*) Pooled Spearman ρ ([48], over seven genes, on Fisher-z transforms) across the 64,178 atlas variants; cells are point estimates; their 95% CIs (DerSimonian–Laird on Fisher-z) are tabulated in Additional file 1: Tables S1 and S2; variant counts per stratum are in Table 1 and coverage in Additional file 1: Table S1. All-stratum cells use each predictor’s scored footprint. Em dashes mark strata a predictor does not score. (*B*) The same grid divided by each stratum’s attenuation ceiling (column headers), from replicate scores (*BRCA1*) or CI-validated standard errors (*BARD1*, *PALB2*). Each cell is a realisation (observed ρ ÷ ceiling); the diverging colour scale is centred at zero, negative values marking cells above the ceiling. Where the three genes disagree by ≥0.10 the ceiling range is given beneath the median. Cells appear only where at least two genes contribute an error model (Methods), blanking the >50 bp column: its ceiling (0.30) rests on *BARD1* alone and lies below the correction’s reliable range. The apparent collapse at ±1–2 in *A* is largely absent in *B*.

Read alone, that grid invites a conclusion it does not support, because the strata differ not only in how hard they are to predict but in how precisely the assays measure them. Three deposits let us quantify this: *BRCA1* publishes replicate scores whose mean reproduces the deposited score, and *BARD1* and *PALB2* per-variant standard errors that reconcile with their published intervals. From these we estimated the reliability of the functional score in each stratum, and hence the attenuation ceiling — the largest correlation any predictor could achieve there. We call the ratio of an observed ρ to that ceiling its realisation, the attenuation-corrected scale used throughout. *BRCA1* carries both estimators and the SE-based one runs 0.049–0.062 higher (per-stratum values in Additional file 1: Table S14), a small consistent optimism that makes the corrected values below conservative. Two further deposits carry repeated measurements that are not clean replicates but bound reliability in known directions, and both show reproducibility falling with intronic offset. In *VHL* two selection conditions of the same assay agree at ρ = 0.045 deep intronic against 0.389 in coding (Additional file 2: Note S5; per-gene reliability and realisation values in Additional file 1: Table S3).

The ceiling varies far more across territory than the predictors do (Fig. 2B): 0.90 in coding/UTR and 0.89 at splice 3–10 bp, but 0.77 at splice ±1–2, 0.71 at 11–50 bp and 0.30 in the deep intron. The three genes agree closely in most strata but disagree by more than 0.20 at ±1–2 and 11–50 bp, so ceilings there are reported as ranges and the realisations inherit that spread.

### The correction reduces error above a ceiling of about 0.45 and amplifies it below

Dividing by an estimated ceiling is defensible only if it recovers the error-free correlation, and the classical result assumes Pearson correlation with normal error while we apply it to Spearman ρ on assay scores that are far from normal and, at canonical splice sites, severely compressed. We therefore tested the whole chain — estimator included — by simulation (Methods). Across 151,200 trials the correction removes almost all of the attenuation: uncorrected, ρ underestimates the error-free correlation by 0.038 to 0.298 as reliability falls; corrected, the residual bias lies between −0.031 and +0.007 at every level of that grid (Additional file 1: Table S4). It survives compression, the property we were most concerned about — at splice ±1–2, the most compressed stratum, the corrected bias is +0.006.

The simulation also identifies a boundary the correction should not cross. Above a ceiling of roughly 0.45 it reduces root mean squared error as well as bias, though by how much depends strongly on the ceiling itself: 14% at 0.451, and 37–49% once the ceiling exceeds about 0.64. Below the boundary, dividing by a small and noisily estimated ceiling amplifies error; at a ceiling of 0.28 the corrected estimate is 1.7 times worse than the uncorrected one. Of the strata measured here only the deep intron, at 0.30, falls below that boundary, which is why it is reported uncorrected throughout. The full simulation grid is given in Additional file 1: Table S4.

Three refinements sharpen that boundary (Additional file 1: Table S15). First, a denser ten-level grid locates the root-mean-squared-error crossover of the Gaussian replicate-based chain at an achieved ceiling of 0.34–0.40 (target reliability 0.15–0.20), so the 0.45 rule sits just above the crossover rather than at it; nor is the crossover uniform across stratum shapes — coding/UTR improves at every level on the grid, while the crossover falls at target reliability 0.30–0.50 for splice ±1–2, 0.15–0.20 for 3–10 bp, 0.25–0.30 for 11–50 bp and 0.20–0.25 for the deep intron. Second, what fails below 0.45 is error amplification, not bias: at an achieved ceiling of 0.274 on the denser grid the corrected bias is only +0.006, and because a nominal reliability of 0.10 yields an achieved ceiling of just 0.274, the Spearman–Brown estimator itself underestimates reliability in this regime — part of the boundary is estimator failure rather than correction failure. Third, the boundary is robust to the noise model with one exception: Gaussian and bounded-truncation noise give similar crossovers; score-dependent noise, with the standard error growing in |score| and calibrated on the *BRCA1*-derived SEs, pushes the crossover later (target reliability 0.25–0.30) and degrades more heavily below it; and under bounded noise the SE-based chain is mis-specified — it is fed the nominal pre-truncation SE — and collapses at low target reliability, its corrected error diverging, the only combination that fails qualitatively rather than by degree. The extended grid also separates the two estimators: at low reliability the SE-based chain tracks the true ceiling more closely and corrects with smaller error than the replicate-based chain, while at mid-to-high reliability it systematically overestimates the ceiling (Gaussian-noise rows at the 0.95 level: bias −0.028 against the replicate-based chain’s −0.007), the same direction as the *BRCA1* cross-check above.

### Most of MaveDB permits this estimate, and the ceilings it yields vary widely

The prescription is only worth making if deposits carry what it needs, so we surveyed all 2,803 published MaveDB score sets rather than generalising from ours. 2,452 (87%) publish either a per-variant error estimate or replicate scores whose mean reproduces the deposited score; 29 publish replicate columns that reconcile with no subset of themselves, and 322 carry neither. These counts are upper bounds — the column-name test accepts on name alone, without the content reconciliation the primary analysis applies (Methods).

That figure is not reachable by the obvious search. A conventional vocabulary of column names — standard error, standard deviation, confidence interval — returns 247 deposits. Inspecting sixty deposits it had classified as carrying nothing showed the shortfall was in the vocabulary, not the resource: ‘sigmà was absent from it, a bare ‘err’ token appeared only inside ‘stderr’, and an interval written ‘score_95CI_low’ places digits where the pattern expected a word boundary. Adding those three forms, each taken from a column name actually observed, recovers the 2,452. Error models in MaveDB are common but not conventionally named, which is why a benchmark author is unlikely to find them. The audit ran in both directions: the sixty deposits classified as carrying nothing were inspected and adjudicated, and a symmetric sample of twenty-five classified as carrying errors was drawn, whose adjudication is still in progress — so the false-negative direction is bounded here and the false-positive direction is not yet.

The accounting in full: 1,202 of the 2,803 published score sets are human, and 991 of those (82.4%) permit an estimate on the survey test; 728 human deposits enter the ceiling computation under our counting rules, and 674 of them yield a usable value (92.6%; Fig. 3) — 24.0% of the entire resource. The median is 0.895 and 51.8% fall below 0.90. 530 of the 674 carry a column named as a standard deviation, which we treat as a standard error — conservative if it is the spread across replicates rather than the error of their mean. We therefore also report the 144 deposits carrying an explicit standard error, reconcilable replicates or a confidence interval (124, 16 and 4 respectively): median 0.949, with 29.9% below 0.90. The true figure lies between, and the strict subset may also be a selected one — deposits whose authors documented their uncertainty carefully may differ in other respects — so the separation between the two readings (Kruskal–Wallis P = 2.2 × 10⁻¹⁰) cannot distinguish the convention effect from a selection effect. Either reading supports the same conclusion: a benchmark pooling across these datasets is comparing correlations with materially different upper bounds. On either reading 3.1% (21 of 674) fall below the 0.45 bound our simulation sets for the correction, and for those the achievable correlation is so low that a benchmark score computed on them is close to uninterpretable. That share is conservative in the SE convention. Treating a deposited standard deviation as the error of a mean of k replicates (SE = SD/√k) moves the below-0.45 share for the 606 sd_as_se deposits — every surveyed deposit whose error column is a standard deviation, not only the human counting class the 530 above sits in — from 3.9% at k = 1 to 0.2% at k = 2 and 0.5% at k = 3, and for the whole distribution of 730 deposits excluding the designed-stability platform — the human counting class together with the other natural-sequence deposits — from 3.3% to 0.4% and 0.7% (Additional file 1: Table S16); the share is not monotone in k because the set of deposits with a computable ceiling itself changes with k. Thirty deposits return a reliability at or below zero, meaning the published error column is not on the scale of the score it accompanies. The spread does not track experiment size or year of deposition (Additional file 2: Figure S17), so it is not a problem that later data will dissolve. It does track which error column was published (Additional file 2: Figure S17C), which is what the two readings above reflect.

**Figure 3.**
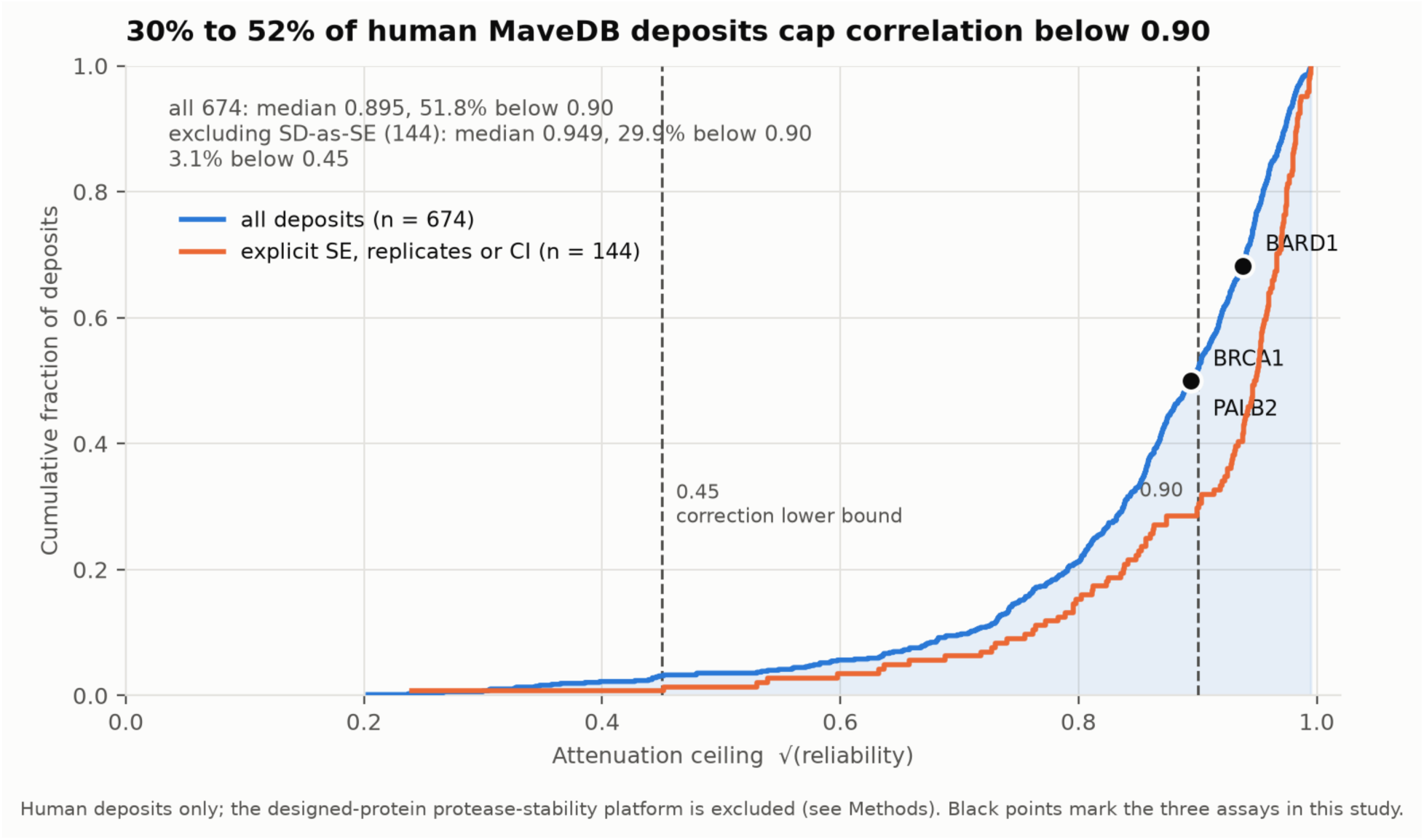
Attenuation ceilings across human MaveDB deposits. Cumulative distribution of the ceiling for the 674 human deposits that yield a usable ceiling (blue), and for the 144 of them whose error column is an explicit standard error, reconcilable replicates or a confidence interval rather than a deposited standard deviation used as one (orange). The gap between the curves is the effect of that convention. Both curves are empirical cumulative distributions of per-deposit point estimates and carry no confidence intervals. Dashed lines mark the 0.45 bound below which our simulation shows the correction should not be used, and 0.90. Black points mark the three assays in this study that carry a validated error model. The designed-protein protease-stability platform is excluded: its targets are not human variants and its published interval is a curve-fitting confidence interval on a thermodynamic parameter rather than a reproducibility interval (Methods).

Our three ceiling-bearing assays sit at the middle of that distribution: *BRCA1* and *PALB2* at the 50th percentile (0.894) and *BARD1* at the 68th (0.938). The demonstration that follows is therefore run on typical human data rather than on unusually precise or unusually noisy examples. Two of the seven fall out under the resource-wide rules for the reasons the primary analysis excluded them by hand: *BAP1*’s standard errors return a negative reliability and *RAD51C*’s are on a different scale from its score. That the general rule reproduces the case-by-case decision is a check on both.

### The collapse at canonical splice sites is a measurement ceiling, not a predictor failure

Every predictor scores poorly at splice ±1–2 as measured, from 0.198 (Pangolin) down to 0.033 (phastCons) (Fig. 2A). The functional data explain why the ceiling is low there. Expressed as a within-assay percentile, ±1–2 variants have a median percentile of 0.906 (interquartile range 0.841–0.957) and 84.5% lie above the 80th percentile of their own assay, against a median of 0.514 in coding/UTR: canonical disruption is near-deterministically damaging, the residual variance is small, and the assays reproduce it correspondingly poorly (reliability 0.60, ceiling 0.77).

What the correction shows is that predictors are not doing worse there than elsewhere. The comparison must be made on a matched set, since the predominantly missense meta-predictors cover too few splice variants to compare and three further columns have no ±1–2 estimate; nine broad-scope predictors carry one in both strata, over the three genes with a validated error model. As measured, coding/UTR exceeds ±1–2 by 1.63 on the best predictor in each territory (0.404 for CADD against 0.248 for AlphaGenome) and 1.69 on the median (0.219 against 0.130). After correction the gap at the frontier closes — 0.429 against 0.423, a ratio of 1.02 — while the median gap narrows to 1.43 (0.233 against 0.163) rather than closing. The frontier ratio also carries a winner’s-curse caveat: it takes the best predictor per territory without cross-validation, so the sampling properties of 1.02 are unknown, and it rests on the *BARD1* preprint and on the SE-based estimator.

The median convergence, unlike the frontier parity, survives stress-testing — with one exception. Leave-one-gene-out over the three ceiling-bearing genes (Additional file 1: Table S13; the pooled-ρ leave-one-gene-out swings of the underlying correlations are tabulated in Additional file 2: Note S7) gives uncorrected/corrected median ratios of 1.69/1.43 on the full set, 1.42/1.25 dropping *BARD1* and 2.41/1.61 dropping *PALB2* — the convergence survives either — but 1.09/0.78 dropping *BRCA1*: the corrected ratio inverts below 1, the corrected ±1–2 median rising above coding/UTR. The mechanism is cross-gene ceiling spread, not estimator bias: the three ±1–2 ceilings are 0.772 (*BRCA1*, the only replicate-based one), 0.794 (*PALB2*) and 0.586 (*BARD1*), and dropping *BRCA1* moves the median ceiling from 0.772 to about 0.69, lifting the corrected ±1–2 realisations past coding/UTR. The SE-based estimator’s small consistent optimism (0.049–0.062; Additional file 1: Table S14) points the other way — it inflates ceilings and therefore shrinks corrected values — so it is conservative here and cannot explain the inversion; were it the cause, the inversion would be stronger still. Holding the estimator uniform instead — SE-based ceilings for all three genes — moves the corrected median ratio only from 1.43 to 1.45 and the median ±1–2 ceiling from 0.772 to 0.794, the conservative direction. We therefore report all three folds rather than claim gene independence. The frontier figure remains the fragile half of the result: 0.423 is the median of three dispersed per-gene values and is the one *BARD1* contributes, and excluding *BARD1* leaves 0.305 against 0.429, a ratio of 1.41.

The widely repeated reading — that models fail at canonical sites — nonetheless attributes to the models a limit belonging substantially to the measurement: a low score at ±1–2 is uninformative because the assay can barely resolve variation there, not because the predictor has broken down.

### Where predictors genuinely fail: 11–50 bp, and an unmeasurable deep intron

The correction relocates the failure to 11–50 bp. At 11–50 bp the ceiling is 0.71, estimated from the same three genes that carry an error model, and the best realisation is 0.156 (Pangolin), with a median across predictors of 0.109 — by a wide margin the worst-served territory in the atlas, and one the uncorrected map presents as merely modest. This is where better models would most clearly help.

The deep intron is a different problem. Its ceiling is 0.30, from *BARD1* alone, below the boundary at which correction is reliable; the stratum is small (n = 683) and confined to two genes; and the minimum ρ detectable at 80% power is 0.107 pooled. Observed values are Pangolin 0.008, AlphaGenome 0.010 and SpliceAI −0.042, with small negatives for phyloP (−0.095) and NT-v2-500M (−0.113) concordant across the two contributing genes rather than a pooling artefact (I² = 0). Estimated against a ceiling of 0.30, their sign is more trustworthy than their magnitude, which we do not interpret. We state the stratum as a limit of the current functional standard rather than a property of the predictors: what these assays can support there is bounded at 0.30, below the range in which the correction is reliable, so a benchmark cannot conclude from it that sequence models are blind deep in introns. Intronic saturation editing at greater depth, not better models, is what would settle it. Quantisation is not the explanation: at full precision the tie-imposed bound rises to 1.000 for both SpliceAI and Pangolin here and the estimates remain at zero (Additional file 2: Note S6).

### Ordering claims require paired tests

Within the splice region the three dedicated splice-aware predictors form a tight group (Pangolin 0.458, SpliceAI 0.451, AlphaGenome 0.451), with Evo2-7B fourth (0.395). Resolving by intronic offset, signal peaks at 3–10 bp — AlphaGenome 0.470, Pangolin 0.447, SpliceAI 0.444 — and decays monotonically thereafter, reaching zero beyond 50 bp (Additional file 2: Figure S14). Because the offset bins have different attenuation ceilings (Fig. 2B), that decay is steeper as measured than in realised terms.

Whether the ordering at the peak is real is a separate question, and overlapping marginal intervals cannot answer it: two correlations computed on the same variants are dependent, and the marginal intervals are far wider than the uncertainty in their difference. We tested each ordering claim pairwise on the variants both predictors score, using Steiger’s test [49] pooled across genes with a gene-cluster bootstrap for the difference on the ρ scale (Fig. 4A). AlphaGenome’s lead over SpliceAI at 3–10 bp is not resolvable (Δρ = 0.014, 95% CI −0.012 to 0.036, P = 0.21), nor is its lead over Pangolin (P = 0.65), nor Pangolin’s over SpliceAI at 11–50 bp (P = 0.67). At the peak of splice territory the three dedicated predictors are, on this standard, interchangeable. All pairwise tests are listed in Additional file 1: Table S6. Steiger and bootstrap P values agree at α = 0.05 for all eighteen pairs (Additional file 1: Table S17).

**Figure 4.**
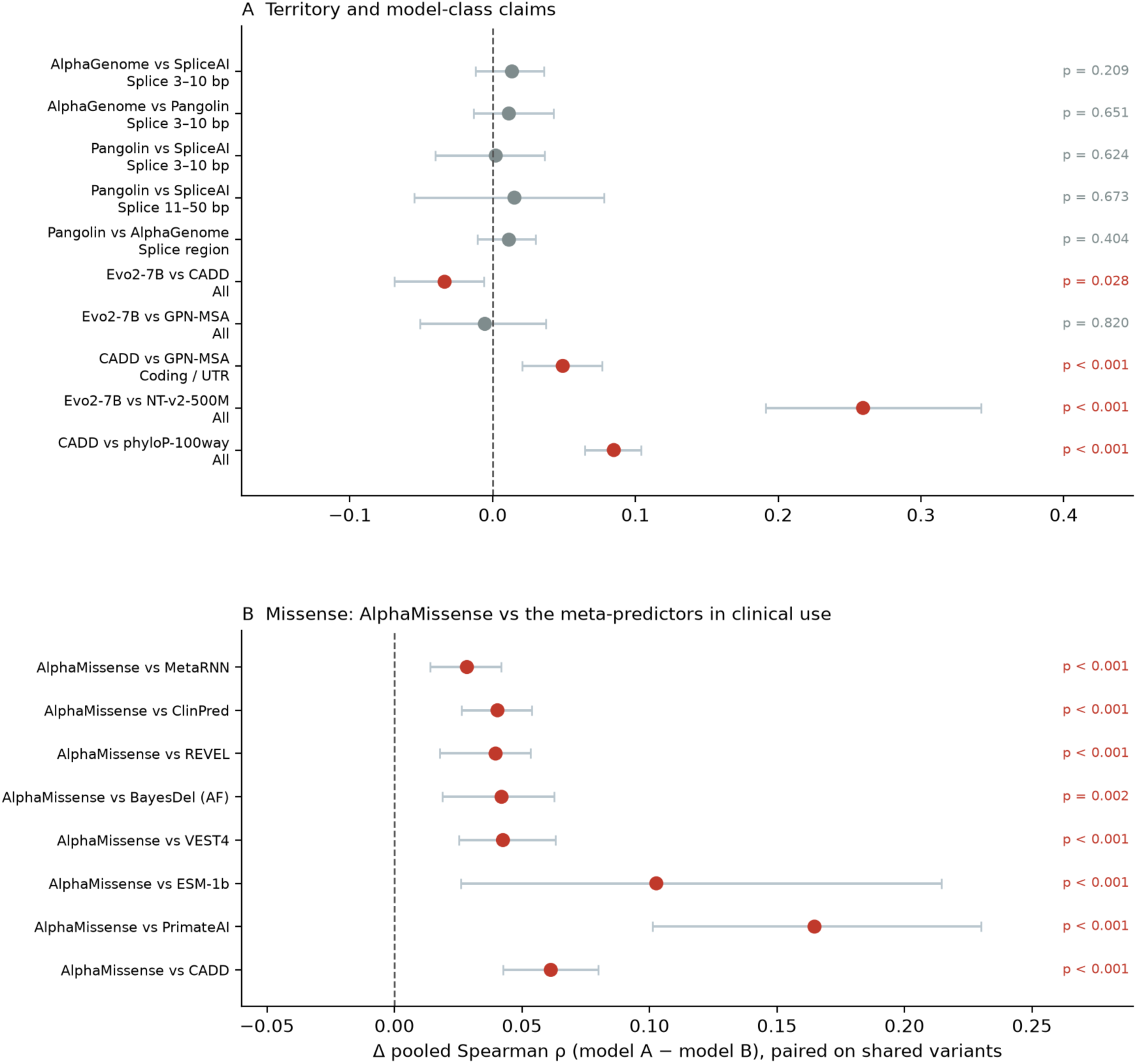
Ordering claims tested pairwise. Δ pooled Spearman ρ between two predictors computed on the variants both score — 3,290 to 46,392 depending on the pair and stratum, listed per comparison in Additional file 1: Table S6 — with a gene-cluster bootstrap 95% CI; *P* from Steiger’s test for dependent correlations, DerSimonian–Laird pooled over genes. Red marks differences resolvable at α = 0.05. (*A*) Territory and model-class claims. (*B*) AlphaMissense against the seven meta-predictors in clinical use, within the missense stratum. Because the two correlations are measured on the same variants they are dependent, so overlap of their marginal intervals is not a test of their difference — the paired test is both more powerful (*B*, and Evo2-7B versus CADD) and more conservative (AlphaGenome at 3–10 bp) than reading Fig. 2A.

The same test resolves a comparison that interval overlap conceals. Evo2-7B, a 7-billion-parameter model scored as a windowed log-likelihood delta using no supervision, conservation track or annotation, reaches ρ = 0.358 (0.300–0.413) overall. It is statistically indistinguishable from GPN-MSA (Δρ = −0.005, P = 0.82), which is trained with alignment context, so a raw-sequence model matches an alignment-based one on experimentally measured function. It does not match CADD: the marginal intervals overlap, but paired on the same 46,392 variants the gap is resolvable (Δρ = −0.034, 95% CI −0.069 to −0.006, Steiger P = 0.028; gene-cluster bootstrap P = 0.0005 — the closest case in the grid, and the bootstrap is the stronger of the two). NT-v2-500M, the same model class 14 times smaller, reaches 0.098 and is 0.259 behind Evo2-7B (P < 0.001).

### AlphaMissense leads every meta-predictor in clinical use

The integrative class is usually represented in benchmarks by one or two scores, which makes claims about it fragile. We added the seven dbNSFP meta-predictors that clinical pipelines use (Methods), predominantly missense, and compared them within the missense stratum. AlphaMissense leads (0.438), followed by MetaRNN (0.409), ClinPred (0.397), REVEL (0.397), BayesDel (0.396) and VEST4 (0.394), with CADD at 0.376, ESM-1b at 0.344 and PrimateAI at 0.272. NT-v2-500M reaches −0.0003, indistinguishable from zero.

Every marginal interval in that list overlaps AlphaMissense’s, and reading them alone would place the nine scores in one indistinguishable group. Paired on shared missense variants, all eight differences resolve (Fig. 4B): versus MetaRNN Δρ = 0.028 (bootstrap 0.014–0.042, P < 0.001), versus REVEL 0.040 (P < 0.001), versus BayesDel 0.042 (P = 0.002), versus CADD 0.061 (P < 0.001), versus PrimateAI 0.165 (P < 0.001). All survive Benjamini–Hochberg adjustment across the full grid of comparisons (Methods). This is the clearest illustration in the dataset of why paired testing is required. Even so, the best of them realises only about half of the ceiling available in coding sequence.

### Model ranking is a property of the readout, not only of the variant

Four deposits publish a second, mechanistically distinct readout of the same variants — RNA abundance of the variant allele — alongside the cell fitness score the atlas freezes. It covers coding/UTR variants only (19,553 across four genes), so it cannot re-test splice territory, but it supplies what a benchmark rarely has: a same-experiment estimate of how far two legitimate measurements of one variant can diverge.

They diverge a long way. Pooled across the four genes the two readouts agree at ρ = 0.251 (0.124–0.369) — lower than five of the twelve primary predictors score against the fitness readout on the same variants (AlphaMissense 0.424 the highest). A predictor agreeing with a saturation editing readout at ρ ≈ 0.4 is therefore performing at, and here above, the level at which the experiment agrees with itself across modalities — a better yardstick than an implicit comparison against 1.0.

Against the RNA readout the ranking inverts (Fig. 5). Every coding-oriented predictor collapses — AlphaMissense 0.424 to 0.095, CADD 0.412 to 0.165 — while two of the three splice-aware models gain — AlphaGenome from 0.087 to 0.100 and Pangolin from 0.133 to 0.163 — the only two of the nineteen to do so; SpliceAI is the exception, falling from 0.141 to 0.109. All three sit near the bottom against fitness in coding regions, seventh to ninth of twelve. This is not noise: *BRCA1* publishes RNA replicates, giving that readout a reliability of 0.611 and a ceiling of 0.782, so CADD realises 21% of the achievable RNA signal against 46% of the achievable fitness signal. The interpretation is mechanistic — in coding sequence, what moves RNA abundance is splicing and nonsense-mediated decay, which only the splice-aware models model — and it is corroborated independently by the synonymous stratum below.

**Figure 5.**
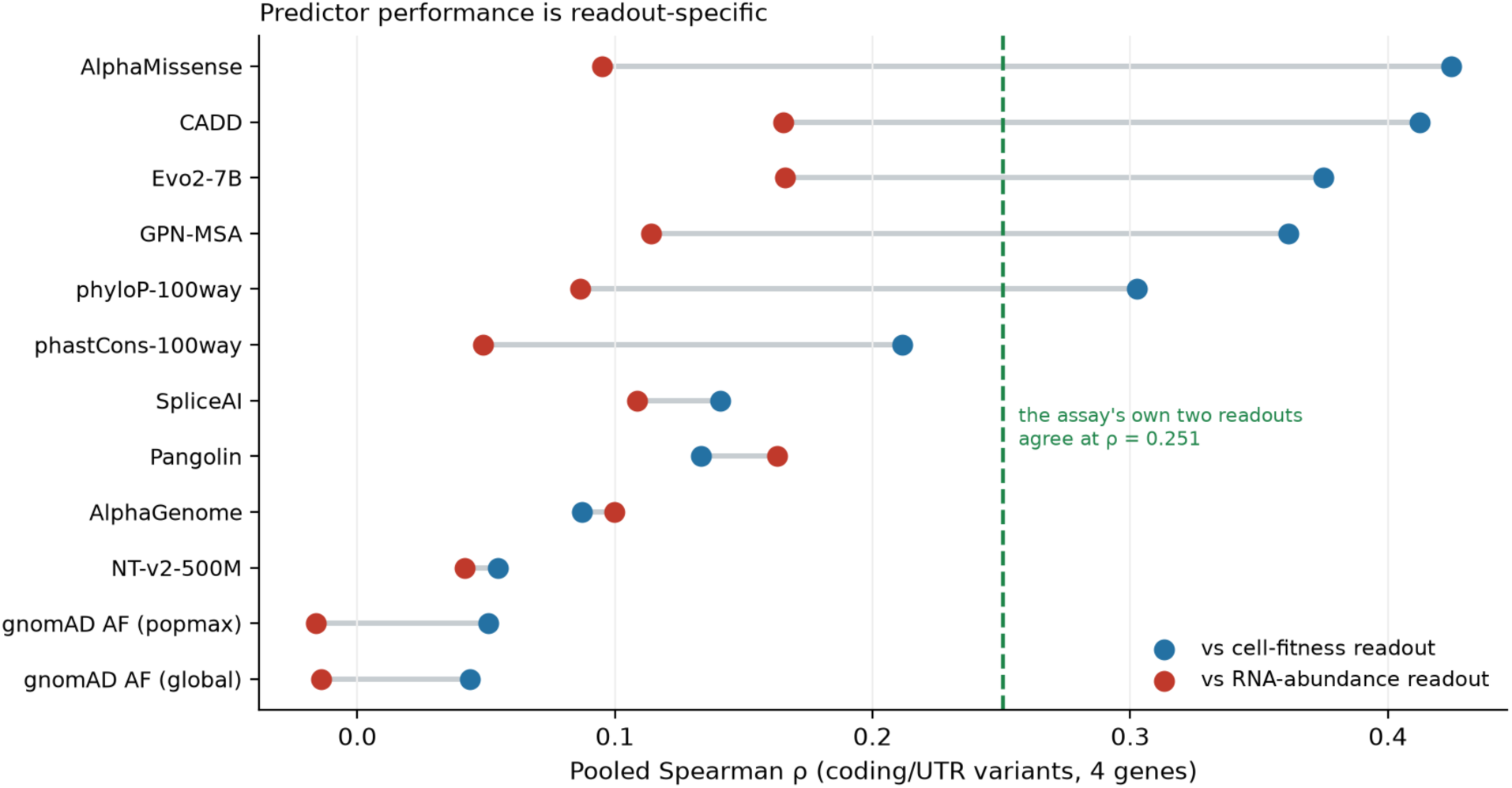
Predictor performance is readout-specific. Pooled Spearman ρ against the cell fitness readout (blue) and the RNA abundance readout (red) of the same assays, on the same 19,553 coding/UTR variants across four genes. Points are DerSimonian–Laird pooled estimates on Fisher-z transforms, as in Fig. 2A. The dashed line is the agreement between the two readouts themselves, ρ = 0.251 (95% CI 0.124–0.369).

### Classification performance, and how much evidence it is worth

Rank correlation asks whether a predictor orders variants by effect size; a diagnostic laboratory asks whether it separates damaging from tolerated. Three deposits publish the assay authors’ own functional calls, giving 31,835 labelled variants across *BARD1*, *PALB2* and *RAD51C*. Scored as classification the same predictors look far stronger than ρ suggests (Fig. 6A): AlphaGenome reaches an area under the receiver operating characteristic curve (AUROC) of 0.963 (0.949–0.974) in splice territory where its ρ is 0.451, and CADD reaches 0.916 (0.890–0.935) in coding/UTR where its ρ is 0.427. The two metrics are not in conflict — a predictor can separate two modes cleanly while ordering poorly within them — but a ρ-based territory map understates binary separation of the assays’ own functional calls and should not be read as one; correlation with a functional assay tracks clinical classification performance in its own right [50].

**Figure 6.**
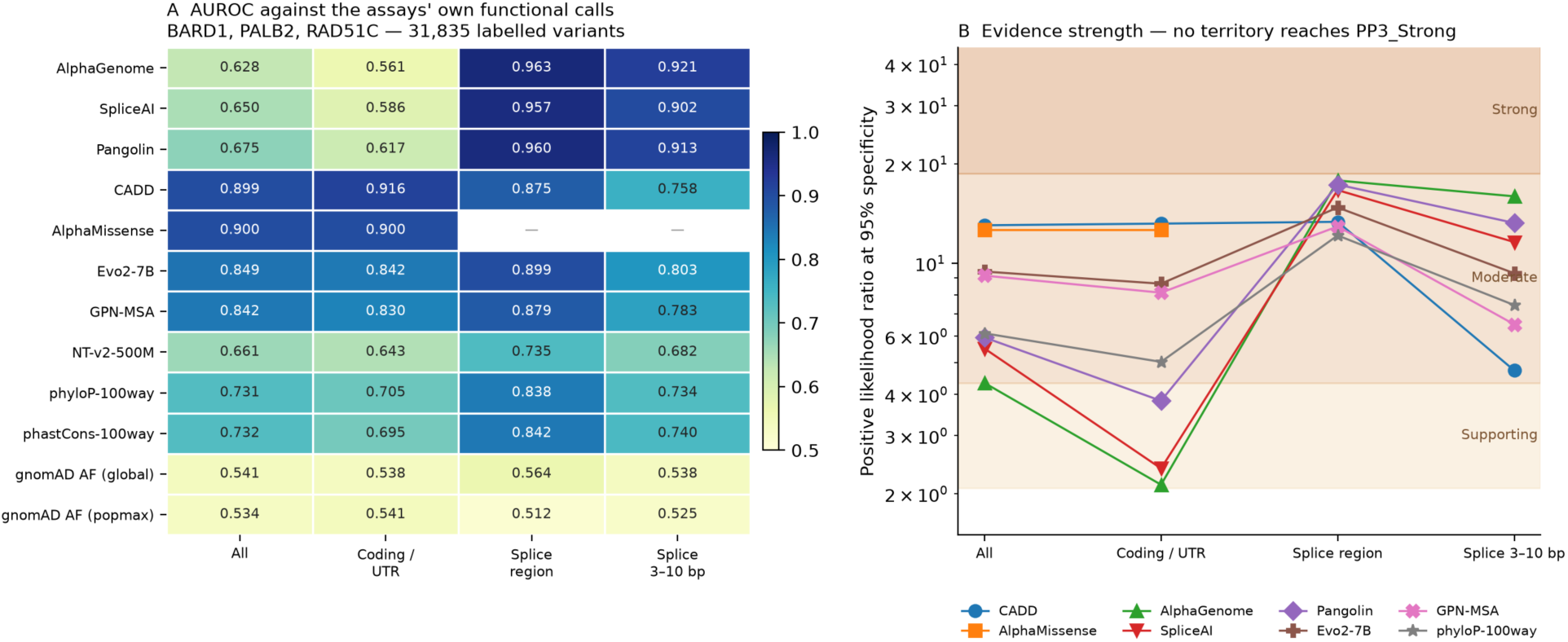
Classification performance and evidence strength by territory. (*A*) AUROC against the assays’ own functional calls (31,835 labelled variants in *BARD1*, *PALB2* and *RAD51C*), pooled across genes on the logit scale with Hanley– McNeil variances [51]. (*B*) Positive likelihood ratio at 95% specificity against Tavtigian-point evidence bands at a prior of 0.10. No predictor in any territory reaches the Strong band. Panel B is indicative only, not a ClinGen-grade calibration: at exactly 95% specificity the positive likelihood ratio cannot exceed 20 by construction, and reaching the Strong threshold (18.7) there requires sensitivity above 0.93. Lines connect discrete territories for readability only; no continuity is implied. Panel B plots the eight predictors for which an evidence-strength mark is shown; phastCons, the two gnomAD allele-frequency columns and NT-v2-500M are not plotted, and their likelihood ratios at 95% specificity are given in Additional file 2: Note S9. The seven predominantly missense dbNSFP meta-predictors enter the same analysis in the All and coding/UTR territories and are likewise tabulated, but not plotted, there.

Converting to the scale clinical guidelines use turns the same limit into a statement about calibration. At 95% specificity the positive likelihood ratio reaches 17.8 for AlphaGenome, 17.3 for Pangolin and 16.6 for SpliceAI in splice territory, and 13.2 for CADD in coding/UTR (Fig. 6B). Under the Tavtigian point system [30] at a prior of 0.10 these fall in the ACMG Moderate band: on this functional standard no predictor in any territory reaches the likelihood ratio of 18.7 required for PP3_Strong, and the three that come closest fall just short. The four ratios quoted above are read at the most sensitive observed score threshold reaching 95% specificity (realised specificity 95.0–95.3% in every gene) and reported as the per-gene median; interpolating each to exactly 95% specificity — the basis of the companion splice-region study [52] — moves it by at most 0.35 in LR+, and no evidence band in Fig. 6B changes (Additional file 2: Note S15). Coarsely quantised conservation and allele-frequency scores are not interpolated: where no observed threshold reaches 95% specificity they are reported as not evaluable, and otherwise at the specificity they achieve.

The general point follows from what these evidence strengths are derived against. A PP3/BP4 calibration built on experimental labels inherits the precision of that experiment in each territory it covers, because the labels themselves are noisier where the assay is noisier. A single evidence weight assigned to a predictor across all variant classes therefore silently averages over territories in which the underlying measurement differs in achievable correlation by a factor we here estimate at up to three. Two consequences follow: such a calibration should be performed per territory, or the reliability of the labels it rests on should be reported alongside it. What we report here is not a ClinGen-grade calibration: it rests on assay-derived labels in three genes and lacks the prior-probability model and bootstrap procedure of Pejaver et al. [32]. It does identify a precondition that any such calibration has to satisfy, before its evidence weights can be read as properties of the predictor rather than of the assay behind the labels.

### ClinVar ascertainment, consequence class, and the structure of the predictor space

Because ClinVar-recorded variants are the evaluation set of most published benchmarks, the gap between performance there and elsewhere estimates the optimism they carry. The informative comparison is against the 24,998 SNVs ClinVar has *not* recorded, not the full atlas, which contains the recorded ones and dilutes the contrast. Recorded versus absent, the median uplift across predictors is 0.082 — two and a half times the 0.033 obtained from the diluted comparison; CADD, for instance, falls from 0.445 to 0.312. The full grid is in Additional file 1: Table S2 and Additional file 2: Note S8.

A predictor could appear strong in coding sequence simply by separating nonsense from missense variants. It does not: within the missense stratum alone CADD retains 0.376 of its 0.427 coding/UTR value, with the ranking preserved. The synonymous stratum is another matter: all predictors are effectively blind to it (CADD 0.023) even though these 7,886 variants carry measurable functional effects, and the splice-aware models are the best performers there despite ranking seventh, eighth and ninth of twelve in coding sequence overall, corroborating the RNA-readout result (Additional file 2: Note S8).

The between-model rank-correlation matrix in coding sequence shows that the nine broad-scope and splice-aware predictors span essentially two axes: a dense cluster of generalists and a largely independent cluster of splice-aware models, correlating at 0.14–0.25 across the two against 0.43–0.82 within them (full matrix in Additional file 2: Note S9). NT-v2-500M sits on neither axis: it correlates with nothing (maximum 0.16), and grouping it with the generalists would place its within-cluster correlations below the cross-cluster ones. Selection strategies were scored leave-one-gene-out, so that no strategy is evaluated on data used to choose it. Territory-aware selection is worth a great deal in splice territory: at 3–10 bp it returns 0.448 against 0.257 for using CADD everywhere. It is worth nothing in coding, where the selector picks CADD in all seven folds; the mean within-gene percentile rank of the three splice-aware models does better still, winning every splice stratum (Additional file 2: Figure S15; per-fold values in Additional file 1: Table S7). Table 2 states the resulting guide.

**Table 2.**
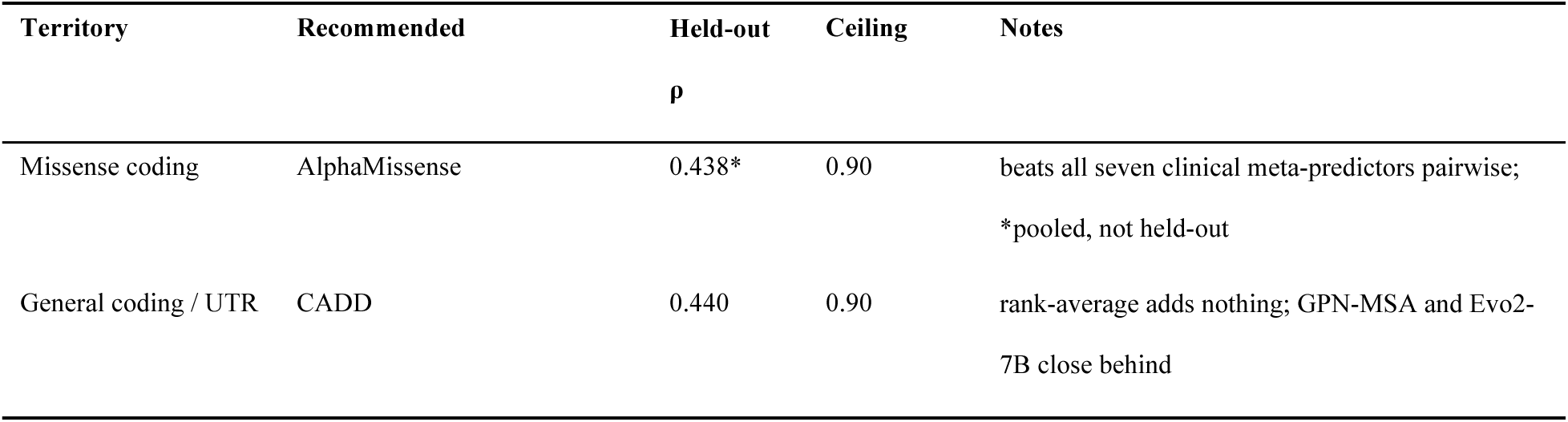
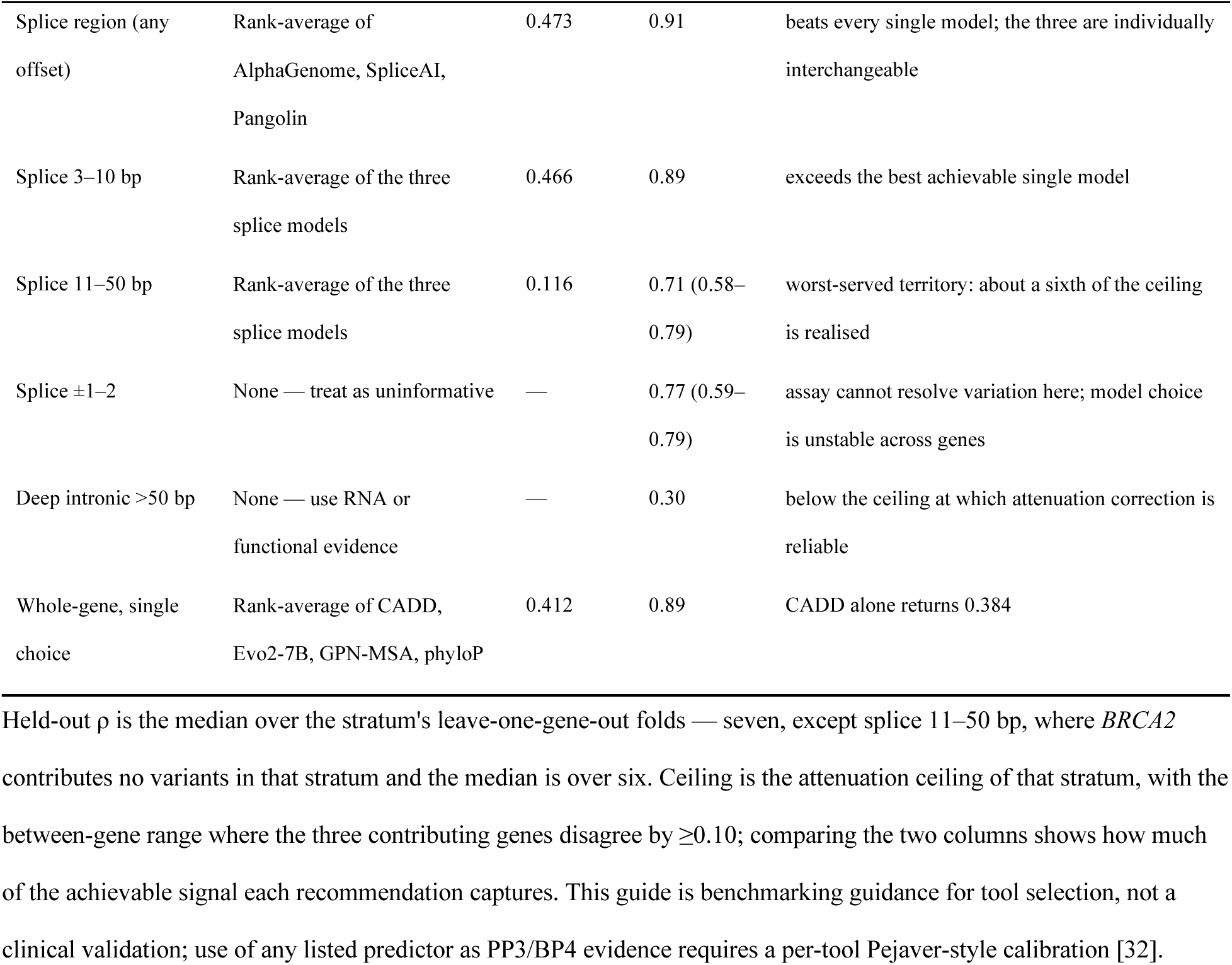
Territory-appropriate predictor selection guide. Recommendations follow the leave-one-gene-out comparison, so the quoted performance is what a user applying the rule to a new gene could expect, not a value fitted on the same data. Held-out ρ is the median over the stratum’s leave-one-gene-out folds — seven, except splice 11–50 bp, where *BRCA2* contributes no variants in that stratum and the median is over six. Ceiling is the attenuation ceiling of that stratum, with the between-gene range where the three contributing genes disagree by ≥0.10; comparing the two columns shows how much of the achievable signal each recommendation captures. This guide is benchmarking guidance for tool selection, not a clinical validation; use of any listed predictor as PP3/BP4 evidence requires a per-tool Pejaver-style calibration [32].

### Score concordance does not predict whether a definition change matters

Benchmarks pin scoring definitions and verify them by concordance. This atlas provides a direct test of whether that verification means anything: our AlphaGenome column and the frozen column of the companion calibration benchmark [52] agree at only ρ = 0.672 on 21,394 shared variants (the companion’s frozen analysis set, which is also this atlas’s ClinVar-recorded stratum; Methods), which read alone implies the definition choice is consequential. It is not. Replicating the legacy definition under the current client gives ρ = 0.995, so the disagreement is definitional rather than a version artefact; and evaluating both published definitions on the same variants shifts pooled ρ by at most 0.047 in any stratum.

A systematic sweep of four windows crossed with four aggregations confirms it, the largest spread in any stratum being 0.109. Score concordance and conclusion stability are different quantities and here they are decoupled, so a concordance gate belongs on a benchmark’s endpoints rather than on the correlation between score columns. The sweep, the version control and the full grid are given in Additional file 2: Note S11, Additional file 2: Figure S16 and Additional file 1: Table S8.

## Discussion

The single change that most alters what this benchmark says is also the cheapest: reporting, alongside each correlation, the correlation the assay’s own precision permits. That assay reproducibility limits achievable correlation is not new — Livesey and Marsh [22, 23] checked it directly when assessing deep mutational scanning as a benchmarking substrate — but estimating the bound per functional territory, correcting for it, and establishing where the correction is trustworthy has not been done, and the cost of not doing it is not cosmetic. It converts a measurement limit at canonical splice sites into an apparent model failure, it hides the territory where models are genuinely weakest, and it invites a strong negative conclusion about deep-intronic prediction from a stratum whose ceiling is 0.30. Our simulation gives the correction an operating envelope rather than a blanket endorsement: it reduces error above a ceiling of about 0.45 and amplifies it below, while remaining close to unbiased on both sides of the boundary, and should not be applied below that.

The definition sweep supports the same discipline from the other side. Two score columns of one predictor that agree at only ρ = 0.672 shift no stratum’s result by more than 0.047, so a concordance gate should be evaluated on a benchmark’s endpoints rather than on the correlation between score columns (Additional file 2: Note S11).

Two further methodological points generalise beyond this dataset. Ordering claims between predictors evaluated on shared variants must be tested pairwise; marginal intervals answer a different question and, as Figure 4 shows, mislead in both directions — they concealed a resolvable gap between Evo2-7B and CADD and the reliable ordering of nine missense predictors, while lending unwarranted support to AlphaGenome’s lead at 3–10 bp. And the ClinVar-recorded contrast must be drawn against variants ClinVar has not recorded; drawing it against the full set understated the ascertainment effect here by a factor of 2.5.

A benchmark against measured function is not automatically independent of the predictors it scores. None of the nineteen was trained on the seven saturation editing datasets used here, which post-date most of them, so there is no direct leakage. The dependency that does exist is indirect and runs through clinical labels: five of the seven dbNSFP meta-predictors — REVEL, BayesDel, ClinPred, MetaRNN and VEST4 — are trained or thresholded on ClinVar or HGMD, and a third of the atlas (21,394 of 64,178) carries a ClinVar record. That is why the ClinVar-recorded contrast above is not only a statement about benchmark optimism but also a bound on this dependency: a median uplift of 0.082 on recorded variants, and 0.445 against 0.312 for CADD.

The remaining predictors carry no label dependency of this kind. PrimateAI is trained on common primate variation and CADD on simulated versus fixed derived alleles, neither of which is a clinical label set; Evo2-7B, NT-v2-500M, GPN-MSA and ESM-1b are self-supervised on sequence or alignments with no variant labels at all; and the conservation tracks are alignment statistics. Read with that split in mind, the ordering within the supervised meta-predictors is the part most exposed to label overlap, and the splice-aware and self-supervised comparisons the least. The training signal behind each of the nineteen, whether it carries clinical labels, and whether any overlap with MAVE or SGE measurements is documented, are tabulated with a source for every row in Additional file 1: Table S11.

The cross-readout comparison reframes what a correlation of 0.4 against a functional assay means. Two readouts of the same experiment, in the same laboratory, on the same variants, agree at 0.251; several predictors exceed that against the fitness readout. Performance that looks modest against an implicit ceiling of 1.0 is close to the reproducibility of the phenomenon being measured. The same comparison exposes a blind spot the fitness axis conceals entirely: on the RNA axis the ranking inverts, and the only models that do better against RNA abundance than against fitness are splice-aware ones, corroborated independently by the synonymous stratum. For a laboratory triaging coding variants for possible splice-altering effects, the implication is concrete — the splice-aware models are the ones to reach for — for prioritisation, not as clinical evidence — even though they rank seventh, eighth and ninth of the twelve in coding sequence overall.

Three claims this benchmark would otherwise have licensed did not survive the checks the Methods make routine: that predictors fail at canonical splice sites, which the attenuation estimate turned into a limit of the assay; that AlphaGenome leads at 3–10 bp, which the paired test turned into a difference that cannot be resolved; and that a score-definition change agreeing at ρ = 0.672 must matter, which the systematic sweep turned into a shift of at most 0.047 in any stratum. We report them because they are evidence for the argument this paper makes: the checks that caught our own overstatements are the same ones a benchmark needs in order to catch anyone else’s.

Several limitations bound these conclusions. Every estimate reported here is computed within the same seven genes; no gene was held out as an independent test set, so the atlas measures agreement on these assays rather than generalisation beyond them. Seven genes, all DNA-damage-response or tumour-suppressor loci, cannot represent the genome, and per-gene heterogeneity is high (I² up to 99%).

The attenuation ceilings rest on three genes, since only *BRCA1*, *BARD1* and *PALB2* publish a usable error model; those three disagree by more than 0.20 at splice ±1–2 and at 11–50 bp, so realisations in exactly the two strata that carry the reinterpretation are ranges rather than point estimates. *BARD1* matters twice over: it is the sole source of the deep-intronic ceiling, and it is the median gene at ±1–2, so the frontier parity reported there does not survive its removal (1.41 rather than 1.02), although the median convergence survives that removal; at 1.43 the median gap narrows rather than closes. It does not survive every removal, though: dropping *BRCA1* inverts the median convergence (0.78; Results), so the convergence is robust to dropping either SE-based gene but not to dropping *BRCA1*. The SE-based estimator is optimistic by about 0.05 relative to the replicate-based one, and the deep-intronic ceiling comes from a single gene and falls below the boundary our simulation identifies. The *BARD1* deposit that carries both the frontier parity and the deep-intronic ceiling is a preprint [40], so the least mature deposit is also the most load-bearing.

The classification and evidence-strength analyses use assay-derived labels in three genes and are not a ClinGen calibration. The RNA readout covers coding variants only. The definition sweep covers one predictor; whether score concordance and conclusion stability are as decoupled for others is untested. Saturation editing scores are assay-specific constructs, not patient phenotypes. Evo2-7B used an 8,192-bp window for throughput and alternatives were not swept, which by this paper’s own argument is a caveat on its numbers. The seven dbNSFP meta-predictor columns were fetched without the dbNSFP release being recorded (Additional file 1: Table S9), so the effect of version drift in those columns is untested. Finally, this manuscript and its companion calibration benchmark [52] analyse the same seven genes — all loss-of-function, homologous-recombination or cell-survival assays, with *BRCA1* and *BRCA2* both measured in HAP1 cells — so the two share their source data and neither constitutes independent evidence for the other’s conclusions.

The gene set bounds these conclusions in a second way. All seven are tumour-suppressor genes whose disease mechanism is loss of function — five in homologous recombination, together with *VHL* and *BAP1* — because that is where saturation editing has been applied at this density, its cell-survival readout tracking the clinical phenotype closely. Channelopathies, metabolic disorders, developmental syndromes and gain-of-function mechanisms are unrepresented, and the MaveDB survey above places these seven inside a human deposit set that is itself concentrated on a few well-studied genes. The mechanism is not what the gene set bounds: reproducibility limits any correlation against any measurement, and that argument carries to any gene and any readout. What does not carry is the numerical map. Reliability ceilings are properties of particular experiments, so which territory is worst-served, and by how much, must be re-estimated wherever the assay changes. The deep-intronic result shows the difference: it rests on 683 variants in two genes whose ceiling is about 0.30, which is a statement about those two experiments rather than about deep-intronic variants at large.

Territory-resolved evaluation against experimental measurements should nonetheless become the default, and it carries one prescription that costs almost nothing and changes what a benchmark can honestly claim: report a per-stratum reliability estimate alongside every reported correlation, or state that the assay does not permit one. The estimate is computed from data the assay authors have already deposited, so the cost falls on reporting rather than on new experiments, and for most of the resource it is already possible.

The obstacle is not deposition but use: assay authors have largely done the work, and benchmarks built on their tables discard it, reporting a correlation against a measurement whose precision is published one column away. What survives that computation is measured rather than extrapolated: of 728 human deposits carrying the necessary columns, 674 yield a usable ceiling, 30 return a reliability at or below zero because the error column is not on the scale of the score, 14 an implausibly high one, and 10 too few rows. Where a deposit cannot support the estimate, saying so is itself informative: it marks the territories in which a benchmark’s numbers cannot be interpreted as statements about predictors at all.

A published error and a reproducibility estimate are also not interchangeable, and the size — even the sign — of the discrepancy depends on what the published error measures. Across 455 targets of the designed-stability platform, where trypsin and chymotrypsin provide an independent repeat measurement of the same variants, the fitting interval implies a higher ceiling than the repeat supports in all 450 targets where both could be computed, by a median of 0.019. Among the eight deposits publishing both replicates and a standard error the direction is not consistent. Two proteases are not technical replicates — their disagreement carries real methodological difference as well as noise — so the ceiling the repeat supports, 0.979, is a lower bound rather than an estimate, and the truth lies between it and the 0.999 the published interval implies. That platform is genuinely precise, more so than any human deposit here; it is simply not as precise as its interval states. We use the same device elsewhere, bounding reliability from *BAP1*’s nested time points and *VHL*’s second selection condition, neither of which is a clean replicate either. An error column cannot be used as a reliability estimate without knowing what it measures.

## Conclusions

A correlation against a functional assay cannot exceed what that assay’s own precision permits, and because precision varies across the territories a benchmark compares, the uncorrected map misplaces where prediction fails. Correcting for it moves the failure from the canonical splice site, where the assay is the binding constraint, to the 11–50 bp intronic window, where the predictors are. The estimate that makes this possible is cheap, and it generalises.

The reliability estimator used here is not specific to this atlas. It needs only what a deposit already publishes, and the survey above found that most of MaveDB already carries it, so anyone benchmarking a predictor against a MAVE can compute the ceiling before computing the correlation, today, without asking anything of the depositor. We would expect the practice to matter most where it is currently absent: in benchmarks that pool across deposits, where correlations with materially different upper bounds are presently averaged as though comparable.

A second use follows from the fact that functional scores are increasingly read as clinical evidence. Under ACMG/AMP criteria, PS3 and BS3 evidence strength is derived from the same assay measurements, calibrated to likelihood ratios, and multiplexed functional data have already resolved variants of uncertain significance at scale under that framework [53]. Whatever bounds the correlation also bounds the evidence, and a per-stratum ceiling is directly convertible into a statement about how much evidence a given assay can supply in a given territory. In our view — and this is an opinion, offered for discussion rather than demonstrated here — calibration frameworks would be strengthened by carrying an explicit reliability floor alongside the likelihood ratios they publish, so that a strength assignment cannot outrun the measurement it rests on.

For depositors, the survey points to a cheap intervention. The information a reliability estimate requires is usually present but rarely findable: a conventional column-name search recovers about a tenth of the deposits that in fact carry it, because error columns appear variously as ‘sè, ‘sigmà, ‘sd’, ‘fitting_err’, ‘score_95CI_low’ and a long tail of local conventions. A recommended column name, or a metadata field recording which column carries the error and on what scale, would make the resource-wide estimate routine. This costs depositors nothing they are not already doing.

For predictor development, the corrected map identifies where effort is likely to pay. The 11–50 bp intronic window is the territory where models genuinely underperform rather than merely appear to, and it is well measured enough that improvement would be visible. The deep intron is the opposite case: performance there is low, but the ceiling is lower still, so the limiting factor is the assay rather than the predictor. We would read that as an argument for assay development rather than model development in that window — a distinction the uncorrected map cannot make, and one that applies equally to any territory where a benchmark reports poor performance without reporting what performance was achievable.

## Methods

### Assays and the frozen atlas

Seven saturation genome editing assays were selected from MaveDB [20, 21] as the complete set of human cancer-susceptibility-gene experiments meeting our coverage criteria (saturation genome editing of an endogenous locus, with variant counts populating the coding/UTR stratum and at least the ±1–2 and 3–10 bp splice strata under our classifier; URNs in Table 1), fetched by script and registered with SHA-256 manifests at the point of download, so the freeze is auditable against its sources. Variants were mapped to GRCh38 via the Mutalyzer 3 reference-model API [33, 34] ([33] describes Mutalyzer 2; the API used here is Mutalyzer 3, queried 25 July 2026) using each assay’s MANE-Select transcript, with genomic anchors from Ensembl REST [54] and cross-validation by Ensembl VEP [55]; all 64,178 rows mapped. Per-assay scores were oriented so that larger values indicate more damaging function. Variants were classified from HGVS-c into coding/UTR and splice-offset bins by a single deterministic classifier. An earlier form of that classifier’s pattern accepted only a ‘*’ prefix before the position and so filed nine *BRCA1* variants lying in an intron within the 5’UTR as coding; the pattern now accepts UTR-relative positions. The correction moves only the ±1–2 stratum materially (largest change in any pooled value 0.036; 81 of 89 model × stratum cells move by less than 0.002) and is covered by a regression test. The *BRCA2* deposit publishes a score column only, with no auxiliary measurement columns, so it contributes no reliability or secondary-readout information (Additional file 2: Note S4).

Comparing a corrected value in one stratum with a corrected value in another requires a matched predictor set, since a stratum’s median otherwise summarises whatever predictors happen to cover it. The seven dbNSFP meta-predictors are predominantly missense — only PrimateAI is strictly missense-only, and the other six carry some nonsense, synonymous or splice-region coverage, substantial only for BayesDel (10.0% of its scored variants) and VEST4 (6.7%) and below 1.5% for the rest, tabulated per predictor in Additional file 1: Table S9 — and were excluded. Six of them have no ±1–2 estimate at all, their splice coverage falling below the thirty-variant threshold in every gene; BayesDel is the exception, scoring 850 ±1–2 variants across six genes, but it takes a single value across all 192 of them in *BAP1*, so its per-gene correlation there is undefined, and the pipeline propagates that to the pooled estimate rather than dropping the gene, uniformly across the grid. Of the remaining twelve, AlphaMissense and the two gnomAD allele-frequency columns carry no ±1–2 estimate, leaving nine. Observed ρ and realisation for the coding/UTR-versus-±1–2 comparison are medians over the three genes with a validated error model, taken from the same rows of the attenuation table (’atlas.matched_realisation’; Additional file 2: Note S13). The unmatched comparison is the more favourable one here — the two allele-frequency columns depress the coding median — so the matched figures are reported instead. We say the corrected convergence survives dropping a gene when the corrected median ratio remains below the uncorrected full-set ratio of 1.69 and keeps its direction (coding/UTR above ±1–2); by that criterion it survives dropping *BARD1* or *PALB2* and does not survive dropping *BRCA1* (Results; Additional file 1: Table S13).

### Protein consequence

No deposit populates hgvs_pro, so consequence was re-derived from the cached Mutalyzer transcript models, which carry the spliced transcript sequence and coding sequence (CDS) offsets. For each coding SNV the c. position is placed in its codon and both codons translated; the reference base implied by the HGVS string is checked against the transcript first and any mismatch recorded rather than classified. All 64,178 variants were classified with no reference mismatches, and the derived missense set is identical to the set AlphaMissense scores.

### Model scoring

Each predictor has one directory containing a pinned environment, a score definition and reference-file provenance. AlphaGenome was scored via the official API (client 0.7.0) using the merged-quantile splice score over a 16-kb window. The companion benchmark froze an AlphaGenome column defined under client 0.6.1, a different score definition: the two columns agree at ρ = 0.672 and are not interchangeable between the two manuscripts (Additional file 2: Note S11). CADD GRCh38-v1.7 precomputed scores were retrieved by remote tabix. AlphaMissense (hg38) [56] applies to missense SNVs only. GPN-MSA used the authors’ precomputed scores, sign-flipped. phyloP-and phastCons-100way came from UCSC Genome Browser track APIs [57]. gnomAD v4 global and popmax allele frequencies [58] were fetched from the gnomAD v4.1 GraphQL API [59] and sign-flipped. Nucleotide Transformer v2-500M-multi-species was scored as a masked 6-mer log-likelihood ratio over a 6,000-bp window and Evo2-7B as LL(REF) − LL(ALT) over an 8,192-bp window, both on one NVIDIA A6000, SNVs only. Seven dbNSFP meta-predictors — REVEL, BayesDel addAF [60], ClinPred, MetaRNN, PrimateAI [61], VEST4 [7], ESM-1b [11] — were taken from dbNSFP [62] via the MyVariant.info API [63] against GRCh38 in batches. Where dbNSFP returns one value per affected transcript they were aggregated to the most damaging, matching how CADD is aggregated over consequence records. ESM-1b was sign-flipped because it reports a log-likelihood ratio.

SpliceAI 1.3.1 and Pangolin (git 5cf94b8) were run at distance 50 with their bundled settings and then re-scored at full floating-point precision. Both tools compute their delta scores as floats and round only when formatting output, which placed 47% and 54% of coding variants and 67% and 71% of deep-intronic variants on a single value. The re-scores take each package’s own scoring function verbatim and change only those format expressions, so the score definition is unaltered; a validation gate requires the full-precision output, re-rounded to two decimals, to reproduce the released column exactly, which it does for every variant. Distinct values rise from 101 to 46,130 (SpliceAI) and 92 to 46,237 (Pangolin), and the tie-imposed bound on Spearman ρ rises to 1.000 in every stratum. Pooled ρ changes by at most 0.010 in six of seven strata; in the seventh, the deep intron — where the tie ceiling was lowest — full precision lowers the pooled estimate by 0.075 for SpliceAI and 0.065 for Pangolin. All analyses use the full-precision columns; the released rounded columns are retained alongside them. Re-scoring at the pinned Pangolin commit (5cf94b8) also reproduces the companion benchmark’s frozen Pangolin column at ρ = 1.0000 (n = 21,394), so the pin is traceable across both manuscripts.

### Evaluation and stratification

Functional scores are not comparable between assays — per-gene standard deviations span 0.062 to 6.84 (Fig. 1B) — and are far from normal. All evaluation is therefore by rank correlation computed within gene: Spearman ρ rather than Pearson, which would be dominated by the largest-scale assay, and rather than a threshold measure such as AUROC, which would discard the graded information the assays provide. The same non-normality is why the attenuation correction, whose classical form assumes Pearson correlation, is validated by simulation rather than assumed (below). For each predictor and stratum we computed the per-gene Spearman ρ against the oriented functional score, requiring at least 30 scored variants per gene, and pooled by DerSimonian–Laird random-effects meta-analysis [48] on Fisher-z transforms. Beyond the eight original strata we added ClinVar-absent, SNV, indel, UTR, non-truncating coding SNV and the protein-consequence classes (missense, synonymous, nonsense), sixteen in all. ClinVar-recorded variants are those in the companion benchmark’s frozen analysis set [52], drawn from the ClinVar GRCh38 release of 15 June 2026 [16] with records of every review status kept (selection rules in the companion’s Methods), and ClinVar-absent is the complement within the atlas. The primary analysis is unadjusted for multiplicity; Benjamini–Hochberg adjusted values across the full model × stratum grid and across the pairwise comparisons are provided (Additional file 1: Table S10, Additional file 1: Table S10b). Adjustment leaves every ordering claim in this paper intact: of 180 grid cells significant at *P* < 0.05, 173 remain at *q* < 0.05, the seven exceptions all being marginal gnomAD allele-fre*q*uency or phyloP cells; all twelve significant pairwise comparisons survive, including Evo2-7B versus CADD (*q* = 0.043) and all eight AlphaMissense comparisons (q ≤ 0.003). Every statistic was read programmatically from pipeline output files and every figure and table generated by a committed script. Two boundary choices differ from the companion benchmark [52], so strata are not directly comparable between the two manuscripts: we stratify intronic offset into 3–10, 11–50 and >50 bp windows where the companion uses a proximal window of |offset| ≤ 8 (a splice core of |offset| ≤ 2 with a 3–8 bp region band); seven exon-side last-base variants counted as coding/UTR here are counted as core splice there; the three UTR-intron variants at offset −3 fall in this atlas’s 3–10 bp band but carry no usable intron offset in the companion’s frozen matrix; and the companion excludes indels (17,786 included here). The companion’s analysis set of 21,410 SNVs overlaps this atlas’s SNV matrix in 21,394; the sixteen the atlas does not carry are *RAD51C* SNVs absent from the *RAD51C* deposit used here, a row-selection difference between the two frozen products.

### Measurement reliability and the attenuation ceiling

A correlation against a noisy measurement is bounded above by the square root of that measurement’s reliability [27]. Reliability was estimated per gene and per stratum by two estimators. Where a deposit publishes replicate scores whose mean is the deposited score (*BRCA1*, verified to floating-point equality), the Spearman correlation between replicates gives the reliability of one replicate and Spearman–Brown steps it up to the two-replicate mean. Where a deposit publishes per-variant standard errors (*BARD1*, *PALB2*), reliability is 1 − mean(SE²)/var(score); those SEs were accepted only after checking that (CI upper − score)/SE equals 1.96 for every row against the deposit’s own published intervals. *BAP1*’s standard errors have no published CI to validate against and *RAD51C*’s are on a different scale from its score, so both were excluded from the primary estimate. Two deposits carry repeated measurements that are not clean replicates — *BAP1* nested time points and *VHL* a second selection condition — and these two are reported separately as bounds in known directions. *BRCA1* carries both estimators and is the cross-check: the SE-based estimator gives ceilings 0.049–0.062 higher (per-stratum values in Additional file 1: Table S14). Realisation is the observed ρ divided by the ceiling, shown only where at least two genes contribute an error model; where those genes disagree by ≥0.10 the ceiling is reported as a range and the realisation inherits it.

### Survey of error models in MaveDB

The prescription this paper makes is only actionable to the extent that deposits publish what a reliability estimate needs, so we measured that across MaveDB rather than generalising from our seven assays. Every published score set was enumerated through the public API and its table retrieved. The search endpoint returns at most one hundred records and accepts no offset, so enumeration partitioned on target gene name — largest bucket forty-two — de-duplicating by URN; all 2,803 were reached. Seventeen score tables (0.6%) could not be retrieved and are counted as not permitting an estimate, which is the conservative direction. A deposit counts as permitting an estimate if its table carries either replicate columns some subset of which has a mean reproducing the deposited score across the table, or a per-variant standard error, standard deviation or confidence interval. The subset search matters: *BRCA1* carries two fitness and two RNA replicates, so testing only the full set would reject the very deposit whose reliability this paper estimates from such a pair. Deposits no subset reconciles are counted separately. Error columns are accepted on column name alone, without the CI/SE reconciliation our primary analysis applies, so these counts are upper bounds on what is usable. The survey is a committed script (’atlas.mavedb_survey’); its summary statistics are Additional file 1: Table S12, Note S14 in Additional file 2 gives the classification, and the per-deposit table is in the archived release. The classification was audited in both directions by inspecting full score-table headers: the sixty sampled deposits classified as carrying nothing were adjudicated, and the shortfall was vocabulary rather than content; a symmetric sample of twenty-five deposits classified as carrying errors has been drawn and its adjudication is in progress, so no false-positive rate is claimed. The sampled rows are archived (mavedb_survey_audit_v1.tsv).

Ceilings were computed for every surveyed deposit that permits one, reusing the estimators above. Reliability was not clipped: 1 - mean(SE^2)/var can only leave the unit interval from below, and does so when the error column is not on the scale of the score, which is a property of the deposit worth counting rather than an error to suppress. Deposits were stratified by reference class, and one platform is reported separately and never pooled: the protease-resistance stability data on de novo designed mini-proteins, identified from target names and titles, whose targets are not human variants and whose published interval is a curve-fitting confidence interval on a thermodynamic parameter rather than a reproducibility interval. Where a deposit publishes both replicates and an error column, the two estimators were compared directly; *BRCA1*, whose error model the primary analysis derives from its replicates, reproduces the 0.049–0.062 optimism reported above, which checks this computation against the main pipeline.

### Simulation validation of the attenuation correction

The classical attenuation result assumes Pearson correlation with normal error, whereas we apply it to Spearman ρ on assay scores that are far from normal and, at canonical splice sites, severely compressed. We therefore validated the whole chain, estimator included, by simulation. True scores are drawn by resampling the observed score distribution of one gene within one stratum — per gene, because the seven assays report on incompatible scales (per-gene standard deviations span 0.062 to 6.84) and a pooled marginal would calibrate the injected noise against the scale differences rather than the within-assay spread. A predictor is coupled to the true scores at a specified Spearman ρ through a Gaussian copula, using r = 2 sin(πρ/6) for the latent Pearson correlation. Two replicates are formed by adding independent Gaussian error whose standard deviation is set to achieve a target reliability of the two-replicate mean, and the mean is taken as the deposited score. Reliability is then estimated rather than taken as known — via Spearman–Brown for the replicate-based branch; the SE-based branch is validated separately in the extended grid. The grid spans five stratum shapes × six true ρ × six reliabilities, with 150 trials per gene in each cell. Between two and seven genes contribute depending on the stratum, so a cell carries 300 to 1,050 trials and the grid 151,200 in total (seed 20260804). An extended grid (ten reliability levels × three noise models × both estimators; 756,000 trials; seed 20260804) adds bounded-truncation noise, score-dependent noise calibrated on the *BRCA1*-derived SEs, and the SE-based estimation branch; its crossover results are reported in the Results and tabulated in Additional file 1: Table S15, with the original grid (151,200 trials) retained as Table S4.

### Tie structure, statistical power and paired comparisons

Where a score vector contains ties, Spearman ρ is bounded below 1 for arithmetic reasons. For each model × stratum cell we computed that bound exactly, as the correlation between the tied mid-ranks and an ideal target increasing strictly within every tie group; the bound is attained by construction. Minimum detectable ρ at 80% power and α = 0.05 was computed from the Fisher-z normal approximation. Ordering claims were tested on the variants both predictors score, since two correlations sharing an outcome variable are dependent: per gene we used Steiger’s test for dependent correlations [49], pooled the per-gene z differences by DerSimonian–Laird, and separately obtained a 95% interval for Δρ on the correlation scale by resampling genes (4,000 gene-cluster bootstrap replicates, seed 20260803). Steiger’s test was applied in its original 1980 form [49], whose variance formula assumes Pearson correlations; we did not apply the 1.060 variance correction proposed for Spearman correlations, which leaves the test slightly anti-conservative here, so every ordering claim carries the gene-cluster bootstrap P as a sensitivity analysis alongside the Steiger P — the two agree at α = 0.05 for all eighteen pairs tested (Additional file 1: Table S17).

### Classification and evidence strength

*BARD1* and *PALB2* publish a two-component posterior probability of abnormal function and *RAD51C* a four-class functional call. Posteriors were binarised at ≥0.9 and ≤0.1 with the ambiguous middle dropped; for *RAD51C* the two depleted classes were taken as abnormal and unchanged as normal, with enriched (n = 49) excluded, giving 31,835 labelled variants. AUROC and average precision were computed per gene and pooled on the logit scale by DerSimonian–Laird with Hanley–McNeil variances [51]; both metrics are implemented in-repository and verified against scikit-learn [64] to floating-point equality so the pinned environment does not need the dependency. The positive likelihood ratio at a target specificity is computed by scanning observed score values rather than taking a quantile of the negatives: with heavily quantised scores a quantile threshold can land on a large tie group, driving the false-positive rate to 1 and collapsing the likelihood ratio even for a perfectly separating score. Where quantisation admits no threshold at the target specificity the cell is reported as not evaluable rather than as weak. ACMG bands follow the Tavtigian point system [30] at a prior of 0.10 and are indicative only; a ClinGen-grade calibration would additionally require the prior-probability model and bootstrap procedure of Pejaver et al. [32], in the framework set out by Brnich et al. [31] and Richards et al. [29].

### Cross-readout replication, ensembles and the definition sweep

Four deposits publish an RNA-level score alongside the fitness score. Each was carried only after confirming it is positively rank-correlated with that assay’s own score, so the atlas orientation rule applies unchanged; *VHL*’s second RNA timepoint failed this check (ρ = −0.047) and was dropped automatically. Ensembles are the mean within-gene percentile rank over a panel fixed a priori, evaluated only on variants every panel member scores. Selection strategies were compared leave-one-gene-out: for each held-out gene the winning model per stratum was chosen using the other six genes only and then scored on the held-out gene. The AlphaGenome definition sweep scored 2,722 variants — sampled stratified by gene and region, oversampling splice strata — under four API-supported window widths (16,384; 131,072; 524,288; 1,048,576 bp). Each response carries both raw and quantile scores for all three splicing output types, so four aggregations (merged quantile, merged raw, max|raw|, max quantile) were derived per call and only the window cost extra calls: 10,888 calls in total, cached per variant × window, with no errors.

### Concordance validation

Where an independent prior run existed, every ported scorer had to reproduce it on shared variants before entering the atlas: CADD, both conservation scores, AlphaMissense, GPN-MSA, both gnomAD allele-frequency columns, SpliceAI and Pangolin passed at |ρ| = 1.000, with ρ = −1.000 counted as a pass for the three columns whose legacy orientation was opposite. NT-v2-500M was validated at ρ = 0.9997. AlphaGenome was deliberately re-defined and the two definitions agree at ρ = 0.672, analysed in the Results section. Concordance scatter plots for every ported column are given in Additional file 2: Figure S13.

### Use of large language models

Large language model assistance (Anthropic Claude, used via Claude Code and the Claude assistant, June–August 2026) was used to write analysis code and in the preparation of this manuscript. No data were generated by the model; all measurements are third-party public deposits cited by accession. All analysis code is committed to the repository [65] and archived with the release [66]; the pipeline is covered by 178 guardrail tests that run from a clean extract of the archived release with no network access, and by a 179th that checks the mapped reference bases against the Ensembl REST API and skips where that service is unreachable (test accounting in Additional file 2: Note S12), and every value reported in the text, tables and figures is read programmatically from pipeline output rather than transcribed. Figures and tables were verified against the archived release by content hash. The author designed the analyses, made all interpretive judgements, and is responsible for the accuracy of the reported results.

## Abbreviations

ACMG: American College of Medical Genetics and Genomics
AMP: Association for Molecular Pathology
API: application programming interface
AUROC: area under the receiver operating characteristic curve
CDS: coding sequence
CI: confidence interval
DOI: digital object identifier
GPU: graphics processing unit
GRCh38: Genome Reference Consortium Human Build 38
HGVS: Human Genome Variation Society
IQR: interquartile range
MANE: Matched Annotation from NCBI and EMBL-EBI
MAVE: multiplexed assay of variant effect
SE: standard error
SGE: saturation genome editing
SNV: single-nucleotide variant
TSV: tab-separated values
URN: Uniform Resource Name
UTR: untranslated region
VEP: Variant Effect Predictor

## Additional files

**Additional file 1: Supplementary Tables.** XLSX. Nineteen machine-readable tables (S1–S17): per-gene Spearman correlations (S1), extended strata (S2), per-gene and per-stratum reliability and attenuation estimates (S3), the simulation grid validating the correction (S4), the tie audit before and after full-precision re-scoring (S5, S5b), pairwise head-to-head comparisons (S6), the leave-one-gene-out selection comparison (S7), the AlphaGenome definition sweep (S8), the predictor inventory with per-column licences (S9), the Benjamini–Hochberg adjusted grids (S10, S10b), the training provenance of every scored predictor (S11), and the summary statistics quoted in the text with the output file each is computed from (S12), so every figure in the Results can be checked without leaving this package. Tables S13–S17 carry the robustness analyses: the leave-one-gene-out realisation folds (S13), the *BRCA1* estimator cross-check (S14), the extended simulation grid with its crossover analysis (S15), the MaveDB accounting and SE-convention sensitivity (S16), and the Steiger-versus-bootstrap comparison for all eighteen pairwise tests (S17). In S9 the version field is left empty for the seven dbNSFP meta-predictors, whose dbNSFP release was not recorded at fetch time; source and access date are as retrieved, and coverage_pct is n_scored as a percentage of the 64,178 atlas variants.

**Additional file 2: Supplementary Information.** PDF. Supplemental Notes S1–S14 and Supplementary Figures S1–S17.

## Declarations

### Ethics approval and consent to participate

Not applicable.

### Consent for publication

Not applicable.

### Availability of data and materials

The datasets generated and/or analysed during the current study are available in the Zenodo repository, https://doi.org/10.5281/zenodo.22751081 [66], which archives the frozen atlas, all nineteen score columns, the per-stratum reliability and attenuation estimates and the definition sweep. That is the version DOI for the release reported here, so the counts and outputs quoted in this paper are the ones it contains; the robustness analyses reported in Tables S13–S17 were produced by the same pipeline and are included in the archived release; the concept DOI https://doi.org/10.5281/zenodo.21828448 resolves to all versions. The saturation genome editing measurements analysed are third-party data, available from MaveDB under the accessions urn:mavedb:00000662-0-1 (*BAP1*), urn:mavedb:00001250-a-2 (*BARD1*), urn:mavedb:00000097-0-2 (*BRCA1*), urn:mavedb:00001225-a-1 (*BRCA2*), urn:mavedb:00001259-a-2 (*PALB2*), urn:mavedb:00000673-0-1 (*RAD51C*) and urn:mavedb:00000675-a-1 (*VHL*). This manuscript and its companion calibration benchmark share the same seven-gene saturation-genome-editing source data, derived predictor score columns, and variant-annotation code. The evidence-strength framework (95% specificity operating point, Tavtigian prior 0.10) was coordinated across both manuscripts and applied to different label sets; the companion reports the complete evidence-strength analysis, and the present manuscript reports the territory-stratified application.

All analysis code, the scoring harness, the figure-generating scripts and the guardrail test suite are in the project repository [65] and are archived with the Zenodo release. The project is platform independent and requires Python 3.12 or newer; environments are pinned exactly, with the resolved environment recorded separately from the direct dependencies, and raw and frozen data carry SHA-256 manifests. Running ‘make fetch && make test’ reproduces the pipeline and its 178 guardrail tests from the archived release, which includes the robustness analyses of Tables S13–S17; a 179th test verifies the mapped reference bases against the Ensembl REST API and therefore runs only where that service is reachable. Because the archived release ships the pipeline outputs, every figure and table in this paper regenerates in under a minute without re-scoring any predictor, and therefore without an AlphaGenome API key or a GPU; both are needed only to re-derive the scored columns from source.

Source code is released under the MIT licence. The frozen assay matrix is released under CC BY 4.0, and the seven MaveDB deposits carry CC0 1.0 or CC BY 4.0 terms that pass through. The score matrices additionally carry nineteen third-party predictor columns under mixed terms: eight are restricted to non-commercial use — AlphaGenome, CADD, AlphaMissense, NT-v2-500M, SpliceAI, REVEL, PrimateAI and VEST4, of which AlphaMissense and NT-v2-500M are also ShareAlike — and three carry no licence terms we were able to locate (BayesDel, ClinPred and MetaRNN). Per-column terms and the source consulted for each are listed in Additional file 1: Table S9 and released with the archive; each restriction travels with its column and with any figure or table derived from it.

Twenty-six machine-readable source files — all TSV except the Markdown mapping audit — giving the full per-gene, per-stratum numeric output behind the Supplemental Note are part of the Zenodo deposit rather than numbered supplemental items (Additional file 2: Note S13).

### Authors’ contributions

N.Z.: Conceptualization, Methodology, Software, Formal analysis, Data curation, Visualization, Writing – original draft, Writing – review & editing.

## Acknowledgements

GPU scoring was run on a rented cloud instance. I thank the MaveDB team and the authors of the seven saturation genome editing assays for making their functional measurements publicly available — including the replicate scores, standard errors and RNA-level readouts on which the central analysis of this paper depends, which are deposited but rarely used.

## References

1. Kircher M, Witten DM, Jain P, O’Roak BJ, Cooper GM, Shendure J. A general framework for estimating the relative pathogenicity of human genetic variants. Nat Genet. 2014;46:310–315. doi:10.1038/ng.2892.

2. Siepel A, Bejerano G, Pedersen JS, Hinrichs AS, Hou M, Rosenbloom K, et al. Evolutionarily conserved elements in vertebrate, insect, worm, and yeast genomes. Genome Res. 2005;15:1034–1050. doi:10.1101/gr.3715005.

3. Pollard KS, Hubisz MJ, Rosenbloom KR, Siepel A. Detection of nonneutral substitution rates on mammalian phylogenies. Genome Res. 2010;20:110–121. doi:10.1101/gr.097857.109.

4. Jaganathan K, Kyriazopoulou Panagiotopoulou S, McRae JF, Darbandi SF, Knowles D, Li YI, et al. Predicting Splicing from Primary Sequence with Deep Learning. Cell. 2019;176:535–548.e24. doi:10.1016/j.cell.2018.12.015.

5. Zeng T, Li YI. Predicting RNA splicing from DNA sequence using Pangolin. Genome Biol. 2022;23:103. doi:10.1186/s13059-022-02664-4.

6. Cheng J, Novati G, Pan J, Bycroft C, Žemgulytė A, Applebaum T, et al. Accurate proteome-wide missense variant effect prediction with AlphaMissense. Science. 2023;381:eadg7492. doi:10.1126/science.adg7492.

7. Carter H, Douville C, Stenson PD, Cooper DN, Karchin R. Identifying Mendelian disease genes with the Variant Effect Scoring Tool. BMC Genomics. 2013;14:S3. doi:10.1186/1471-2164-14-s3-s3.

8. Ioannidis NM, Rothstein JH, Pejaver V, Middha S, McDonnell SK, Baheti S, et al. REVEL: An Ensemble Method for Predicting the Pathogenicity of Rare Missense Variants. Am J Hum Genet. 2016;99:877–885. doi:10.1016/j.ajhg.2016.08.016.

9. Alirezaie N, Kernohan KD, Hartley T, Majewski J, Hocking TD. ClinPred: Prediction Tool to Identify Disease-Relevant Nonsynonymous Single-Nucleotide Variants. Am J Hum Genet. 2018;103:474–483. doi:10.1016/j.ajhg.2018.08.005.

10. Li C, Zhi D, Wang K, Liu X. MetaRNN: differentiating rare pathogenic and rare benign missense SNVs and InDels using deep learning. Genome Med. 2022;14:115. doi:10.1186/s13073-022-01120-z.

11. Brandes N, Goldman G, Wang CH, Ye CJ, Ntranos V. Genome-wide prediction of disease variant effects with a deep protein language model. Nat Genet. 2023;55:1512–1522. doi:10.1038/s41588-023-01465-0.

12. Benegas G, Albors C, Aw AJ, Ye C, Song YS. A DNA language model based on multispecies alignment predicts the effects of genome-wide variants. Nat Biotechnol. 2025;43:1960–1965. doi:10.1038/s41587-024-02511-w.

13. Dalla-Torre H, Gonzalez L, Mendoza-Revilla J, Lopez Carranza N, Grzywaczewski AH, Oteri F, et al. Nucleotide Transformer: building and evaluating robust foundation models for human genomics. Nat Methods. 2025;22:287–297. doi:10.1038/s41592-024-02523-z.

14. Brixi G, Durrant MG, Ku J, Naghipourfar M, Poli M, Sun G, et al. Genome modelling and design across all domains of life with Evo 2. Nature. 2026;652:1349–1361. doi:10.1038/s41586-026-10176-5.

15. Avsec Ž, Latysheva N, Cheng J, Novati G, Taylor KR, Ward T, et al. Advancing regulatory variant effect prediction with AlphaGenome. Nature. 2026;649:1206–1218. doi:10.1038/s41586-025-10014-0.

16. Landrum MJ, Lee JM, Benson M, Brown GR, Chao C, Chitipiralla S, et al. ClinVar: improving access to variant interpretations and supporting evidence. Nucleic Acids Res. 2018;46:D1062–D1067. doi:10.1093/nar/gkx1153.

17. Findlay GM, Boyle EA, Hause RJ, Klein JC, Shendure J. Saturation editing of genomic regions by multiplex homology-directed repair. Nature. 2014;513:120–123. doi:10.1038/nature13695.

18. Fowler DM, Fields S. Deep mutational scanning: a new style of protein science. Nat Methods. 2014;11:801–807. doi:10.1038/nmeth.3027.

19. Findlay GM, Daza RM, Martin B, Zhang MD, Leith AP, Gasperini M, et al. Accurate classification of *BRCA1* variants with saturation genome editing. Nature. 2018;562:217–222. doi:10.1038/s41586-018-0461-z.

20. Esposito D, Weile J, Shendure J, Starita LM, Papenfuss AT, Roth FP, et al. MaveDB: an open-source platform to distribute and interpret data from multiplexed assays of variant effect. Genome Biol. 2019;20:223. doi:10.1186/s13059-019-1845-6.

21. Rubin AF, Stone J, Bianchi AH, Capodanno BJ, Da EY, Dias M, et al. MaveDB 2024: a curated community database with over seven million variant effects from multiplexed functional assays. Genome Biol. 2025;26:13. doi:10.1186/s13059-025-03476-y.

22. Livesey BJ, Marsh JA. Using deep mutational scanning to benchmark variant effect predictors and identify disease mutations. Mol Syst Biol. 2020;16:e9380. doi:10.15252/msb.20199380.

23. Livesey BJ, Marsh JA. Updated benchmarking of variant effect predictors using deep mutational scanning. Mol Syst Biol. 2023;19:e11474. doi:10.15252/msb.202211474.

24. Notin P, Kollasch AW, Ritter D, van Niekerk L, Paul S, Spinner H, et al. ProteinGym: Large-Scale Benchmarks for Protein Fitness Prediction and Design. Adv Neural Inf Process Syst. 2023;36:64331–64379. doi:10.52202/075280-2810.

25. Smith C, Kitzman JO. Benchmarking splice variant prediction algorithms using massively parallel splicing assays. Genome Biol. 2023;24:294. doi:10.1186/s13059-023-03144-z.

26. Livesey BJ, Badonyi M, Dias M, Frazer J, Kumar S, Lindorff-Larsen K, et al. Guidelines for releasing a variant effect predictor. Genome Biol. 2025;26:97. doi:10.1186/s13059-025-03572-z.

27. Spearman C. Correlation calculated from faulty data. Br J Psychol. 1910;3:271–295. doi:10.1111/j.2044-8295.1910.tb00206.x.

28. Hutcheon JA, Chiolero A, Hanley JA. Random measurement error and regression dilution bias. BMJ. 2010;340:c2289. doi:10.1136/bmj.c2289.

29. Richards S, Aziz N, Bale S, Bick D, Das S, Gastier-Foster J, et al. Standards and guidelines for the interpretation of sequence variants: a joint consensus recommendation of the American College of Medical Genetics and Genomics and the Association for Molecular Pathology. Genet Med. 2015;17:405–424. doi:10.1038/gim.2015.30.

30. Tavtigian SV, Greenblatt MS, Harrison SM, Nussbaum RL, Prabhu SA, Boucher KM, et al. Modeling the ACMG/AMP variant classification guidelines as a Bayesian classification framework. Genet Med. 2018;20:1054–1060. doi:10.1038/gim.2017.210.

31. Brnich SE, Abou Tayoun AN, Couch FJ, Cutting GR, Greenblatt MS, Heinen CD, et al; ClinGen Sequence Variant Interpretation Working Group. Recommendations for application of the functional evidence PS3/BS3 criterion using the ACMG/AMP sequence variant interpretation framework. Genome Med. 2020;12:3. doi:10.1186/s13073-019-0690-2.

32. Pejaver V, Byrne AB, Feng BJ, Pagel KA, Mooney SD, Karchin R, et al. Calibration of computational tools for missense variant pathogenicity classification and ClinGen recommendations for PP3/BP4 criteria. Am J Hum Genet. 2022;109:2163–2177. doi:10.1016/j.ajhg.2022.10.013.

33. Lefter M, Vis JK, Vermaat M, den Dunnen JT, Taschner PEM, Laros JFJ. Mutalyzer 2: next generation HGVS nomenclature checker. Bioinformatics. 2021;37:2811–2817. doi:10.1093/bioinformatics/btab051.

34. Mutalyzer 3 reference-model API. Mutalyzer. https://mutalyzer.nl/api (accessed 27 August 2026).

35. Waters AJ, Brendler-Spaeth T, Smith D, Offord V, Tan HK, Zhao Y, et al. Saturation genome editing of *BAP1* functionally classifies somatic and germline variants. Nat Genet. 2024;56:1434–1445. doi:10.1038/s41588-024-01799-3.

36. Olvera-León R, Zhang F, Offord V, Zhao Y, Tan HK, Gupta P, et al. High-resolution functional mapping of *RAD51C* by saturation genome editing. Cell. 2024;187:5719–5734.e19. doi:10.1016/j.cell.2024.08.039.

37. Buckley M, Terwagne C, Ganner A, Cubitt L, Brewer R, Kim DK, et al. Saturation genome editing maps the functional spectrum of pathogenic *VHL* alleles. Nat Genet. 2024;56:1446–1455. doi:10.1038/s41588-024-01800-z.

38. Huang H, Hu C, Na J, Hart SN, Gnanaolivu RD, Abozaid M, et al; CARRIERS Consortium. Functional evaluation and clinical classification of *BRCA2* variants. Nature. 2025;638:528–537. doi:10.1038/s41586-024-08388-8.

39. Boonen RACM, Knaup SC, Menafra R, Braspenning ME, Rother MB, Kleiblova P, et al; CZECANCA consortium. Site-saturation functional screens identify *PALB2* missense variants associated with increased breast cancer risk. Nat Commun. 2026;17:775. doi:10.1038/s41467-025-67252-z.

40. Woo I, Casadei S, Snyder MW, Smith NT, Best S, Tejura M, et al. Saturation genome editing of *BARD1* resolves VUS and provides insight into *BRCA1*-*BARD1* tumor suppression. medRxiv. 2025. doi:10.1101/2025.11.03.25339440 (preprint).

41. Waters AJ, Brendler-Spaeth T, Smith D, Offord V, Tan HK, Zhao Y, et al. BAP1 SGE. MaveDB. 2026. urn:mavedb:00000662-0-1. https://mavedb.org/score-sets/urn:mavedb:00000662-0-1.

42. Woo I, Casadei S, Snyder MW, Smith NT, Best S, Tejura M, et al. Saturation Genome Editing of *BARD1* scores. MaveDB. 2026. urn:mavedb:00001250-a-2. https://mavedb.org/score-sets/urn:mavedb:00001250-a-2.

43. Findlay GM, Daza RM, Martin B, Zhang MD, Leith AP, Gasperini M, et al. *BRCA1* SGE Normalized Scores. MaveDB. 2025. urn:mavedb:00000097-0-2. https://mavedb.org/score-sets/urn:mavedb:00000097-0-2.

44. Huang H, Hu C, Na J, Hart SN, Gnanaolivu RD, Abozaid M, et al. Scores from sGE of *BRCA2* exons 15 to 26 in Hap1 cells. MaveDB. 2025. urn:mavedb:00001225-a-1. https://mavedb.org/score-sets/urn:mavedb:00001225-a-1.

45. Boonen RACM, Knaup SC, Menafra R, Braspenning ME, Rother MB, Kleiblova P, et al. Saturation Genome Editing of *PALB2* scores. MaveDB. 2026. urn:mavedb:00001259-a-2. https://mavedb.org/score-sets/urn:mavedb:00001259-a-2.

46. Olvera-León R, Zhang F, Offord V, Zhao Y, Tan HK, Gupta P, et al. *RAD51C* SGE. MaveDB. 2026. urn:mavedb:00000673-0-1. https://mavedb.org/score-sets/urn:mavedb:00000673-0-1.

47. Buckley M, Terwagne C, Ganner A, Cubitt L, Brewer R, Kim DK, et al. Complete *VHL* SGE data. MaveDB. 2024. urn:mavedb:00000675-a-1. https://mavedb.org/score-sets/urn:mavedb:00000675-a-1.

48. DerSimonian R, Laird N. Meta-analysis in clinical trials. Control Clin Trials. 1986;7:177–188. doi:10.1016/0197-2456(86)90046-2.

49. Steiger JH. Tests for comparing elements of a correlation matrix. Psychol Bull. 1980;87:245–251. doi:10.1037/0033-2909.87.2.245.

50. Livesey BJ, Marsh JA. Variant effect predictor correlation with functional assays is reflective of clinical classification performance. Genome Biol. 2025;26:104. doi:10.1186/s13059-025-03575-w.

51. Hanley JA, McNeil BJ. The meaning and use of the area under a receiver operating characteristic (ROC) curve. Radiology. 1982;143:29–36. doi:10.1148/radiology.143.1.7063747.

52. Zhang N. Ranking is saturated, calibration is not, and neither changes the evidence strength: an independent benchmark of splice-region variant effect predictors against a functional standard. Submitted.

53. Fayer S, Horton C, Dines JN, Rubin AF, Richardson ME, McGoldrick K, et al. Closing the gap: systematic integration of multiplexed functional data resolves variants of uncertain significance in *BRCA1*, *TP53*, and *PTEN*. Am J Hum Genet. 2021;108:2248–2258. doi:10.1016/j.ajhg.2021.11.001.

54. Yates A, Beal K, Keenan S, McLaren W, Pignatelli M, Ritchie GRS, et al. The Ensembl REST API: Ensembl Data for Any Language. Bioinformatics. 2015;31:143–145. doi:10.1093/bioinformatics/btu613.

55. McLaren W, Gil L, Hunt SE, Riat HS, Ritchie GRS, Thormann A, et al. The Ensembl Variant Effect Predictor. Genome Biol. 2016;17:122. doi:10.1186/s13059-016-0974-4.

56. Cheng J, Novati G, Pan J, Bycroft C, Žemgulytė A, Applebaum T, et al. Predictions for AlphaMissense. Zenodo. 2023. doi:10.5281/zenodo.8208688.

57. Kent WJ, Sugnet CW, Furey TS, Roskin KM, Pringle TH, Zahler AM, et al. The Human Genome Browser at UCSC. Genome Res. 2002;12:996–1006. doi:10.1101/gr.229102.

58. Chen S, Francioli LC, Goodrich JK, Collins RL, Kanai M, Wang Q, et al; Genome Aggregation Database Consortium. A genomic mutational constraint map using variation in 76,156 human genomes. Nature. 2024;625:92–100. doi:10.1038/s41586-023-06045-0.

59. gnomAD v4.1 GraphQL API. Broad Institute. https://gnomad.broadinstitute.org (accessed 27 August 2026).

60. Feng BJ. PERCH: A Unified Framework for Disease Gene Prioritization. Hum Mutat. 2017;38:243–251. doi:10.1002/humu.23158.

61. Sundaram L, Gao H, Padigepati SR, McRae JF, Li Y, Kosmicki JA, et al. Predicting the clinical impact of human mutation with deep neural networks. Nat Genet. 2018;50:1161–1170. doi:10.1038/s41588-018-0167-z.

62. Liu X, Li C, Mou C, Dong Y, Tu Y. dbNSFP v4: a comprehensive database of transcript-specific functional predictions and annotations for human nonsynonymous and splice-site SNVs. Genome Med. 2020;12:103. doi:10.1186/s13073-020-00803-9.

63. Xin J, Mark A, Afrasiabi C, Tsueng G, Juchler M, Gopal N, et al. High-performance web services for querying gene and variant annotation. Genome Biol. 2016;17:91. doi:10.1186/s13059-016-0953-9.

64. Pedregosa F, Varoquaux G, Gramfort A, Michel V, Thirion B, Grisel O, et al. Scikit-learn: Machine Learning in Python. J Mach Learn Res. 2011;12:2825–2830.

65. functional-standard-atlas source repository. GitHub. https://github.com/oneone00-11/functional-standard-atlas (accessed 27 August 2026).

66. Zhang N. functional-standard-atlas: attenuation-corrected, territory-resolved benchmarking of variant effect predictors against saturation genome editing. Version v2.5.0. Zenodo. 2026. doi:10.5281/zenodo.22751081.

