## Additional file 2: Supplementary Information for "Measurement reliability bounds functional benchmarks and relocates where variant effect prediction fails"

Every statistic below is read from a pipeline output file at build time by `atlas.supplement`; the source file is named in each section.

#### Contents

S1. Variant mapping and atlas freezing

S2. Concordance validation

S3. GPU scoring runs

S4. Auxiliary assay measurements and protein consequence

S5. Measurement reliability and the attenuation ceiling

S6. Score quantisation and statistical power

S7. Paired comparisons and leave-one-gene-out sensitivity

S8. Extended strata

S9. Classification, evidence strength, readouts and ensembles

S10. Simulation validation of the attenuation correction

S11. AlphaGenome definition sweep

S12. Repository and reproduction

S13. Machine-readable source files

S14. What MaveDB deposits permit

#### S1. Variant mapping and atlas freezing

All 64,178 assay rows (7 MaveDB SGE assays) were mapped to GRCh38 with a **100.00% success rate** (`results/mapping_report_v1.md`). Provenance was fetched live by script and hash-registered: MANE Select transcripts (NCBI MANE GRCh38 v1.5); transcript exon models (Mutalyzer3 API); genomic exon anchors and reference sequence (Ensembl REST, GRCh38); cross-validation by Ensembl VEP.

Orientation rule, recorded per assay: `functional_pathogenicity = −score` (SGE raw score: higher = more normal function).

Coordinates are GRCh38, `pos` 1-based, indels left-anchored VCF-style, alleles on the + strand.

##### S1.1 Region-classifier correction

An earlier form of the intronic-offset pattern accepted only a `*` prefix before the position, and therefore filed nine BRCA1 variants lying in an intron *within the 5′UTR* (`c.-19-1`, `c.-19-2`, `c.-19-3`) as coding rather than splice. The pattern now accepts UTR-relative positions. Six of the nine move into splice ±1–2 and three into splice 3–10 bp. Only the ±1–2 stratum changes materially: the largest change in any pooled ρ is 0.036 (phastCons at ±1–2, −0.003 → 0.033) and 81 of 89 model × stratum cells move by less than 0.002. `tests/test_hardening.py::test_classify_region` pins the corrected behaviour.

#### S2. Concordance validation

Every scorer ported from the companion benchmark had to reproduce the independent prior run on shared variants before entering the atlas (`results/concordance_*.json`). ρ = ±1.0 counts as a pass where the legacy column used the opposite orientation.

| Model | Concordance ρ | n shared | Verdict |
|---|---|---|---|
| CADD | 1.000 | 19,883 | pass |
| AlphaMissense | 1.000 | 11,911 | pass |
| phyloP-100way | 1.000 | 21,394 | pass |
| phastCons-100way | 1.000 | 21,394 | pass |
| SpliceAI | 1.000 | 21,394 | pass |
| Pangolin | 1.000 | 21,394 | pass |
| GPN-MSA | −1.000 | 21,394 | pass (orientation flip) |
| gnomAD AF (global) | −1.000 | 7,702 | pass (orientation flip) |
| gnomAD AF (popmax) | −1.000 | 7,702 | pass (orientation flip) |
| NT-v2-500M | 0.9997 (per-gene 0.9994–0.9998) | 21,394 | pass |
| AlphaGenome | **0.672** | 21,394 | **deliberate re-definition — analysed in main text** |
| Evo2-7B | — (no legacy column) | — | new standard definition |

#### S3. GPU scoring runs

Both foundation models were scored on a rented NVIDIA A6000 (48 GB) under tmux with append-only TSV score caches for crash resumability (environments pinned in `models/{nt,evo2}/requirements.txt` with full `pip freeze` archives).

**NT-v2-500M**: masked 6-mer LLR, 6,000-bp window, transformers 4.46.3 / torch 2.11.0+cu128; 46,392 SNVs.

**Evo2-7B**: windowed LL delta, 8,192-bp window, evo2 0.6.0 / torch 2.7.1+cu128 / flash-attn 2.8.0.post2, bf16; 46,392 SNVs in 25.0 h (5,024 batches, batch size 8).

Skipped (unscored): FASTA reference-base mismatch; ambiguous (N) bases in window; windows truncated below 1 kb at chromosome ends.

#### S4. Auxiliary assay measurements and protein consequence

Per-gene and pooled correlations for every model × stratum cell are in Supplemental_Table_S1; per-model forest plots are Supplemental_Fig_S1–S12 and the concordance scatters Supplemental_Fig_S13.

`results/assay_aux_v1.parquet`, `results/consequence_v1.parquet`

The frozen atlas keeps one number per variant. The deposits carry more; `atlas.assay_aux` lifts the additional columns into one table, validating each. Coverage within the atlas:

| Column | Variants |
|---|---|

| | |
|---|---|
| fit_rep1 | 3,893 |
| se | 54,951 |
| rna | 19,553 |
| rna_alt | 0 |
| rna_rep1 | 2,743 |
| label_abnormal | 32,901 |

Per-assay decisions:

**BRCA1** — fitness_reps=score_rep1,score_rep2; se=sd(reps)/sqrt(2) (derived_from_replicates); rna=score_rna (rho=+0.429); rna_reps=score_rna_rep1,score_rna_rep2

**BAP1** — se=SE_bind_continuous (unvalidated)

**RAD51C** — se=SE_bind_continuous (scale_mismatch); label_class=functional_classification

**VHL** — rna=rna_score_d6 (rho=+0.266); rna_alt=rna_score_d20 DROPPED (rho=-0.047 < 0.05)

**BRCA2** — score only, no auxiliary columns

**BARD1** — se=standard_error (validated); rna=rna_score (rho=+0.129); label_prob=gmm_density_abnormal

**PALB2** — se=standard_error (validated); rna=rna_score (rho=+0.166); label_prob=gmm_density_abnormal

Standard errors were accepted only where the deposit also publishes a 95% CI to check them against; BARD1 and PALB2 both give (CI upper − score)/SE = 1.960 exactly. BAP1's SE has no published CI and RAD51C's is on a different scale from its score, so neither is used for reliability. RNA columns were carried only if positively rank-correlated with that assay's own score, so the atlas orientation rule applies unchanged; VHL's second RNA timepoint failed this check ($\rho = -0.047$) and was dropped automatically.

`atlas.consequence` re-derives protein consequence from the cached Mutalyzer transcript models, because `hgvs_pro` is empty in all seven deposits. All 64,178 variants classified, no reference-base mismatches:

| Consequence | n |
|---|---|
| missense | 26,022 |
| indel_or_complex | 15,288 |
| intronic | 11,655 |
| synonymous | 7,886 |
| nonsense | 1,656 |
| utr3 | 823 |
| utr5 | 764 |
| start_lost | 46 |
| stop_lost | 38 |

As an external check the derived missense set (26,022) is identical, variant for variant, to the set AlphaMissense scores.

#### S5. Measurement reliability and the attenuation ceiling

`results/reliability_v1.tsv`, `results/attenuation_v1.tsv` (Supplemental_Table_S3)

The table below reports the **attenuation ceiling** (= √reliability) per gene × stratum, not the reliability itself. Reliability is estimated from replicate scores (BRCA1, Spearman–Brown on the replicate correlation; the deposited score is exactly the replicate mean) or from CI-validated per-variant standard errors (BARD1, PALB2; `1 − mean(SE²)/var(score)`). To recover a reliability, square the tabulated value: the coding/UTR median ceiling of 0.897 corresponds to a reliability of 0.805, and the splice ±1–2 median ceiling of 0.772 to a reliability of 0.596.

The between-gene spread is small in most strata but large at splice ±1–2 and splice 11–50 bp, where the three genes disagree by more than 0.20. Realisations in those two strata should be read as ranges, not point estimates; the main text quotes the median-ceiling value and states the range.

| Stratum | BARD1 | BRCA1 | PALB2 | Spread | Median |
|---|---|---|---|---|---|
| All | 0.938 | 0.894 | 0.894 | 0.044 | 0.894 |
| Coding / UTR | 0.940 | 0.897 | 0.889 | 0.052 | 0.897 |
| Splice region | 0.923 | 0.887 | 0.912 | 0.037 | 0.912 |
| Splice ±1–2 | 0.586 | 0.772 | 0.794 | 0.208 | 0.772 |
| Splice 3–10 bp | 0.928 | 0.881 | 0.887 | 0.048 | 0.887 |
| Splice 11–50 bp | 0.788 | 0.583 | 0.710 | 0.205 | 0.710 |
| Splice >50 bp | 0.297 | — | — | 0.000 | 0.297 |

**Cross-check.** BRCA1 carries both estimators. The SE-based estimator returns a ceiling 0.049–0.062 higher than the replicate-based one in every stratum:

| Stratum | Ceiling (replicates) | Ceiling (SE) | Difference |
|---|---|---|---|
| All | 0.894 | 0.945 | 0.050 |
| Coding / UTR | 0.897 | 0.946 | 0.049 |
| Splice 11–50 bp | 0.583 | 0.645 | 0.062 |
| Splice ±1–2 | 0.772 | 0.829 | 0.056 |
| Splice 3–10 bp | 0.881 | 0.938 | 0.057 |
| Splice region | 0.887 | 0.941 | 0.054 |

BARD1 and PALB2 ceilings are therefore, if anything, slight overestimates, which makes the realisations in the main text slightly conservative.

Realisation (observed ρ ÷ ceiling), median over the contributing genes, shown only where at least two genes contribute an error model:

| Predictor | All | Coding / UTR | Splice region | Splice ±1–2 | Splice 3–10 bp | Splice 11–50 bp | Splice >50 bp |
|---|---|---|---|---|---|---|---|
| AlphaGenome | 0.158 | 0.064 | 0.522 | 0.423 | 0.583 | 0.126 | — |
| SpliceAI | 0.151 | 0.108 | 0.474 | 0.163 | 0.586 | 0.152 | — |
| Pangolin | 0.150 | 0.107 | 0.552 | 0.289 | 0.567 | 0.156 | — |
| CADD | 0.400 | 0.429 | 0.417 | 0.095 | 0.322 | -0.023 | — |
| AlphaMissense | 0.367 | 0.367 | — | — | — | — | — |
| Evo2-7B | 0.389 | 0.368 | 0.443 | 0.096 | 0.471 | 0.126 | — |
| GPN-MSA | 0.339 | 0.339 | 0.398 | 0.173 | 0.339 | 0.002 | — |

| | | | | | | | |
|---|---|---|---|---|---|---|---|
| NT-v2-500M | 0.099 | 0.077 | 0.226 | 0.297 | 0.214 | 0.053 | — |
| phyloP-100way | 0.289 | 0.284 | 0.368 | 0.143 | 0.250 | 0.050 | — |
| phastCons-100way | 0.274 | 0.233 | 0.372 | 0.157 | 0.274 | 0.066 | — |
| gnomAD AF (global) | 0.073 | 0.060 | 0.136 | — | 0.051 | 0.137 | — |
| gnomAD AF (popmax) | 0.058 | 0.057 | 0.081 | — | 0.050 | 0.109 | — |
| REVEL | 0.380 | 0.379 | — | — | — | — | — |
| BayesDel (AF) | 0.511 | 0.483 | 0.041 | 0.048 | — | — | — |
| ClinPred | 0.350 | 0.350 | — | — | — | — | — |
| MetaRNN | 0.355 | 0.354 | — | — | — | — | — |
| PrimateAI | 0.257 | 0.257 | — | — | — | — | — |
| VEST4 | 0.453 | 0.452 | — | — | — | — | — |
| ESM-1b | 0.314 | 0.314 | — | — | — | — | — |

#### S6. Score quantisation and statistical power

`results/tie_audit_v1.tsv` (Supplemental_Table_S5; rounded vs full precision in Supplemental_Table_S5b), `results/power_v1.tsv`

Where a score vector contains ties, Spearman ρ is bounded below 1 for arithmetic reasons. The bound below is exact and attained by construction. The twelve most tie-limited cells:

| **Predictor** | **Stratum** | **n** | **Distinct values** | **Modal value** | **Modal share** | **ρ ceiling from ties** |
|---|---|---|---|---|---|---|
| phastCons-100way | Splice ±1–2 | 1056 | 32 | 1.000 | 0.876 | 0.572 |
| phastCons-100way | Splice >50 bp | 683 | 49 | 0.000 | 0.650 | 0.852 |
| phastCons-100way | Splice 11–50 bp | 5961 | 129 | 0.000 | 0.618 | 0.874 |
| gnomAD AF (popmax) | Splice >50 bp | 202 | 85 | -0.000 | 0.559 | 0.908 |
| gnomAD AF (global) | Splice >50 bp | 202 | 87 | 0.000 | 0.550 | 0.913 |
| phastCons-100way | Coding / UTR | 52523 | 128 | 1.000 | 0.490 | 0.935 |
| phastCons-100way | Splice region | 11655 | 174 | 0.000 | 0.465 | 0.946 |
| phastCons-100way | All | 64178 | 175 | 1.000 | 0.429 | 0.952 |
| BayesDel (AF) | Splice ±1–2 | 850 | 354 | 0.625 | 0.332 | 0.979 |
| phastCons-100way | Splice 3–10 bp | 3955 | 121 | 0.000 | 0.322 | 0.980 |
| BayesDel (AF) | Splice region | 955 | 457 | 0.625 | 0.295 | 0.985 |
| VEST4 | Splice region | 34 | 22 | 0.157 | 0.235 | 0.992 |

The SpliceAI and Pangolin command-line tools round delta scores to two decimals, which is the origin of most of this. It matters in one place — phastCons at ±1–2, capped at 0.572 — and does not explain the deep-intronic result, where the caps are still 0.833 (SpliceAI) and 0.804 (Pangolin).

Minimum ρ detectable at 80% power, α = 0.05, from the per-gene n actually used:

| Stratum | Genes | Smallest gene n | MDR, single gene | MDR, pooled |
|---|---|---|---|---|
| All | 7 | 748 | 0.102 | 0.027 |
| Coding / UTR | 7 | 557 | 0.118 | 0.031 |
| Splice region | 7 | 34 | 0.465 | 0.465 |
| Splice ±1–2 | 6 | 96 | 0.283 | 0.098 |
| Splice 3–10 bp | 7 | 76 | 0.317 | 0.109 |
| Splice 11–50 bp | 6 | 49 | 0.391 | 0.391 |
| Splice >50 bp | 2 | 74 | 0.321 | 0.197 |

The deep-intronic stratum can only detect ρ ≥ 0.107 pooled for the predictors that score it, which bounds how strongly a null result there can be stated. The 0.197 in the table above is the worst-covered column rather than a predictor: the two gnomAD allele-frequency columns carry 74 variants in their smallest gene against 249 to 296 for everything else.

#### S7. Paired comparisons and leave-one-gene-out sensitivity

`results/head_to_head_v1.tsv` (Supplemental_Table_S6), `results/logo_v1.tsv`

Two correlations measured on the same variants are dependent, so overlapping marginal CIs are not a test of their difference. Each ordering claim was tested on the variants both predictors score, with Steiger's test pooled across genes and a 4,000-replicate gene-cluster bootstrap for Δρ (seed 20260803).

| Model A | Model B | Stratum | n paired | ρ(A) | ρ(B) | Δρ | Bootstrap 95% CI | Steiger p |
|---|---|---|---|---|---|---|---|---|
| AlphaGenome | SpliceAI | Splice 3–10 bp | 3290 | 0.458 | 0.444 | 0.014 | -0.012 to 0.036 | 0.209 |
| AlphaGenome | Pangolin | Splice 3–10 bp | 3290 | 0.458 | 0.447 | 0.011 | -0.013 to 0.042 | 0.651 |
| Pangolin | SpliceAI | Splice 3–10 bp | 3290 | 0.447 | 0.444 | 0.002 | -0.040 to 0.036 | 0.624 |
| Pangolin | SpliceAI | Splice 11–50 bp | 4707 | 0.175 | 0.159 | 0.015 | -0.055 to 0.078 | 0.673 |
| Pangolin | AlphaGenome | Splice region | 9489 | 0.458 | 0.446 | 0.011 | -0.011 to 0.030 | 0.404 |
| Evo2-7B | CADD | All | 46392 | 0.358 | 0.391 | -0.034 | -0.069 to -0.006 | 0.028 |
| Evo2-7B | GPN-MSA | All | 46392 | 0.358 | 0.363 | -0.005 | -0.051 to 0.037 | 0.820 |
| CADD | GPN-MSA | Coding / UTR | 36903 | 0.427 | 0.378 | 0.049 | 0.021 to 0.077 | 0.000 |
| Evo2-7B | NT-v2-500M | All | 46392 | 0.358 | 0.098 | 0.259 | 0.191 to 0.342 | 0.000 |
| CADD | phyloP-100way | All | 46392 | 0.391 | 0.307 | 0.085 | 0.065 to 0.104 | 0.000 |
| AlphaMissense | MetaRNN | Missense | 26004 | 0.438 | 0.409 | 0.028 | 0.014 to 0.042 | 0.000 |
| AlphaMissense | ClinPred | Missense | 26022 | 0.438 | 0.397 | 0.040 | 0.026 to 0.054 | 0.000 |
| AlphaMissense | REVEL | Missense | 25866 | 0.436 | 0.397 | 0.040 | 0.018 to 0.053 | 0.001 |
| AlphaMissense | BayesDel | Missense | 26022 | 0.438 | 0.396 | 0.042 | 0.019 to | 0.002 |

| | | | | | | | | |
|---|---|---|---|---|---|---|---|---|
| | (AF) | | | | | | 0.063 | |
| AlphaMissense | VEST4 | Missense | 25909 | 0.437 | 0.394 | 0.043 | 0.025 to 0.063 | 0.000 |
| AlphaMissense | ESM-1b | Missense | 21289 | 0.447 | 0.344 | 0.103 | 0.026 to 0.215 | 0.001 |
| AlphaMissense | PrimateAI | Missense | 25818 | 0.437 | 0.272 | 0.165 | 0.101 to 0.230 | 0.000 |
| AlphaMissense | CADD | Missense | 26022 | 0.438 | 0.376 | 0.061 | 0.043 to 0.080 | 0.000 |

Leave-one-gene-out: the ten largest swings in pooled ρ when a single gene is dropped.

| **Predictor** | **Stratum** | **Pooled ρ** | **LOGO min** | **LOGO max** | **Most influential gene** | **Δ if dropped** |
|---|---|---|---|---|---|---|
| BayesDel (AF) | Splice region | 0.235 | 0.109 | 0.309 | BAP1 | -0.126 |
| Pangolin | Splice ±1–2 | 0.198 | 0.130 | 0.250 | BARD1 | -0.068 |
| phyloP-100way | Splice ±1–2 | 0.037 | -0.002 | 0.106 | RAD51C | 0.069 |
| GPN-MSA | Splice ±1–2 | 0.121 | 0.063 | 0.170 | BAP1 | -0.057 |
| NT-v2-500M | Splice ±1–2 | 0.101 | 0.047 | 0.140 | PALB2 | -0.054 |
| AlphaGenome | Splice ±1–2 | 0.158 | 0.109 | 0.197 | PALB2 | -0.049 |
| SpliceAI | Splice ±1–2 | 0.140 | 0.089 | 0.169 | BRCA2 | -0.051 |
| CADD | Splice 11–50 bp | 0.062 | 0.018 | 0.097 | VHL | -0.043 |
| ESM-1b | All | 0.345 | 0.308 | 0.384 | PALB2 | 0.039 |
| ESM-1b | Coding / UTR | 0.345 | 0.308 | 0.384 | PALB2 | 0.039 |

All of the largest swings are in the ±1–2 stratum. The headline coding and all-variant numbers are stable (e.g. AlphaMissense coding/UTR 0.438, range 0.410–0.475 across the seven leave-one-out folds).

#### S8. Extended strata

`results/eval_ext_v1.tsv` (Supplemental_Table_S2)

Pooled ρ over the original eight strata plus ClinVar-absent, SNV, indel, UTR, non-truncating coding SNV and the protein-consequence classes (missense, synonymous, nonsense) — sixteen in all. The ten non-splice strata are shown below; the complete grid is Supplemental_Table_S2.

| **Predictor** | **All** | **ClinVar-recorded** | **ClinVar-absent** | **Coding / UTR** | **Missense** | **Synonymous** | **Nonsense** | **UTR** | **SNV** | **Indel** |
|---|---|---|---|---|---|---|---|---|---|---|
| AlphaGenome | 0.146 | 0.181 | 0.102 | 0.077 | 0.056 | 0.063 | -0.059 | -0.036 | 0.155 | 0.053 |
| SpliceAI | 0.177 | 0.207 | 0.130 | 0.120 | 0.102 | 0.061 | 0.051 | 0.030 | 0.177 | — |
| Pangolin | 0.190 | 0.224 | 0.135 | 0.133 | 0.081 | 0.074 | 0.048 | 0.061 | 0.190 | — |
| CADD | 0.391 | 0.445 | 0.312 | 0.427 | 0.376 | 0.023 | 0.115 | 0.029 | 0.391 | — |
| AlphaMissense | 0.438 | 0.472 | 0.369 | 0.438 | 0.438 | — | — | — | 0.438 | — |
| Evo2-7B | 0.358 | 0.403 | 0.283 | 0.362 | 0.287 | 0.050 | 0.129 | 0.035 | 0.358 | — |
| GPN-MSA | 0.363 | 0.415 | 0.286 | 0.378 | 0.364 | 0.047 | 0.196 | 0.043 | 0.363 | — |
| NT-v2-500M | 0.098 | 0.116 | 0.091 | 0.068 | -0.000 | -0.033 | 0.019 | 0.091 | 0.098 | — |
| phyloP-100way | 0.292 | 0.349 | 0.223 | 0.278 | 0.296 | 0.036 | 0.087 | -0.002 | 0.307 | 0.214 |
| phastCons-100way | 0.254 | 0.260 | 0.224 | 0.221 | 0.229 | -0.011 | 0.126 | 0.023 | 0.236 | 0.285 |

| | | | | | | | | | |
|---|---|---|---|---|---|---|---|---|---|
| gnomAD AF (global) | 0.073 | 0.077 | 0.089 | 0.076 | 0.080 | 0.036 | 0.124 | 0.046 | 0.073 | — |
| gnomAD AF (popmax) | 0.075 | 0.078 | 0.072 | 0.074 | 0.076 | 0.035 | 0.113 | 0.017 | 0.075 | — |
| REVEL | 0.395 | 0.419 | 0.337 | 0.397 | 0.397 | — | — | — | 0.395 | — |
| BayesDel (AF) | 0.498 | 0.546 | 0.414 | 0.473 | 0.396 | 0.114 | 0.131 | — | 0.498 | — |
| ClinPred | 0.403 | 0.430 | 0.351 | 0.405 | 0.397 | 0.142 | — | — | 0.403 | — |
| MetaRNN | 0.407 | 0.439 | 0.340 | 0.410 | 0.409 | 0.140 | — | — | 0.407 | — |
| PrimateAI | 0.272 | 0.301 | 0.223 | 0.272 | 0.272 | — | — | — | 0.272 | — |
| VEST4 | 0.466 | 0.508 | 0.384 | 0.468 | 0.394 | 0.119 | 0.139 | — | 0.466 | — |
| ESM-1b | 0.345 | 0.362 | 0.291 | 0.345 | 0.344 | — | — | — | 0.345 | — |

Median ClinVar-recorded minus ClinVar-absent uplift across the twelve predictors: **0.082**. Against the full atlas instead of the absent set the same quantity is 0.033, because the full set contains the recorded variants and dilutes the contrast.

#### S9. Classification, evidence strength, readouts and ensembles

`results/clinical_evidence_v1.tsv`, `results/rna_readout_v1.tsv`, `results/ensemble_logo_v1.tsv` (Supplemental_Table_S7), `results/model_correlation_*_v1.tsv`

**Labels.** 31,835 variants across BARD1, PALB2, RAD51C: GMM posterior ≥ 0.9 / ≤ 0.1 for BARD1 and PALB2 (ambiguous middle dropped), the authors' functional class for RAD51C ('fast depleted' and 'slow depleted' abnormal, 'unchanged' normal, 'enriched' n = 49 excluded).

BARD1: n = 10,733, abnormal = 1,829 (17.0%)

PALB2: n = 11,963, abnormal = 1,657 (13.9%)

RAD51C: n = 9,139, abnormal = 3,094 (33.9%)

AUROC and positive likelihood ratio at 95% specificity, by territory:

The ACMG band column applies the Tavtigian point system at a prior of 0.10 for orientation only; it is not a ClinGen-grade calibration.

**All**

| Predictor | n abnormal | n normal | AUROC | 95% CI | AUPRC | LR+ @95% spec | ACMG band |
|---|---|---|---|---|---|---|---|
| BayesDel (AF) | 2952 | 11454 | 0.908 | 0.888–0.925 | 0.811 | 12.593 | Moderate |
| AlphaMissense | 1703 | 11230 | 0.900 | 0.852–0.934 | 0.707 | 12.601 | Moderate |
| CADD | 3328 | 19632 | 0.899 | 0.872–0.921 | 0.749 | 13.014 | Moderate |
| VEST4 | 2564 | 11293 | 0.896 | 0.879–0.912 | 0.701 | 10.737 | Moderate |
| ESM-1b | 1671 | 11157 | 0.877 | 0.821–0.918 | 0.687 | 10.543 | Moderate |
| ClinPred | 1808 | 11331 | 0.876 | 0.830–0.911 | 0.629 | 10.077 | Moderate |
| MetaRNN | 1735 | 11383 | 0.870 | 0.836–0.897 | 0.545 | 10.100 | Moderate |
| REVEL | 1671 | 11142 | 0.868 | 0.837–0.894 | 0.595 | 9.808 | Moderate |
| Evo2-7B | 3328 | 19632 | 0.849 | 0.776–0.901 | 0.551 | 9.424 | Moderate |
| GPN-MSA | 3328 | 19632 | 0.842 | 0.779–0.890 | 0.576 | 9.160 | Moderate |
| PrimateAI | 1663 | 11124 | 0.795 | 0.755–0.829 | 0.376 | 4.572 | Moderate |
| phastCons- | 6580 | 25255 | 0.732 | 0.683–0.776 | 0.344 | — | not evaluable |

| | | | | | | | |
|---|---|---|---|---|---|---|---|
| 100way | | | | | | | |
| phyloP-100way | 6580 | 25255 | 0.731 | 0.678–0.779 | 0.433 | 6.123 | Moderate |
| Pangolin | 3328 | 19632 | 0.675 | 0.636–0.712 | 0.426 | 5.949 | Moderate |
| NT-v2-500M | 3328 | 19632 | 0.661 | 0.626–0.694 | 0.363 | 5.844 | Moderate |
| SpliceAI | 3328 | 19632 | 0.650 | 0.625–0.674 | 0.390 | 5.509 | Moderate |
| AlphaGenome | 6580 | 25255 | 0.628 | 0.579–0.674 | 0.400 | 4.332 | Moderate |
| gnomAD AF (global) | 680 | 4512 | 0.541 | 0.517–0.564 | 0.130 | 1.484 | below supporting |
| gnomAD AF (popmax) | 680 | 4512 | 0.534 | 0.510–0.558 | 0.130 | — | not evaluable |

#### Coding / UTR

| Predictor | n abnormal | n normal | AUROC | 95% CI | AUPRC | LR+ @95% spec | ACMG band |
|---|---|---|---|---|---|---|---|
| CADD | 2651 | 15711 | 0.916 | 0.890–0.935 | 0.777 | 13.168 | Moderate |
| BayesDel (AF) | 2585 | 11431 | 0.902 | 0.881–0.919 | 0.783 | 12.498 | Moderate |
| AlphaMissense | 1703 | 11230 | 0.900 | 0.852–0.934 | 0.707 | 12.601 | Moderate |
| VEST4 | 2551 | 11272 | 0.900 | 0.888–0.910 | 0.701 | 10.774 | Moderate |
| ESM-1b | 1671 | 11157 | 0.877 | 0.821–0.918 | 0.687 | 10.543 | Moderate |
| ClinPred | 1801 | 11331 | 0.877 | 0.830–0.912 | 0.629 | 10.077 | Moderate |
| MetaRNN | 1724 | 11372 | 0.872 | 0.837–0.900 | 0.545 | 10.115 | Moderate |
| REVEL | 1671 | 11142 | 0.868 | 0.837–0.894 | 0.595 | 9.808 | Moderate |
| Evo2-7B | 2651 | 15711 | 0.842 | 0.769–0.894 | 0.530 | 8.656 | Moderate |
| GPN-MSA | 2651 | 15711 | 0.830 | 0.756–0.884 | 0.535 | 8.130 | Moderate |
| PrimateAI | 1663 | 11124 | 0.795 | 0.755–0.829 | 0.376 | 4.572 | Moderate |
| phyloP-100way | 5683 | 20400 | 0.705 | 0.638–0.765 | 0.426 | 5.016 | Moderate |
| phastCons-100way | 5683 | 20400 | 0.695 | 0.621–0.761 | 0.338 | — | not evaluable |
| NT-v2-500M | 2651 | 15711 | 0.643 | 0.588–0.694 | 0.393 | 6.057 | Moderate |
| Pangolin | 2651 | 15711 | 0.617 | 0.567–0.664 | 0.262 | 3.823 | Supporting |
| SpliceAI | 2651 | 15711 | 0.586 | 0.553–0.618 | 0.214 | 2.384 | Supporting |
| AlphaGenome | 5683 | 20400 | 0.561 | 0.520–0.602 | 0.265 | 2.128 | Supporting |
| gnomAD AF (popmax) | 541 | 3594 | 0.541 | 0.515–0.568 | 0.127 | — | not evaluable |
| gnomAD AF (global) | 541 | 3594 | 0.538 | 0.510–0.565 | 0.140 | 1.098 | below supporting |

#### Splice region

| Predictor | n abnormal | n normal | AUROC | 95% CI | AUPRC | LR+ @95% spec | ACMG band |
|---|---|---|---|---|---|---|---|
| AlphaGenome | 897 | 4855 | 0.963 | 0.949–0.974 | 0.925 | 17.790 | Moderate |
| Pangolin | 677 | 3921 | 0.960 | 0.934–0.976 | 0.906 | 17.287 | Moderate |
| SpliceAI | 677 | 3921 | 0.957 | 0.930–0.974 | 0.898 | 16.636 | Moderate |
| Evo2-7B | 677 | 3921 | 0.899 | 0.777–0.958 | 0.821 | 14.735 | Moderate |
| GPN-MSA | 677 | 3921 | 0.879 | 0.861–0.895 | 0.741 | 12.909 | Moderate |

| | | | | | | | |
|---|---|---|---|---|---|---|---|
| CADD | 677 | 3921 | 0.875 | 0.809–0.921 | 0.793 | 13.331 | Moderate |
| phastCons-100way | 897 | 4855 | 0.842 | 0.809–0.870 | 0.602 | — | not evaluable |
| phyloP-100way | 897 | 4855 | 0.838 | 0.812–0.861 | 0.684 | 12.124 | Moderate |
| NT-v2-500M | 677 | 3921 | 0.735 | 0.671–0.790 | 0.471 | 5.433 | Moderate |
| gnomAD AF (global) | 139 | 918 | 0.564 | 0.510–0.617 | 0.226 | — | not evaluable |
| gnomAD AF (popmax) | 139 | 918 | 0.512 | 0.459–0.564 | 0.187 | — | not evaluable |

#### Splice 3–10 bp

| Predictor | n abnormal | n normal | AUROC | 95% CI | AUPRC | LR+ @95% spec | ACMG band |
|---|---|---|---|---|---|---|---|
| AlphaGenome | 353 | 1294 | 0.921 | 0.882–0.948 | 0.863 | 15.927 | Moderate |
| Pangolin | 248 | 1048 | 0.913 | 0.849–0.951 | 0.798 | 13.227 | Moderate |
| SpliceAI | 248 | 1048 | 0.902 | 0.829–0.945 | 0.804 | 11.574 | Moderate |
| Evo2-7B | 248 | 1048 | 0.803 | 0.694–0.880 | 0.596 | 9.309 | Moderate |
| GPN-MSA | 248 | 1048 | 0.783 | 0.746–0.817 | 0.497 | 6.516 | Moderate |
| CADD | 248 | 1048 | 0.758 | 0.708–0.802 | 0.418 | 4.724 | Moderate |
| phastCons-100way | 353 | 1294 | 0.740 | 0.673–0.797 | 0.379 | — | not evaluable |
| phyloP-100way | 353 | 1294 | 0.734 | 0.694–0.770 | 0.471 | 7.456 | Moderate |
| NT-v2-500M | 248 | 1048 | 0.682 | 0.568–0.778 | 0.334 | 4.654 | Moderate |
| gnomAD AF (global) | 51 | 213 | 0.538 | 0.447–0.627 | 0.287 | — | not evaluable |
| gnomAD AF (popmax) | 51 | 213 | 0.525 | 0.434–0.613 | 0.279 | — | not evaluable |

'Not evaluable' means the score is too coarsely quantised for any threshold to reach 95% specificity — it is not a statement that the predictor is weak.

**Cross-readout.** RNA-level scores are published for 19,553 coding/UTR variants across BARD1, BRCA1, PALB2, VHL; the readout does not cover splice-region variants. The two readouts of the same assay agree at $\rho = 0.251$ (0.124–0.369), $I^2 = 99\%$.

RNA readout reliability (BRCA1, n = 2,743, from replicates): 0.611, ceiling 0.782.

| Predictor | vs fitness | vs RNA | Drop |
|---|---|---|---|
| VEST4 | 0.476 | 0.176 | 0.299 |
| BayesDel (AF) | 0.473 | 0.194 | 0.279 |
| AlphaMissense | 0.424 | 0.095 | 0.329 |
| CADD | 0.412 | 0.165 | 0.247 |
| MetaRNN | 0.408 | 0.102 | 0.307 |
| REVEL | 0.391 | 0.103 | 0.287 |
| ClinPred | 0.383 | 0.100 | 0.283 |
| Evo2-7B | 0.375 | 0.166 | 0.209 |
| GPN-MSA | 0.361 | 0.114 | 0.247 |
| phyloP-100way | 0.303 | 0.087 | 0.216 |

| | | | |
|---|---|---|---|
| ESM-1b | 0.294 | 0.044 | 0.250 |
| PrimateAI | 0.238 | 0.035 | 0.203 |
| phastCons-100way | 0.211 | 0.049 | 0.163 |
| SpliceAI | 0.141 | 0.109 | 0.032 |
| Pangolin | 0.133 | 0.163 | -0.030 |
| AlphaGenome | 0.087 | 0.100 | -0.013 |
| NT-v2-500M | 0.054 | 0.042 | 0.013 |
| gnomAD AF (popmax) | 0.051 | -0.016 | 0.067 |
| gnomAD AF (global) | 0.044 | -0.014 | 0.058 |

**Selection strategies, leave-one-gene-out.** Median over the seven held-out genes; for each fold the strategy is fixed on the other six.

| Stratum | CADD everywhere | Best model per territory | Rank-avg (broad panel) | Rank-avg (splice panel) | Oracle single model |
|---|---|---|---|---|---|
| All | 0.384 | 0.384 | 0.412 | 0.184 | 0.407 |
| Coding / UTR | 0.440 | 0.440 | 0.428 | 0.105 | 0.449 |
| Splice region | 0.360 | 0.432 | 0.403 | 0.473 | 0.467 |
| Splice ±1–2 | 0.127 | -0.015 | 0.144 | 0.185 | 0.370 |
| Splice 3–10 bp | 0.257 | 0.448 | 0.319 | 0.466 | 0.456 |
| Splice 11–50 bp | 0.023 | 0.085 | 0.076 | 0.116 | 0.188 |

**Complementarity.** Between-model Spearman correlation in coding/UTR. The generalists are near-copies of one another and the splice-aware models form a second, largely independent cluster.

| Predictor | CADD | Evo2-7B | GPN-MSA | phyloP-100way | AlphaGenome | SpliceAI | Pangolin | phastCons-100way | NT-v2-500M |
|---|---|---|---|---|---|---|---|---|---|
| CADD | 1.00 | 0.73 | 0.82 | 0.81 | 0.16 | 0.21 | 0.21 | 0.68 | 0.13 |
| Evo2-7B | 0.73 | 1.00 | 0.72 | 0.60 | 0.14 | 0.17 | 0.16 | 0.52 | 0.16 |
| GPN-MSA | 0.82 | 0.72 | 1.00 | 0.79 | 0.17 | 0.21 | 0.16 | 0.74 | 0.10 |
| phyloP-100way | 0.81 | 0.60 | 0.79 | 1.00 | 0.15 | 0.18 | 0.14 | 0.80 | 0.06 |
| AlphaGenome | 0.16 | 0.14 | 0.17 | 0.15 | 1.00 | 0.53 | 0.43 | 0.19 | 0.09 |
| SpliceAI | 0.21 | 0.17 | 0.21 | 0.18 | 0.53 | 1.00 | 0.56 | 0.25 | 0.04 |
| Pangolin | 0.21 | 0.16 | 0.16 | 0.14 | 0.43 | 0.56 | 1.00 | 0.17 | 0.07 |
| phastCons-100way | 0.68 | 0.52 | 0.74 | 0.80 | 0.19 | 0.25 | 0.17 | 1.00 | 0.06 |
| NT-v2-500M | 0.13 | 0.16 | 0.10 | 0.06 | 0.09 | 0.04 | 0.07 | 0.06 | 1.00 |

#### S10. Simulation validation of the attenuation correction

`results/attenuation_simulation_v1.tsv` (Supplemental_Table_S4)

150 trials per gene in each cell, over five stratum shapes x six true rho x six reliabilities; between 2 and 7 genes contribute depending on the stratum, so a cell carries 300 to 1,050 trials and the grid 151,200 in total (seed 20260804). True scores resample the observed distribution of one gene within one stratum — per gene, because the seven assays report on incompatible scales, so a pooled marginal would calibrate the injected noise against the scale differences rather than the within-assay spread. Reliability is re-estimated from simulated replicates exactly as the pipeline estimates it, so the whole chain is tested.

| Reliability | Ceiling | Bias, uncorrected | Bias, corrected | RMSE, uncorrected | RMSE, corrected |
|---|---|---|---|---|---|
| 0.100 | 0.277 | -0.298 | 0.007 | 0.309 | 0.515 |
| 0.250 | 0.451 | -0.237 | -0.017 | 0.248 | 0.212 |
| 0.500 | 0.642 | -0.167 | -0.031 | 0.178 | 0.112 |
| 0.650 | 0.731 | -0.131 | -0.029 | 0.143 | 0.073 |
| 0.800 | 0.817 | -0.092 | -0.022 | 0.104 | 0.053 |
| 0.950 | 0.922 | -0.038 | -0.007 | 0.048 | 0.025 |

The correction is close to unbiased at every level, and reduces RMSE while the ceiling exceeds about 0.45; below that, dividing by a small and noisily estimated ceiling amplifies error. Of the strata measured here only the deep intron (ceiling 0.297) falls below that boundary, and it is reported uncorrected throughout.

Bias by stratum shape, which tests whether the compressed distribution at splice +-1-2 breaks the correction (it does not):

| Stratum | Bias, uncorrected | Bias, corrected |
|---|---|---|
| coding_or_utr | -0.180 | -0.053 |
| splice_11_50 | -0.173 | -0.014 |
| splice_1_2 | -0.138 | 0.006 |
| splice_3_10 | -0.182 | -0.039 |
| splice_deep | -0.129 | 0.018 |

#### S11. AlphaGenome definition sweep

This section carries the full definition sweep, demoted from the main text: the sixteen window × aggregation combinations, the client-version control, and the endpoint comparison between the two published score columns. The figure is Supplemental_Fig_S16 and the grid is Supplemental_Table_S8.

`results/definition_sweep_performance_v1.tsv` (Supplemental_Table_S8), `results/definition_sweep_concordance_v1.tsv`

2,722 variants, sampled stratified by gene and region with splice strata oversampled, scored under four API-supported window widths. Each response carries both raw and quantile scores for all three splicing output types, so four aggregations were derived per call and only the window cost extra calls: 10,888 calls, cached per variant x window, no errors.

| stratum | width | max_abs_raw | max_quantile | merged_quantile | merged_raw |
|---|---|---|---|---|---|
| coding_or_utr | 16384 | 0.067 | 0.101 | 0.069 | 0.098 |
| coding_or_utr | 131072 | 0.063 | 0.070 | 0.111 | 0.083 |
| coding_or_utr | 524288 | 0.070 | 0.059 | 0.094 | 0.082 |
| coding_or_utr | 1048576 | 0.081 | 0.077 | 0.069 | 0.103 |
| splice_11_50 | 16384 | 0.228 | 0.214 | 0.188 | 0.220 |
| splice_11_50 | 131072 | 0.222 | 0.230 | 0.188 | 0.227 |
| splice_11_50 | 524288 | 0.223 | 0.232 | 0.168 | 0.221 |
| splice_11_50 | 1048576 | 0.218 | 0.227 | 0.136 | 0.209 |
| splice_1_2 | 16384 | 0.063 | 0.078 | 0.173 | 0.097 |

| | | | | | |
|---|---|---|---|---|---|
| splice_1_2 | 131072 | 0.123 | 0.120 | 0.087 | 0.137 |
| splice_1_2 | 524288 | 0.097 | 0.156 | 0.098 | 0.120 |
| splice_1_2 | 1048576 | 0.093 | 0.144 | 0.097 | 0.117 |
| splice_3_10 | 16384 | 0.464 | 0.463 | 0.476 | 0.478 |
| splice_3_10 | 131072 | 0.466 | 0.465 | 0.461 | 0.478 |
| splice_3_10 | 524288 | 0.463 | 0.459 | 0.455 | 0.470 |
| splice_3_10 | 1048576 | 0.458 | 0.457 | 0.454 | 0.468 |
| splice_deep | 16384 | -0.006 | 0.022 | 0.041 | -0.011 |
| splice_deep | 131072 | 0.030 | 0.029 | 0.045 | 0.031 |
| splice_deep | 524288 | -0.026 | -0.024 | 0.060 | -0.014 |
| splice_deep | 1048576 | -0.004 | 0.005 | 0.056 | 0.014 |

**Client version.** Replicating the legacy definition (max|raw|, widest window) under the current client reproduces the legacy column at rho = 0.995 (n = 1,295), so the client version contributes nothing to the rho = 0.672 disagreement; it is entirely definitional.

**Score agreement between the two published columns**: 0.672 overall; coding_or_utr 0.615, splice_11_50 0.706, splice_1_2 0.383, splice_3_10 0.839, splice_deep 0.952.

**Performance of the two definitions on the same variants** — the quantity that actually matters:

| Stratum | Atlas definition | Legacy definition | Difference |
|---|---|---|---|
| all | 0.181 | 0.190 | 0.009 |
| coding_or_utr | 0.101 | 0.118 | 0.017 |
| splice_region | 0.583 | 0.592 | 0.009 |
| splice_1_2 | 0.145 | 0.118 | -0.027 |
| splice_3_10 | 0.463 | 0.480 | 0.017 |
| splice_11_50 | 0.088 | 0.134 | 0.047 |

The largest difference in any stratum is 0.047. A score-level concordance of 0.672 changed no conclusion this benchmark draws, so concordance gates should be evaluated on the benchmark's endpoints rather than on the correlation between score columns.

#### S12. Repository and reproduction

One-command reproduction from a clean clone: `make fetch && make test`. The pipeline is covered by 202 collected guardrail tests, 43 of them covering the analyses introduced here. A bare clone runs 130 and skips 72 for want of data; a clean extract of the release archive runs 178 and skips 24 with no network access. Twenty-three of those skips are the same on any machine: they need a git checkout, the author's submission directory, or the hygiene denylist, which is deliberately not archived. The twenty-fourth is not -- it verifies the mapped reference bases against the Ensembl REST API, so it runs only where that service is reachable, and the same extract then passes one more. Frozen data are immutable under `data/{raw,frozen}/` with SHA-256 manifests; every figure, table and number in the manuscript is produced by a committed script.

Analysis modules added for this work: `atlas.assay_aux`, `atlas.consequence`, `atlas.reliability`, `atlas.robustness`, `atlas.evaluate_ext`, `atlas.clinical_evidence`, `atlas.ensemble`, `atlas.matched_realisation`, `atlas.supplement`; figures from `figures/hardening_figures.py`.

#### S13. Machine-readable source files

Two of the Fourteen Supplemental Tables render a repository file directly. `Supplemental_Table_S11` is the per-predictor training provenance — training data, whether it carries clinical labels, whether any overlap with MAVE or SGE measurements is documented, and a documentation URL for every row. `Supplemental_Table_S12` collects the summary statistics quoted in the manuscript together with the output file each is computed from, so those figures can be checked without opening the deposit.

Twenty-six analysis-level source files carry the full numeric output behind the Note sections named below, at a granularity too fine to typeset — every model × stratum × gene cell rather than the summarised rows shown in the Supplemental Tables (all TSV except the Markdown mapping audit). They are deliberately not numbered as Supplemental Tables: they are part of the Zenodo data deposit (concept DOI 10.5281/zenodo.21828448), where they sit under `results/`; `results/` is not tracked in the GitHub repository and these files are not part of the submission package. Nothing in the manuscript depends on a reader opening them.

`mavedb_survey_v1.tsv` — every published MaveDB score set, with the replicate and error columns found in its score table, the replicate subset (if any) whose mean reproduces the deposited score, and the resulting classification (Note S14; Methods)

`mavedb_survey_audit_v1.tsv` — the sampled deposits whose full score-table header was inspected to check the column-name classification in both directions, with every column seen (Methods)

`mavedb_ceilings_v1.tsv` — per-deposit reliability and attenuation ceiling for every surveyed deposit that permits one, with the estimator used, the reference-class stratum and the status classification (Fig. 6; Methods)

`mavedb_estimator_comparison_v1.tsv` — deposits publishing both reconcilable replicates and an error column, with the reliability each estimator gives and the difference between them

`protease_agreement_v1.tsv` — trypsin-versus-chymotrypsin agreement per designed protein, the Spearman-Brown reliability it implies, and the ceiling the deposited fitting interval implies

`mavedb_metadata_v1.tsv` — deposit metadata used for the correlates in Supplemental_Fig_S17: publication and creation dates, variant counts and target genes

`matched_realisation_v1.tsv` — coding/UTR versus splice ±1-2 on the nine broad-scope predictors that carry an estimate in both strata, observed and attenuation-corrected, with the ratio between territories under each (Results; Methods)

`reliability_v1.tsv` — per-gene replicate-based measurement reliability and the implied attenuation ceiling for each stratum (source of Note S5 and Table S3)

`reliability_corroboration_v1.tsv` — independent corroboration of the reliability estimates: agreement ρ and implied ceiling from held-out assay comparisons, per gene and stratum (Note S5)

`power_v1.tsv` — minimum Spearman ρ detectable at 80% power, per gene and pooled, for every model × stratum cell (Note S6)

`logo_v1.tsv` — leave-one-gene-out pooled ρ for every model × stratum cell, with the held-out gene identified (Note S7) `rna_readout_v1.tsv` — per-gene correlations of each predictor with the RNA-level readout and with fitness, the cross-readout comparison of Note S9

`definition_sweep_concordance_v1.tsv` — pairwise score concordance between AlphaGenome definitions at each API window width (Note S11)

`clinical_evidence_v1.tsv` — AUROC, AUPRC and positive likelihood ratio at 95% specificity per model and territory against assay-derived functional labels, with bootstrap intervals (Note S9)

`ensemble_pooled_v1.tsv` — pooled ρ of the five predictor-selection strategies per stratum over all seven genes; Table S7 reports the leave-one-gene-out median counterpart (Note S9)

`model_correlation_all_v1.tsv` — between-model Spearman correlation matrix over all scored variants

`model_correlation_coding_or_utr_v1.tsv` — between-model Spearman correlation matrix over coding/UTR variants (the Note S9 heatmap)

`model_correlation_splice_region_v1.tsv` — between-model Spearman correlation matrix over splice-region variants

`predictor_resources_v1.tsv` — the curated source of truth behind Supplemental_Table_S9: per-predictor resource, version, source, access date, and the licence governing that score column with the URL it was read from (generated by `atlas.predictor_resources`; S9 renders it with the scope, score definition and coverage columns joined on)

`mapping_report_v1.md` — the human-readable mapping audit: per-assay variant counts, Mutalyzer3 versus Ensembl exon-model agreement, VEP cross-validation pass rates and the orientation check behind the frozen matrix (Methods; the machine-readable counterpart is mapping_summary_v1.json)

`estimator_consistency_v1.tsv` — the BRCA1 estimator cross-check behind Table S14: per-stratum reliability and ceiling under the replicate chain and under the SE estimator, with the difference between them

`matched_realisation_loo_v1.tsv` — leave-one-gene-out forms of the matched coding-versus-splice ratio behind Table S13: median coding and splice ±1–2 realisations and their ratio with each ceiling-bearing gene dropped in turn

`mavedb_accounting_v1.tsv` — the MaveDB deposit ledger behind Table S16's accounting: every counting definition from 2,803 published score sets down to the 674 computable ceilings, with the denominator and fraction for each

`mavedb_ceiling_k_sensitivity_v1.tsv` — sensitivity of the MaveDB ceiling distribution to treating a deposited standard deviation as the error of a mean of k replicates, for k = 1, 2, 3 (Table S16)

`head_to_head_pvalues_v1.tsv` — Steiger and gene-cluster bootstrap P values side by side for all eighteen claimed orderings, with the significance verdict under each and whether they agree (Table S17)

`attenuation_simulation_v2.tsv` — the hardened attenuation simulation behind Table S15: every noise scenario x estimator x stratum x reliability target cell with bias and RMSE before and after correction (1,800 rows; the v1 grid is untouched in attenuation_simulation_v1.tsv)

#### S14. What MaveDB deposits permit

`results/mavedb_survey_v1.tsv`

Every published MaveDB score set (2,803) was enumerated through the public API and its score table retrieved, then classified by whether it carries replicate columns whose mean reproduces the deposited score, or a per-variant error estimate. 2,452 of 2,803 (87%) permit a reliability estimate on that test.

| permits | score sets | % of published |
|---|---|---|
| errors | 2430 | 86.700 |
| none | 322 | 11.500 |
| replicates_unverified | 29 | 1.000 |
| replicates | 22 | 0.800 |

The test is looser than the reconciliation the primary analysis applies, so these are upper bounds on what is usable; deposits whose replicate columns do not reproduce the score are listed as `replicates_unverified` and are not counted. The per-deposit table, including the column names matched in each case, is the TSV named above.

#### S15. Likelihood ratios at 95% specificity on two bases

`results/lr_basis_check_v1.tsv`, `results/lr_basis_check_genes_v1.tsv`

The four likelihood ratios quoted in the Results, at the observed threshold this study reports and interpolated to exactly 95% specificity:

| Predictor · territory | Scanned LR+ | Realised spec. | Interp. LR+ (95%) | Band |
|---|---|---|---|---|
| AlphaGenome · Splice region | 17.79 | 95.02–95.06% | 17.56 | Moderate |
| Pangolin · Splice region | 17.29 | 95.00–95.10% | 16.94 | Moderate |
| SpliceAI · Splice region | 16.64 | 95.00–95.10% | 16.64 | Moderate |
| CADD · Coding / UTR | 13.17 | 95.08–95.32% | 13.08 | Moderate |

Positive likelihood ratios in this study are read at the most sensitive observed score threshold that reaches 95% specificity; the companion splice-region study interpolates to exactly 95% specificity on the empirical ROC. For the well-resolved predictors above the two bases differ by at most 0.35 and agree on the evidence band. For coarsely quantised scores (phastCons and the two gnomAD allele-frequency columns) no observed threshold reaches 95% specificity in several territories, because the top score value is itself a tie group holding more than 5% of the normal variants. An interpolated value there would be read on the ROC segment between calling no variant positive and calling that whole tie group positive, which no single score threshold achieves; those cells are therefore reported as not evaluable, and cells where a threshold does reach 95% specificity at the operating point it achieves, not interpolated. The full scanned-and-interpolated comparison for all 69 cells is in the deposited analysis (`docs/lr-basis-check.md`, written by `atlas.lr_basis_check`).

### Supplemental Fig. S1 — AlphaGenome: per-gene correlation with functional score, by stratum

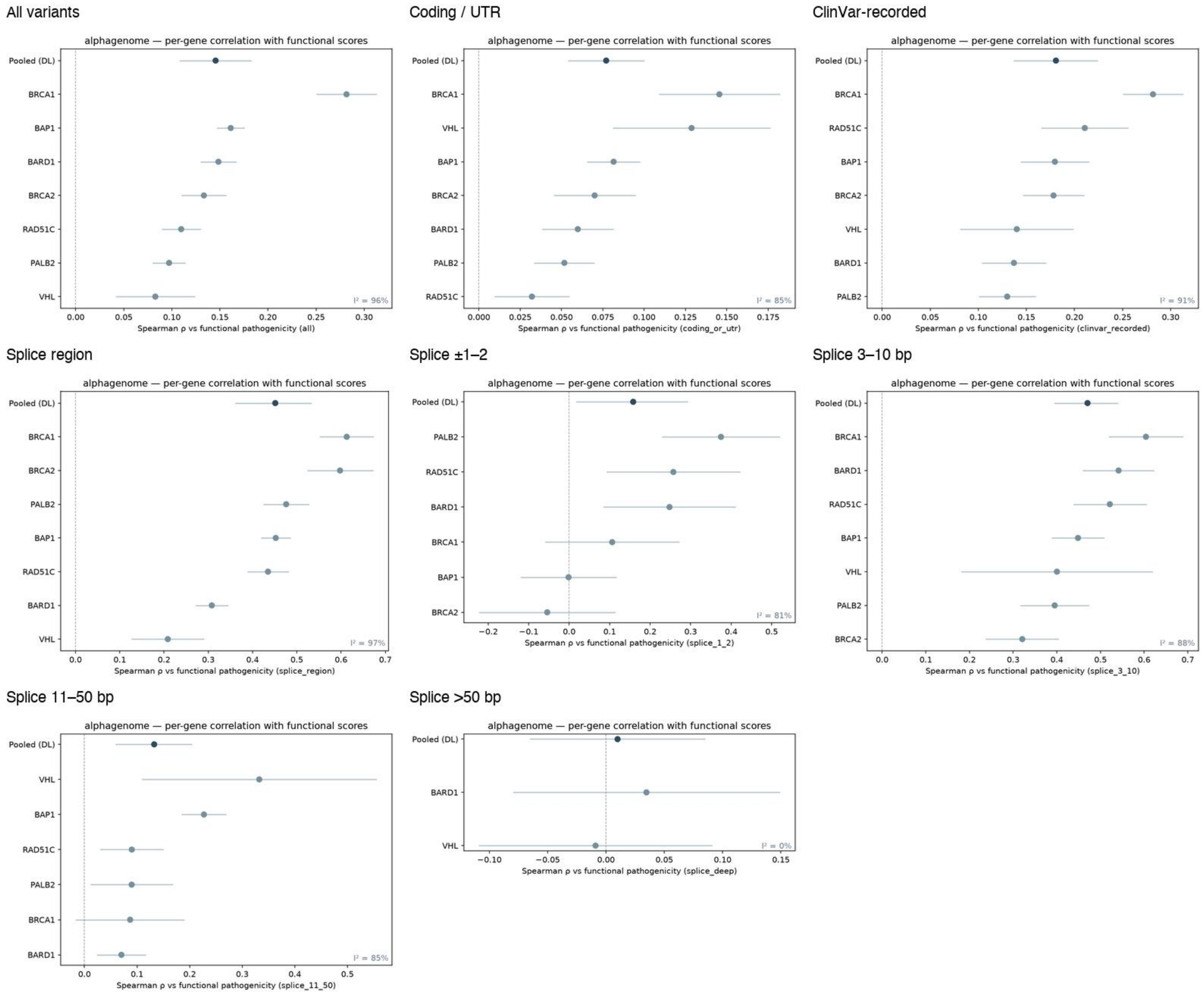

### Supplemental Fig. S2 — SpliceAI: per-gene correlation with functional score, by stratum

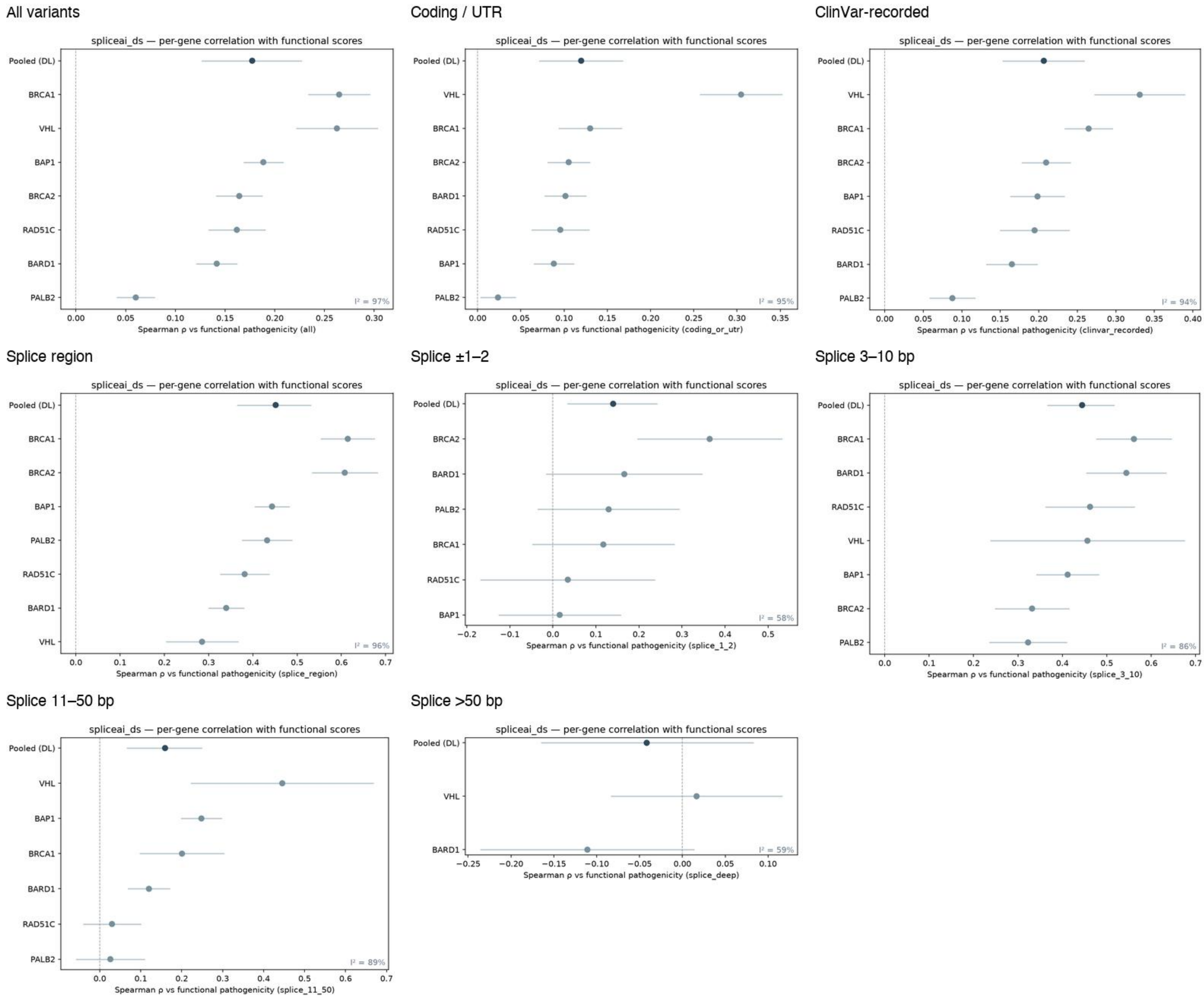

### Supplemental Fig. S3 — Pangolin: per-gene correlation with functional score, by stratum

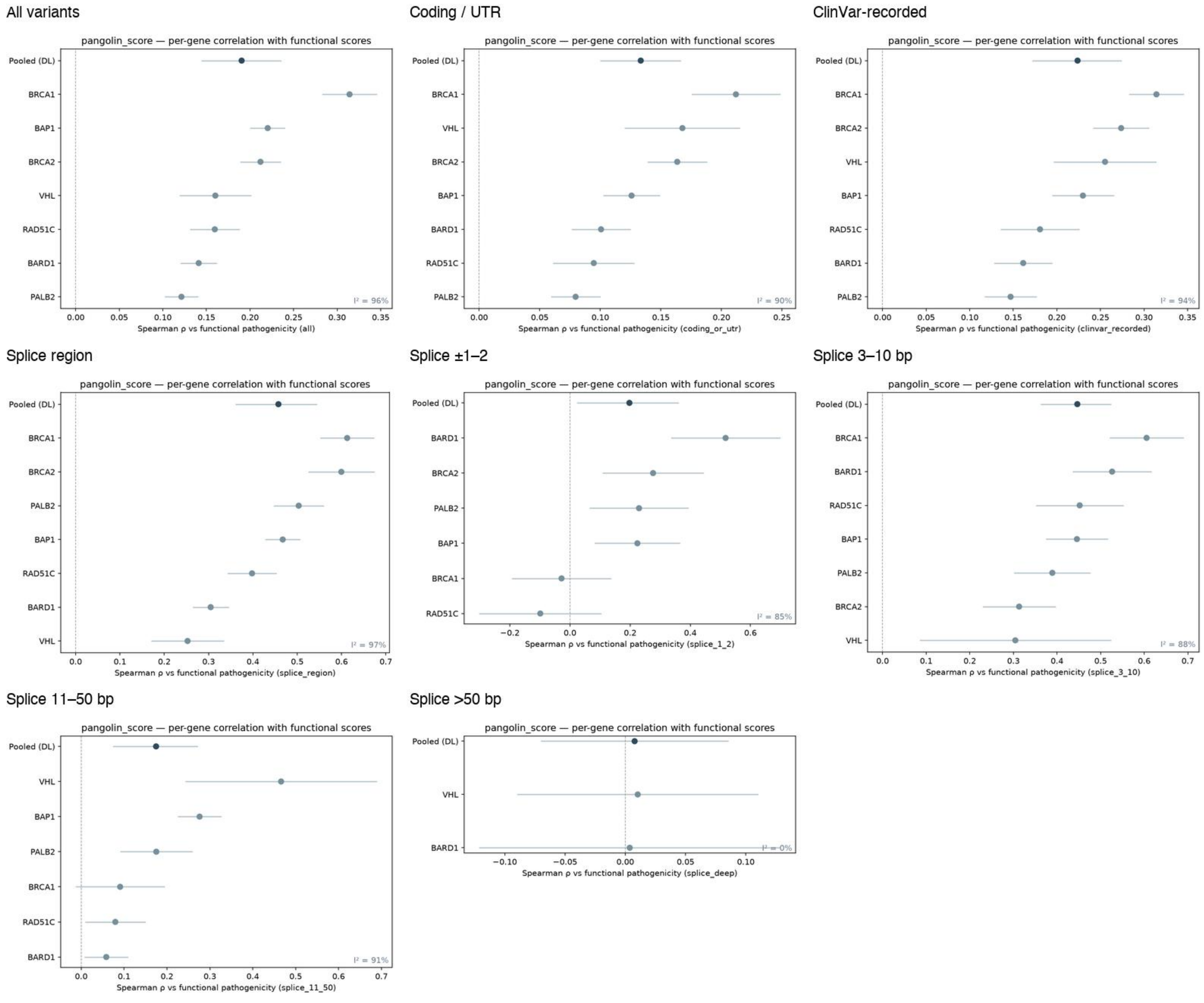

Supplemental Fig. S4 — CADD: per-gene correlation with functional score, by stratum

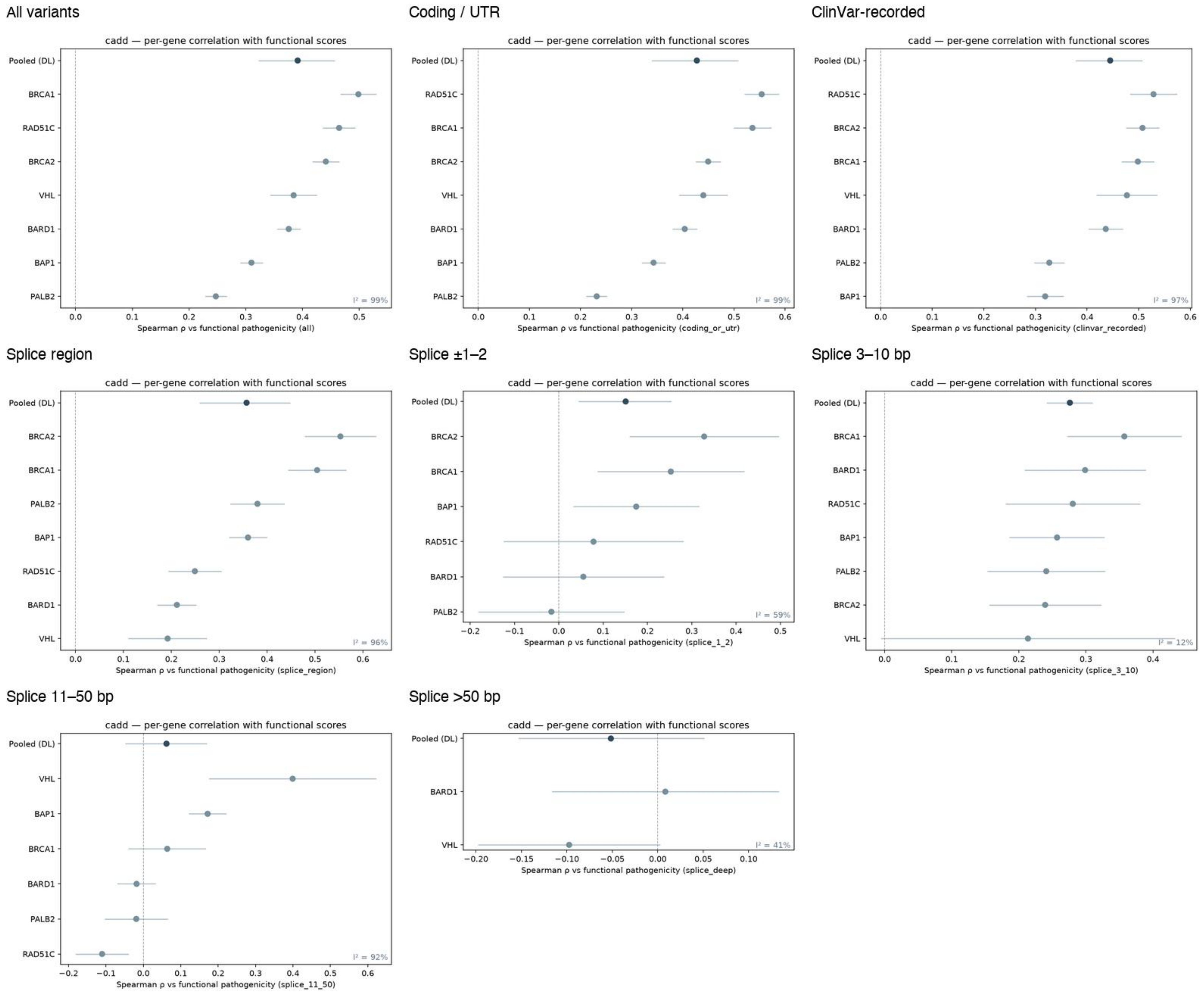

### Supplemental Fig. S5 — AlphaMissense: per-gene correlation with functional score, by stratum

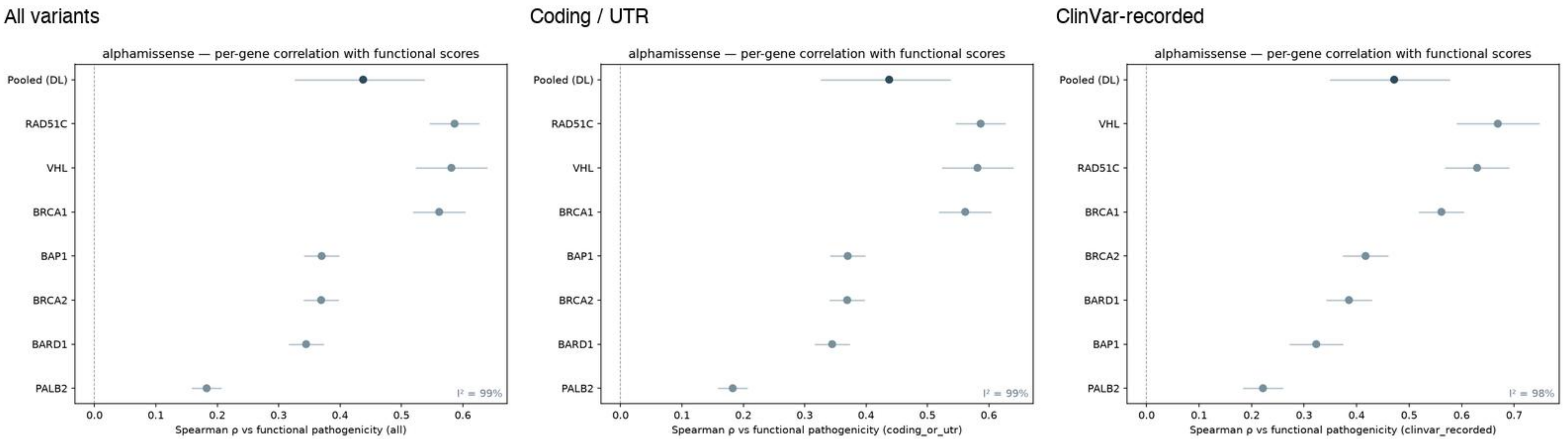

Supplemental Fig. S6 — Evo2-7B: per-gene correlation with functional score, by stratum

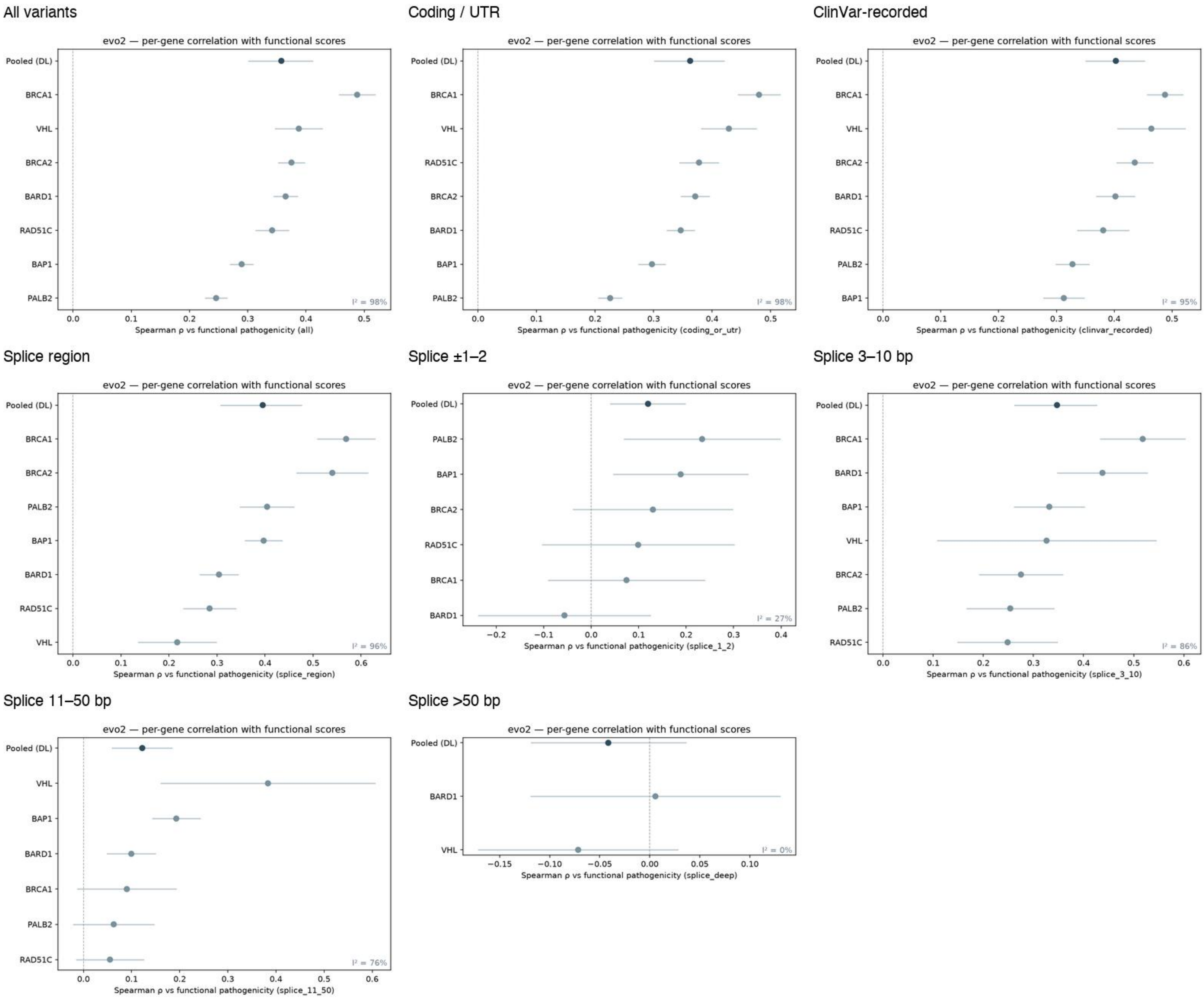

### Supplemental Fig. S7 — GPN-MSA: per-gene correlation with functional score, by stratum

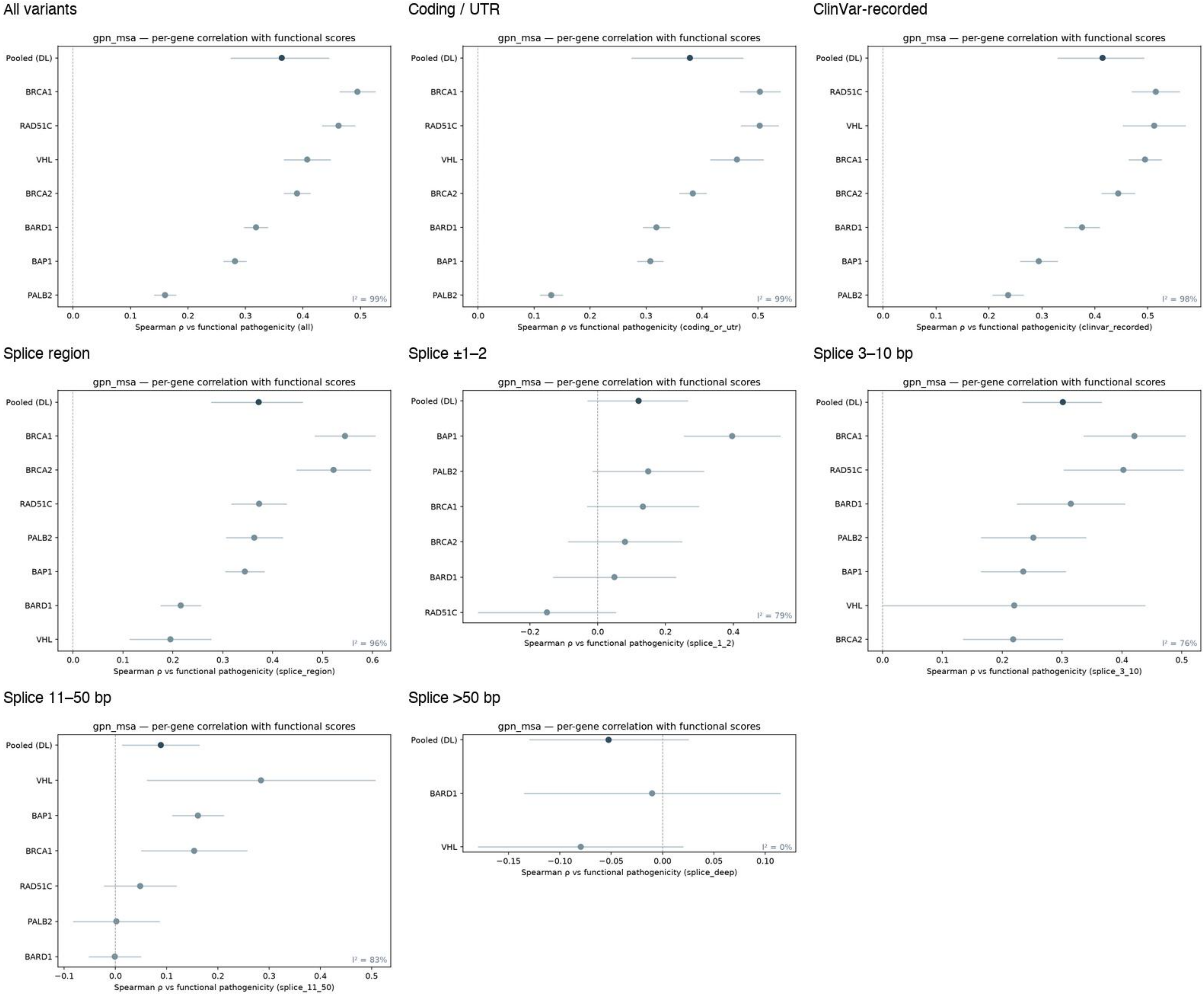

#### Supplemental Fig. S8 — NT-v2-500M: per-gene correlation with functional score, by stratum

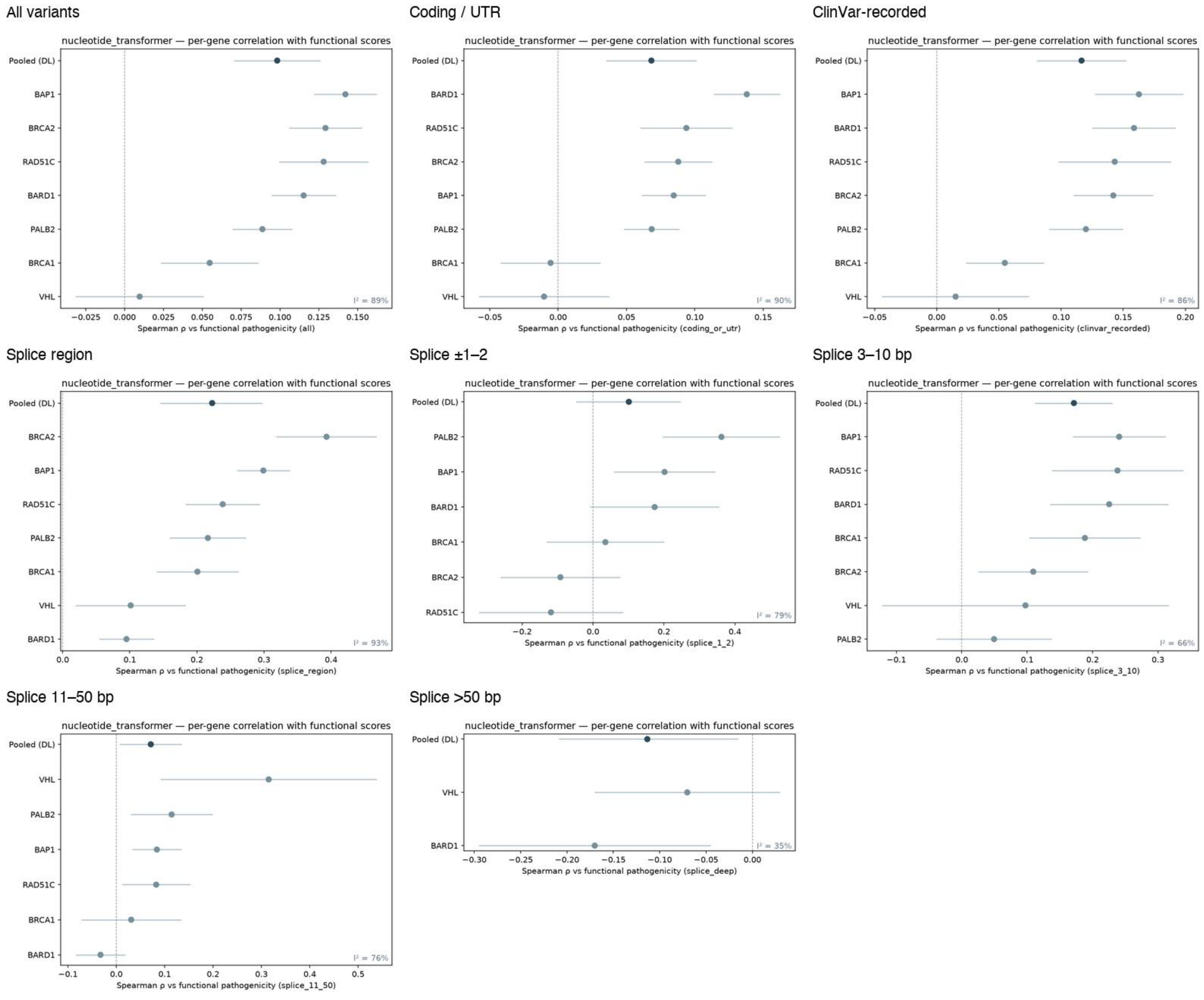

### Supplemental Fig. S9 — phyloP-100way: per-gene correlation with functional score, by stratum

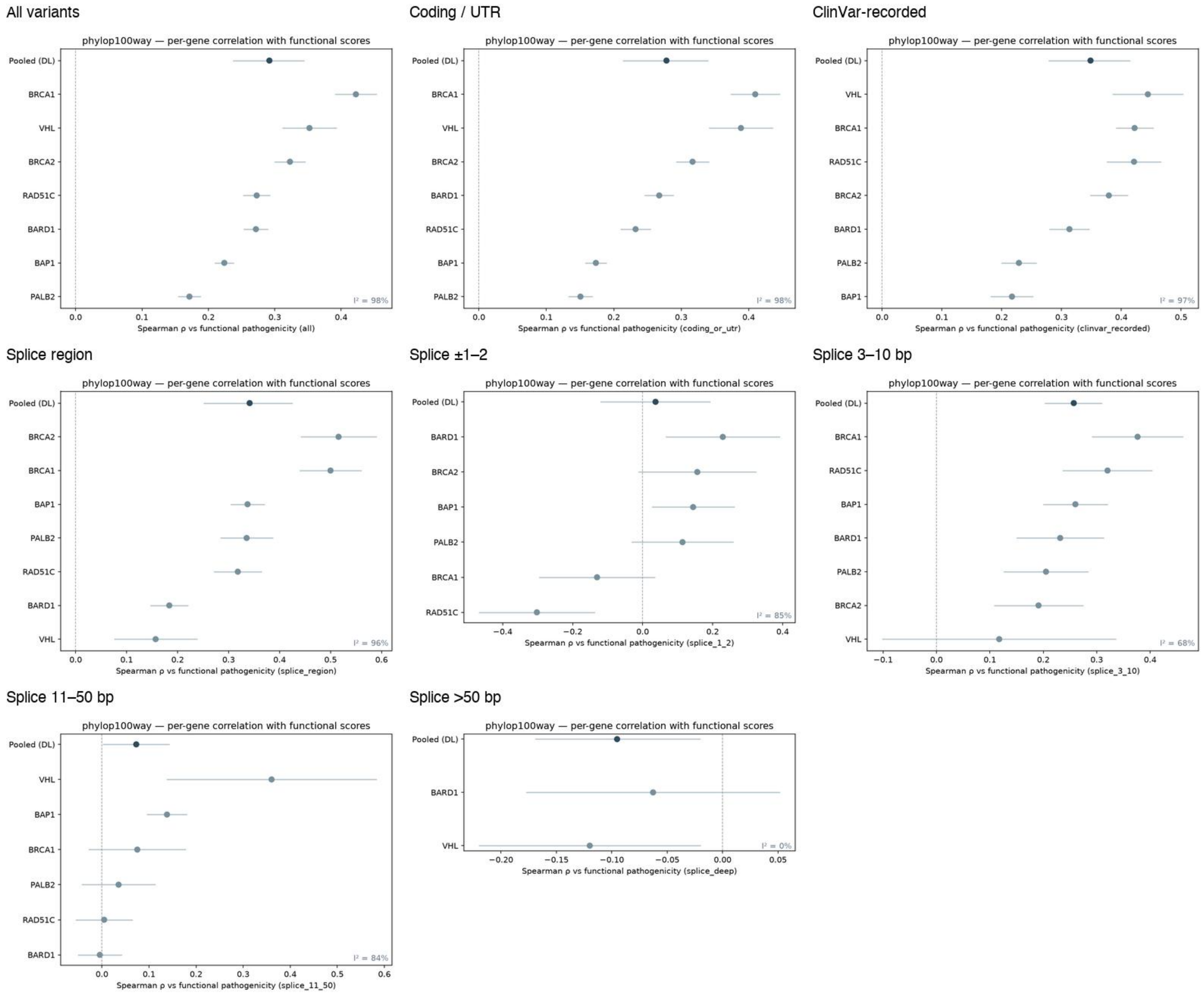

### Supplemental Fig. S10 — phastCons-100way: per-gene correlation with functional score, by stratum

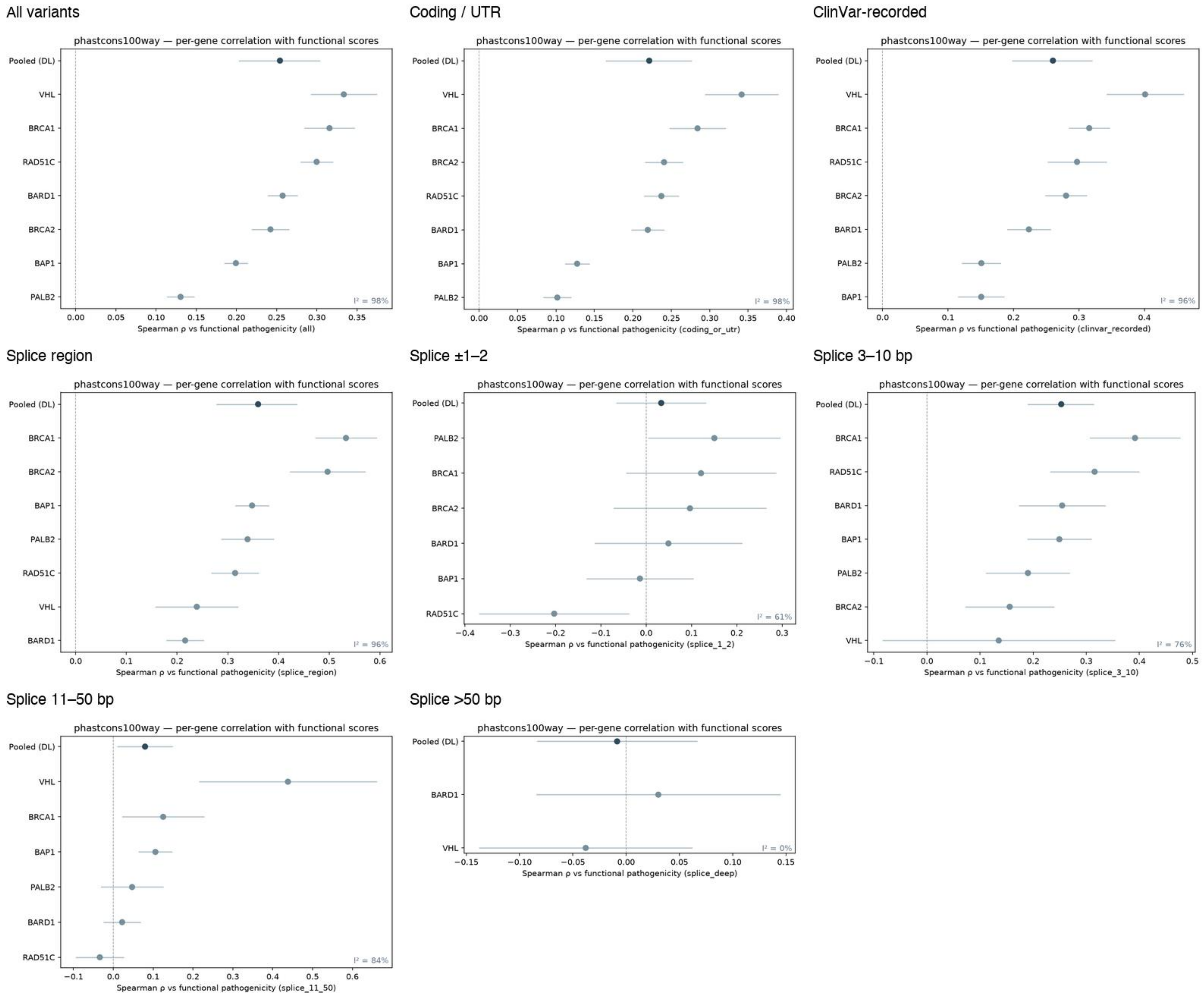

Supplemental Fig. S11 — gnomAD AF (global): per-gene correlation with functional score, by stratum

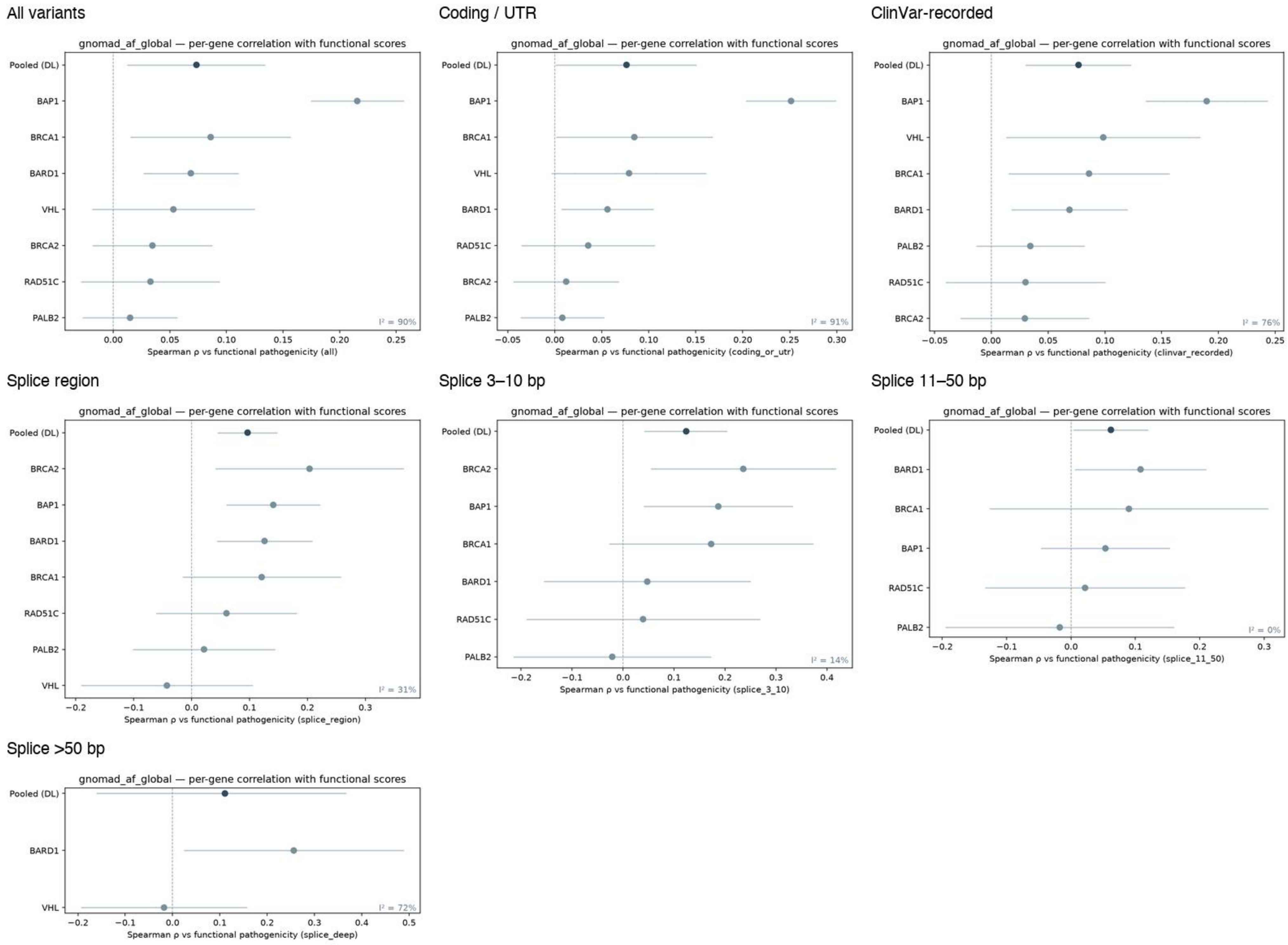

Supplemental Fig. S12 — gnomAD AF (popmax): per-gene correlation with functional score, by stratum

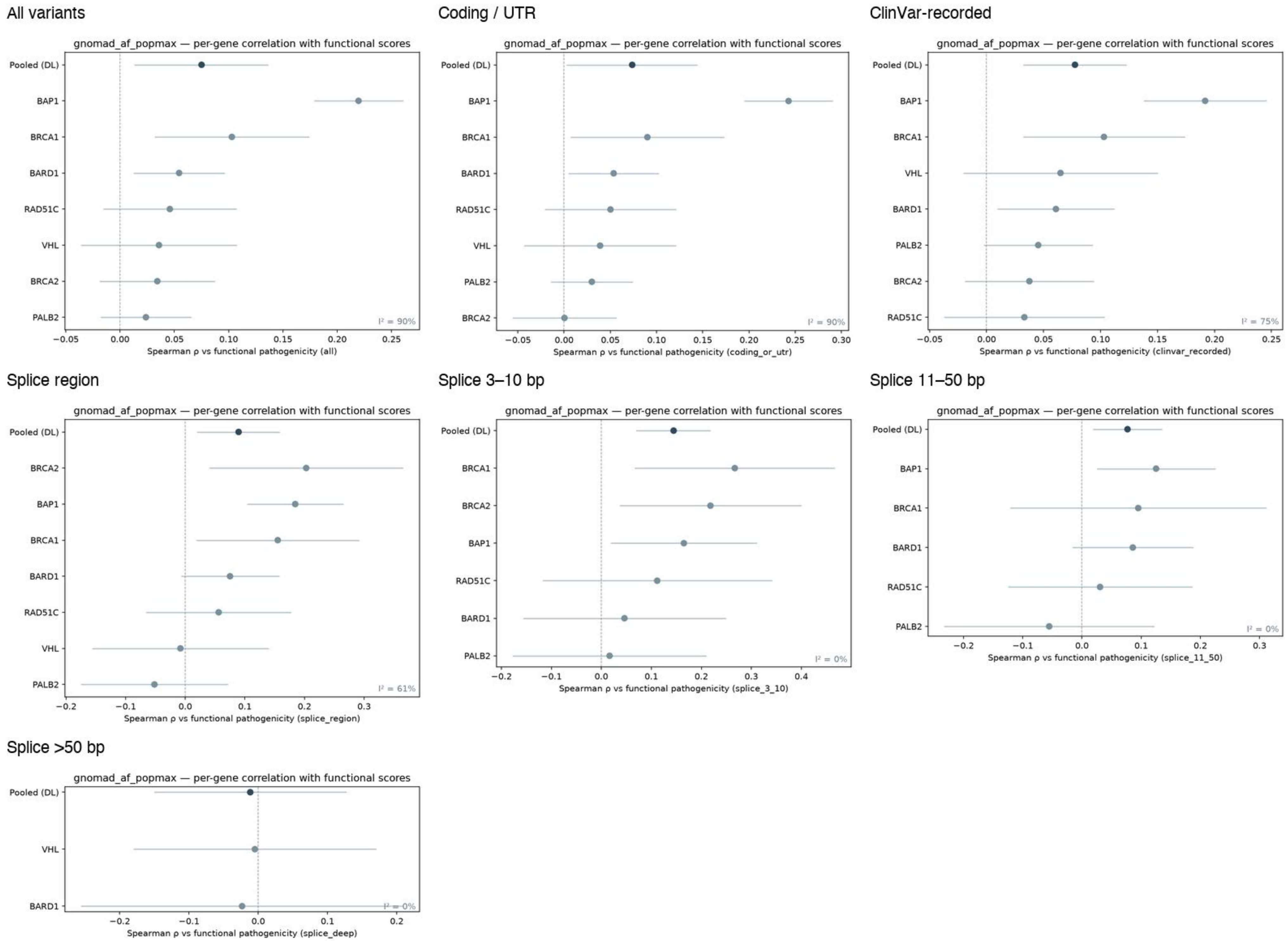

Supplemental Fig. S13 — Concordance with the companion benchmark, per ported score

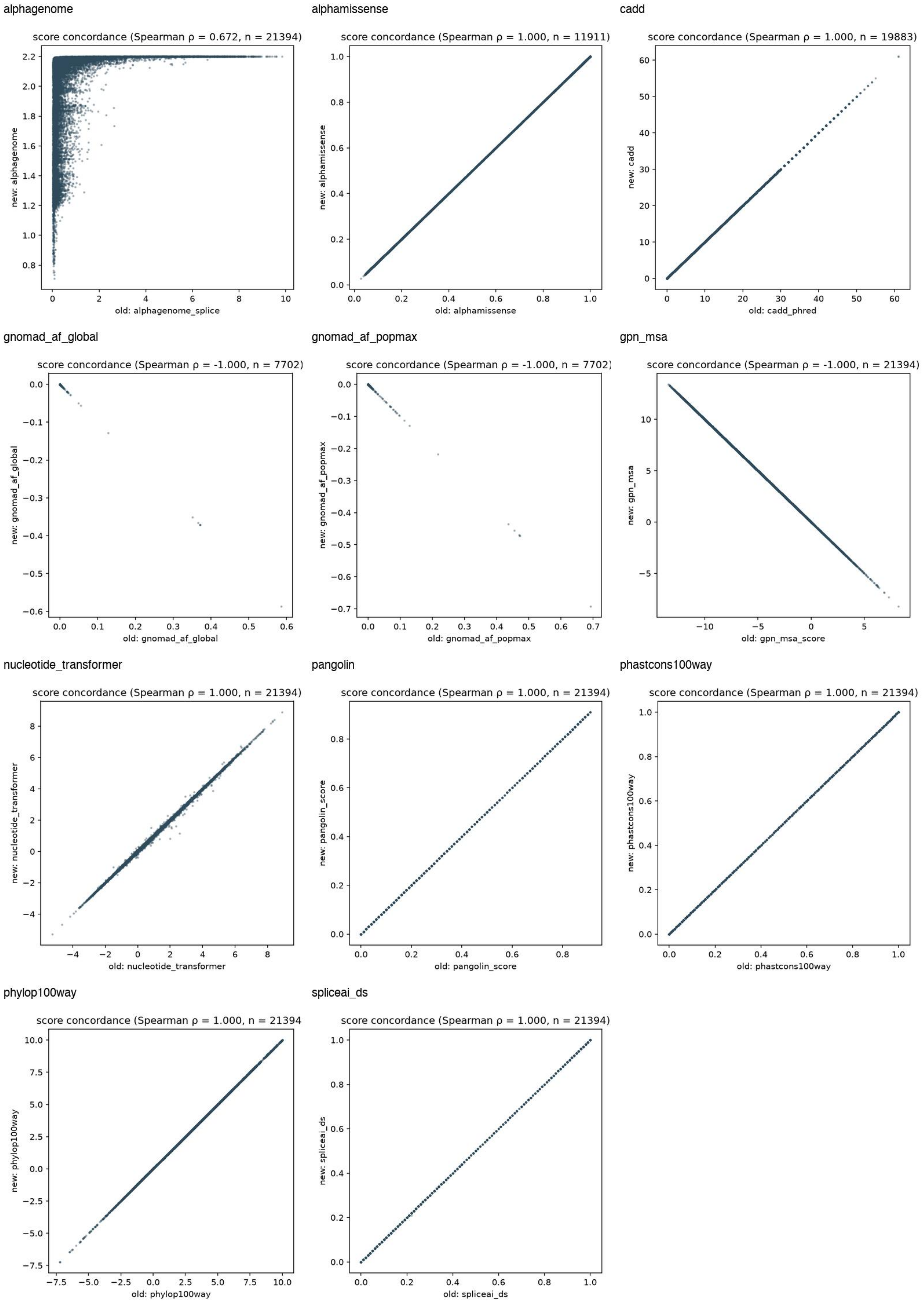

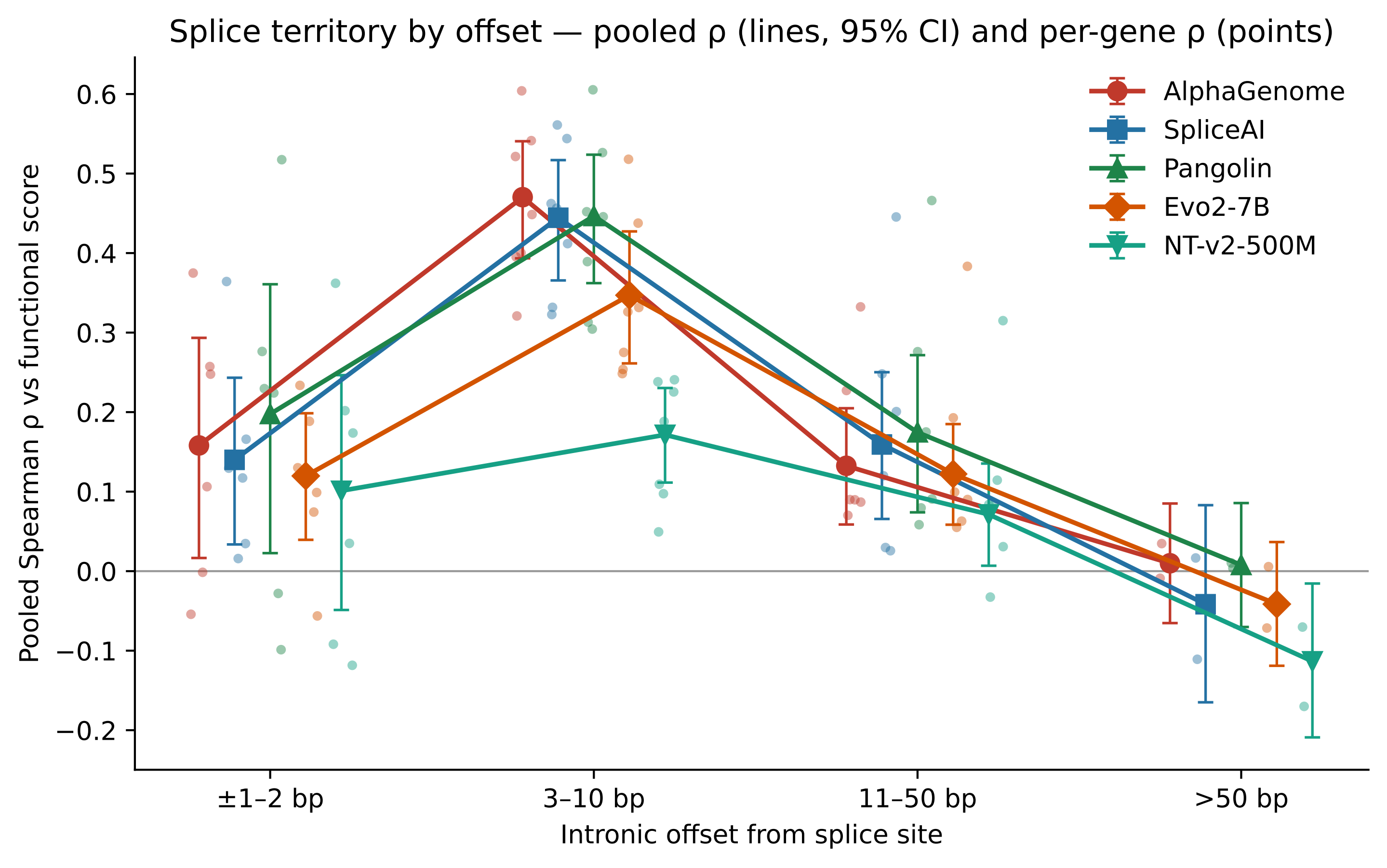
Splice territory by offset — pooled ρ (lines, 95% CI) and per-gene ρ (points)
Pooled Spearman ρ vs functional score
Intronic offset from splice site
±1–2 bp
3–10 bp
11–50 bp
>50 bp
0.6
0.5
0.4
0.3
0.2
0.1
0.0
−0.1
−0.2
AlphaGenome
SpliceAI
Pangolin
Evo2-7B
NT-v2-500M

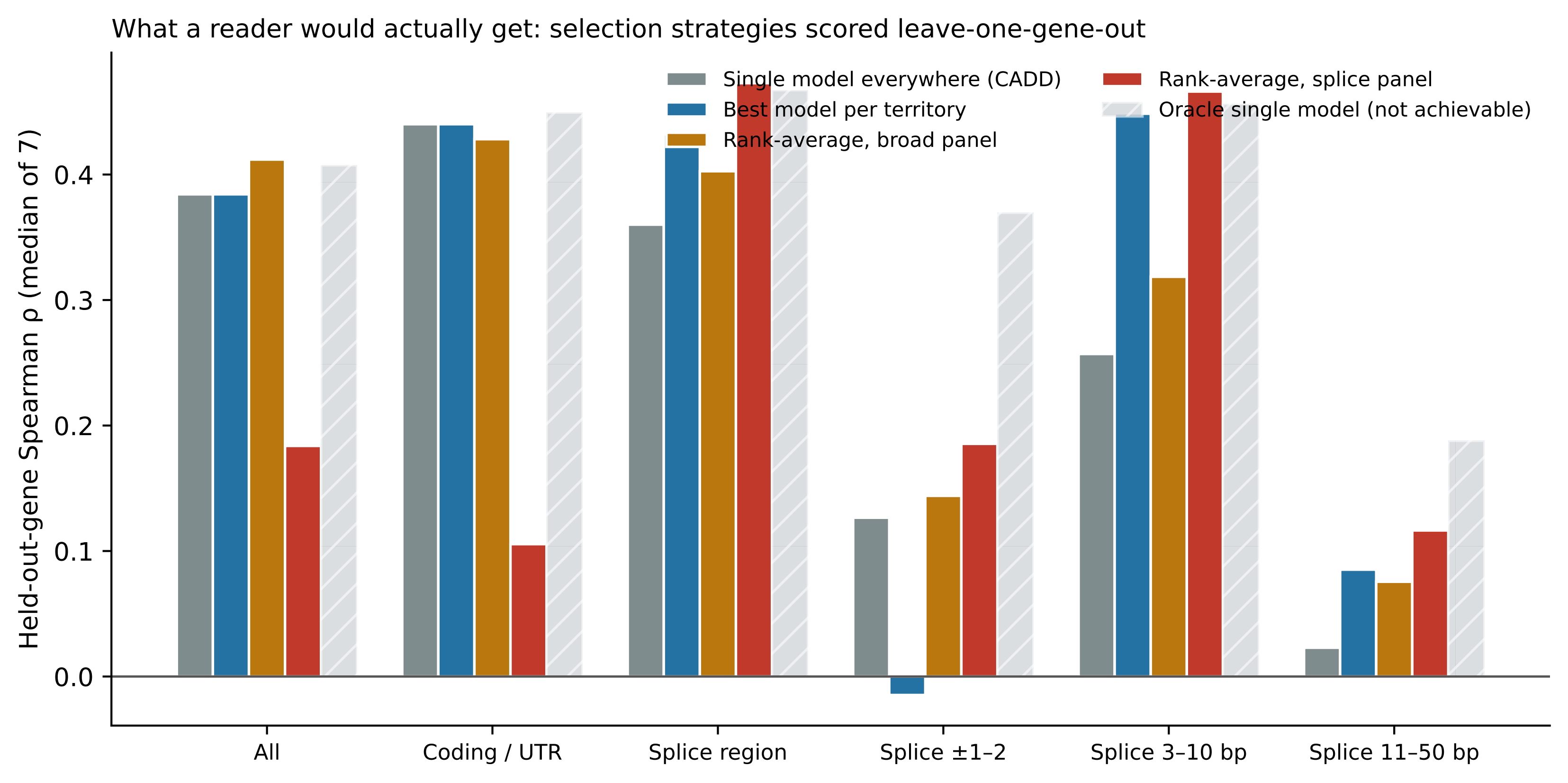
What a reader would actually get: selection strategies scored leave-one-gene-out
Held-out-gene Spearman ρ (median of 7)
0.0
0.1
0.2
0.3
0.4
Single model everywhere (CADD)
Best model per territory
Rank-average, broad panel
Rank-average, splice panel
Oracle single model (not achievable)
All
Coding / UTR
Splice region
Splice ±1–2
Splice 3–10 bp
Splice 11–50 bp

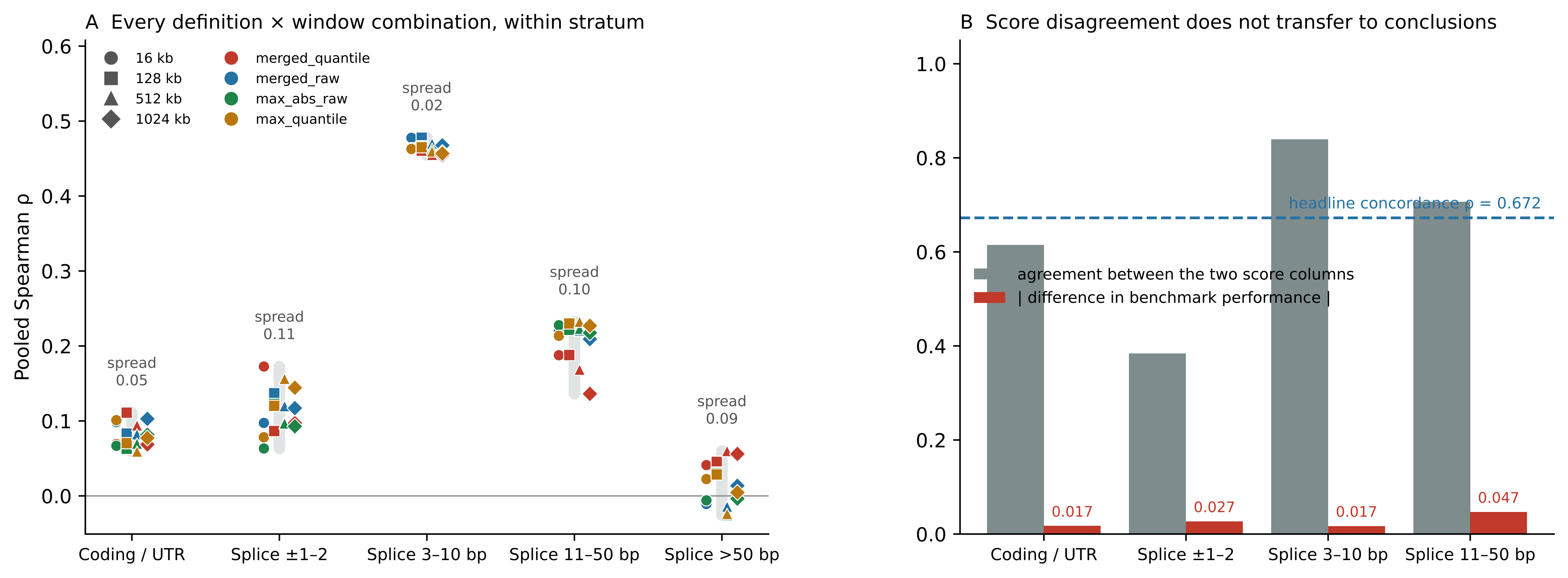

A Every definition × window combination, within stratum
Pooled Spearman ρ
16 kb
128 kb
512 kb
1024 kb
merged_quantile
merged_raw
max_abs_raw
max_quantile
spread 0.05
spread 0.11
spread 0.02
spread 0.10
spread 0.09
Coding / UTR
Splice ±1–2
Splice 3–10 bp
Splice 11–50 bp
Splice >50 bp
B Score disagreement does not transfer to conclusions
headline concordance ρ = 0.672
agreement between the two score columns
| difference in benchmark performance |
0.017
0.027
0.017
0.047
Coding / UTR
Splice ±1–2
Splice 3–10 bp
Splice 11–50 bp

#### Experiment size and deposition year explain almost none of the spread

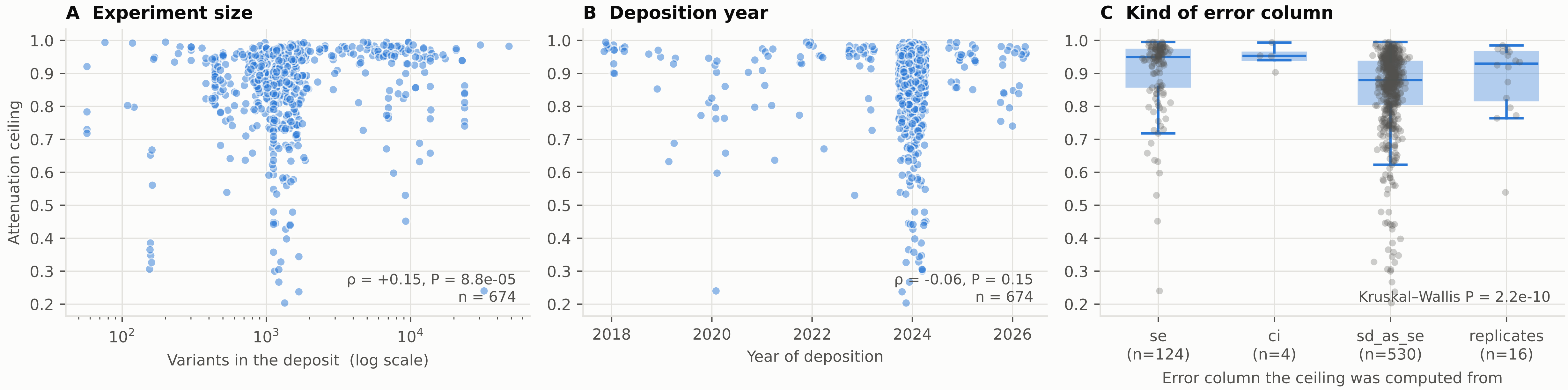


Human deposits with a computable ceiling (n = 674). Deposition years are heavily unbalanced (most deposits are 2023 or later), so B has limited power to detect a trend. The size correlation in A is statistically detectable but weak (it accounts for about 2% of the variance).
